# Disparate demographic impacts of the Roman Colonization and the Migration Period in the Iberian Peninsula

**DOI:** 10.1101/2024.09.23.614606

**Authors:** Pablo Carrión, Iñigo Olalde, Juan Manuel Jiménez-Arenas, Neus Coromina, David Vivó, Josep María Vergès, Ana Costa, Daniel Botella, Macarena Bustamante-Álvarez, Javier Heras-Mora, Ricardo Ortega-Ruiz, Celia Chaves, Maite Iris García-Collado, Juan Antonio Quirós-Castillo, Jordi Roig, José Suárez-Padilla, Ildelfonso Navarro-Luengo, Miguel Ángel Cuadrado, Isidro Aguilera, Jordi Morera, Raúl Catalán, María Luisa Cerdeño, Josep F. Roig-Pérez, Moisés Díaz-García, Paula Chirosa-Cañavate, Tatiana Piza-Ruiz, Elena Vallejo-Casas, Sergio Vidal-Álvarez, Josep Burch, Jordi Sagrera, Jordi Vivo, Adrià Cubo-Córdoba, Virgilio Martínez-Enamorado, Francisca Rengel-Castro, Virginia García-Entero, Alicia Rodero, Enrique Viguera, Nadin Rohland, Juan Ignacio Morales, María Soto, Swapan Mallick, Artur Cebrià, Pablo García-Borja, Paz Calduch-Bardoll, Pilar Ulloa-Eres, Andrés Carretero, Teresa Espinosa, Beatriz Campderá-Gutiérrez, Paula Pagés-Alonso, Consuelo Vara-Izquierdo, José Martínez-Peñarroya, Samuel Sardà-Seuma, José Manuel Castaño-Aguilar, Sonia López-Chamizo, Ron Pinhasi, Carles Lalueza-Fox, David Reich

## Abstract

It has been unclear how the periods of Roman and later Germanic political control shaped the demography of the Iberian Peninsula and how Iberia differs in these respects from other parts of the Roman Empire. We report genome-wide data from 248 ancient individuals from the largely unsampled period 100-800 CE and co-analyze them with previously reported data. In the Roman era, we document profound demographic transformation, with an influx of people with ancestry from the Central and Eastern Mediterranean in all the areas under study and of North Africans, especially in central and southern Iberia. Germanic (Buri, Suebi, Vandals & Visigoths) and Sarmatian (Alans) took over political control beginning in the 5^th^ century, and although we identify individuals with Germanic-associated ancestry at sites with Germanic-style ornaments and observe that such individuals were closely related across large distances as in the case of two siblings separated by 700 km, for Iberia as a whole, we observe high continuity with the previous Hispano-Roman population. The demographic patterns in Iberia contrast sharply with those in Britain, which showed the opposite pattern of little change in the Roman period followed by great change in the Migration period, and also from demographic patterns in the central Mediterranean where both periods were associated with profound transformation, raising broader questions about the forces that precipitated change over this time.

## Introduction

Rome became a Mediterranean power by the 3^rd^ century BCE and, at its peak, controlled the entire shoreline of the Mediterranean and penetrated substantially into the interior of three continents (Mattingly, 2013). One of Rome’s earliest major conquests was the Iberian Peninsula, controlling the only waterway link between the Atlantic Ocean and the Mediterranean. Because of its unique geographic position and rich physical resources, it was important to the Phoenicians, Greeks, and, later, the Carthaginians, who established trading colonies on its eastern shores (Hodos, 2020).

Romans arrived in Iberia in the late 3^rd^ century BCE during the Second Punic War, while the last independent tribes in the northwest submitted to Roman rule during Augustus’s Cantabrian Wars (Richardson, 2004). The Romans heavily exploited the peninsula’s resources, including metal and minerals, agricultural products, and manpower (Mattingly, 2013). Roman control began to weaken during the “Great Migration” period of the 5^th^ century CE (Harper, 2017; Heather, 2010). Alans, Vandals, and Suebi began settling by 409 CE. The Suebi established a kingdom in 411 CE in the Northwest, which lasted until 585 CE (Harper, 2017), while the Romans reestablished control of most of the peninsula in 429 CE after Vandals and Alans crossed the strait of Gibraltar into North Africa. The Visigothic kingdom, established by 507 CE, would rule until the Muslim conquests of the 8th century (Martínez-Jiménez et al., 2017) and, unlike other Germanic groups, had an enduring influence in the formation of the Hispanic kingdoms’ identity during the Medieval period.

The period of Roman control profoundly changed linguistic, religious, and legal landscapes (Mattingly, 2013). With the exception of the Vasconic language, all the Paleohispanic languages became extinct by the first century CE (Díaz et al., 2019) and were replaced by Latin, whose daughter languages are still widely spoken in the region. *Hispania* became a core province within the Roman Empire, and several Roman emperors came from the Iberian Peninsula including Trajan, Hadrian, and Theodosius I. Germanic kingdoms during Late Antiquity sometimes portrayed themselves as the inheritors of Roman rule, and aspects of Roman administration and governance continued to function under the new rulers; for instance, the Visigothic Kingdom created a hybrid legal system, deeply influenced by Roman law (Martínez-Jiménez et al., 2017).

Whether the profound cultural and political changes driven by the Roman colonization and the Migration Period were accompanied by major demographic changes has been a matter of debate. Demographic transitions in this period in the city of Rome (Antonio et al., 2019), the central Italian Region (Posth et al., 2021), and the Balkans (Olalde et al., 2023) have been investigated with ancient DNA. Genetic studies of the Roman expansion through the Mediterranean detected gene flow from the eastern Mediterranean (where the major imperial urban centers were located) to much of the rest of the empire (Antonio et al., 2019; Olalde et al., 2023; Posth et al., 2021). Although genomic data from a handful of Iberian individuals from this time have been studied (Olalde et al., 2019), the sampling is too thin to provide a full understanding of the demographic impact. Although previous studies and historical estimates (Collins, 2008; Olalde et al., 2019) suggest that the Germanic migrations brought a limited number of new settlers to the peninsula (estimates range around 100,000 Visigoth immigrants and 20,000 Vandals Alans and Suebi (Collins, 2008)), the overall impact of these migrants has not been addressed through genetic data.

## Results

### Data generation

We generated genome-wide data from 248 ancient individuals after in-solution hybridization enrichment with either the ‘1240k’ panel of around 1.2 million single nucleotide polymorphisms (SNPs)(Fu et al., 2015; Mathieson et al., 2015) or the ‘Twist’ panel which targets an enlarged set of 1.4 million SNPs (Rohland et al., 2022).

The newly-reported individuals were excavated from 36 archaeological sites (Figure 1A) dating from the 2^nd^ to the 8^th^ century CE (Figure 1B) and located in four main areas: the southeastern region of present-day Andalusia, the northeastern region around the ancient province capital city of *Tarraco*, the central Iberian Plateau, and the ancient provincial capital of *Emerita Augusta* in the interior southwest. The sites are spread over a diversity of geographical areas with different degrees of Romanization and cover different contexts such as rural/urban, coastal/interior, and cultural affiliations such as Hispano-Roman or Germanic. We highlight 30 individuals from Los Blanes at *Emerita Augusta*, which was briefly the capital of the Suebian kingdom in 440 CE, of which 10 have a Roman period archaeological context and dating, and 20 are associated with the period of Suebian rule. We also analyzed 35 individuals from the Late Antiquity site of Pla de l’Horta (Girona, Catalonia, Spain) with possible Germanic influences in its material culture (Agustí & Llinàs, 2012) and 32 dated to the height of the Roman period from Coracho Basílica (Lucena, Andalusia, Spain), one of the earliest Christian Basilicas in Iberia (Diéguez-Ramírez, 2016).

**FIGURE 1.**
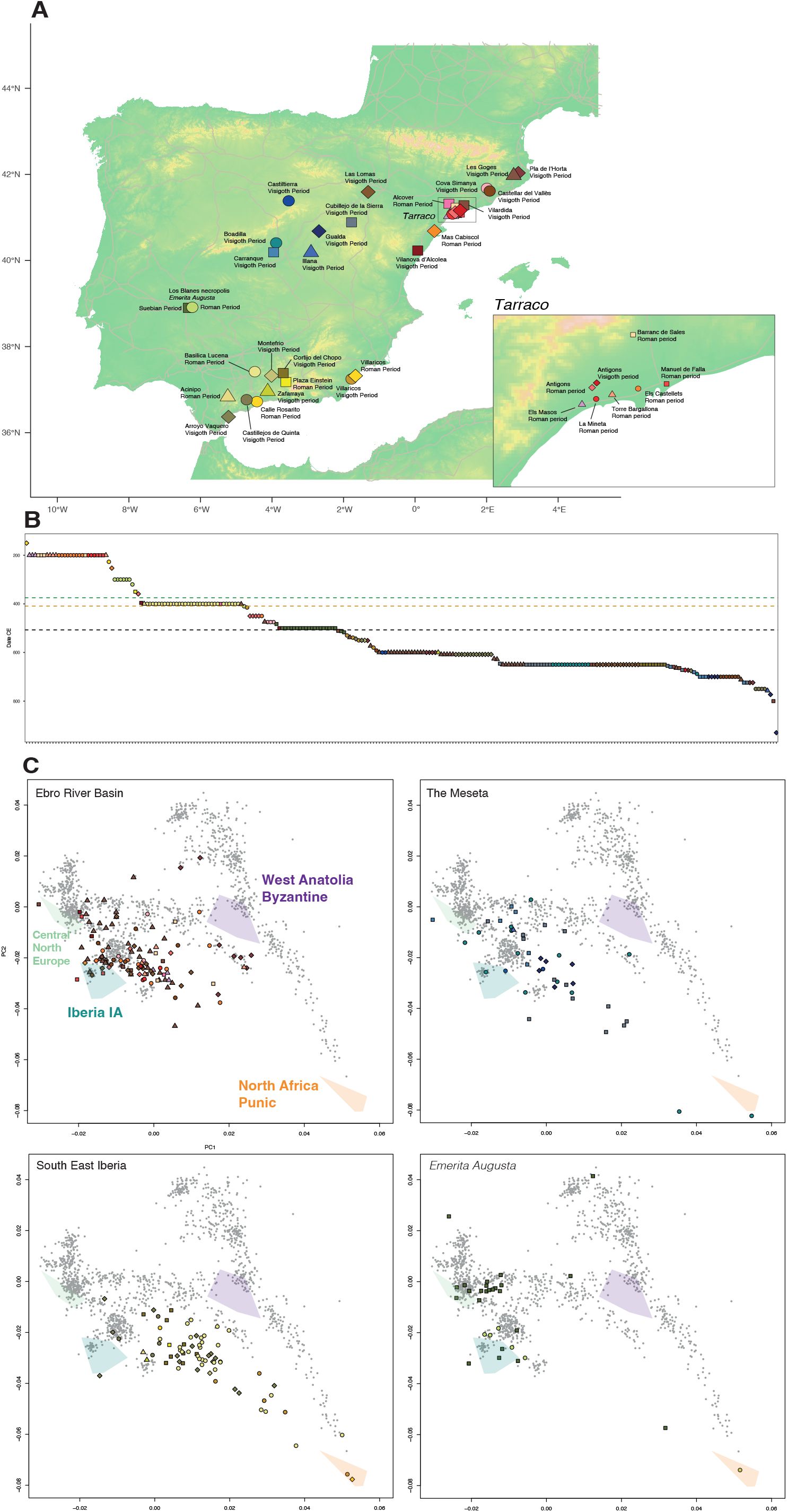
Outline of the newly reported ancient Iberian samples in this study A. Geographical location of archaeological sites relevant to this study. B. Chronological distribution of the individuals dated both by archaeological context or direct radiocarbon dating. The main Germanic arrivals are marked: Arrival of Alans, Suebi, and Vandals (green dotted line), the establishment of the Suebian kingdom (orange dotted line), and the establishment of the Visigoth kingdom (black dotted line). C. Principal Component Analysis (PCA) of West Eurasian genetic variability for the newly reported individuals, classified in 4 main regions of the Iberian Peninsula: Northeast Iberia and the Ebro River basin (red tones), the central Plateau (blue tones), Southeast Iberia (yellow tones), and the city of *Emerita Augusta* (green tones).

For genome-wide analysis, we retained 212 newly reported individuals who yielded more than 15,000 SNPs with overlapping sequencing data and without signals of contamination.

### A turn to extreme ancestry heterogeneity in the Roman period

To develop a qualitative picture of the genetic affinities of Roman and Late Antiquity populations, we performed Principal Component Analysis (PCA). We projected the newly reported ancient individuals onto the two main Principal Components (PC1 and PC2) computed based on patterns of genetic variation among 1036 present-day West-Eurasian (WE) individuals (Figure 1C) genotyped on the Affymetrix Human Origins Array.

The peoples of Iberia became highly genetically heterogeneous in the Roman period, as shown in their remarkable dispersal in the PCA plot (Figure 1C). Their high degree of variability differs markedly from that of the previous Iberian Iron Age populations (blue polygon in PCA). High genetic heterogeneity is observed equally in the Northern and Southern parts of the peninsula.

Ancestry modeling of the ancient individuals with qpAdm allows us to quantify the impacts of this change. We find that only five individuals dated to the Roman period can be modeled using Iberian Iron Age populations as the only ancestry source. However, ancestry from the Iron Age Iberian population is present in admixed form, and even in the Late Antiquity period, we find a handful of individuals (*n*=6, out of 162 individuals dating to post-roman Iberia) consistent with deriving all their ancestry from local Iberian populations.

Population structure during Late Antiquity is remarkably similar to that of Roman Iberia, with similar degrees of ancestry heterogeneity. This suggests either continuing population input from the same areas or sub-structured populations within Iberia that prevented the homogenization of the ancestry profiles.

### An Eastern Mediterranean source for much of the new ancestry in the Roman period

Ancestry analysis of the Roman Period individuals reveals the significant presence of East Mediterranean-related ancestry (Figure 2B) that we modeled with Roman and Byzantine groups from West Anatolia. This signal can be detected as soon as the 3^rd^ century CE at *Tarraco* (Barranc de Sales archaeological site) or at Empúries, and it is attested as well at the Y-chromosome level, with lineages like J-L210, J-L26, and J-Z2177 (Table S1) that are absent during the Iron Age in Iberia to the limits of our resolution, but typical in the Eastern Mediterranean. These patterns are consistent with those observed in other parts of the Roman Empire, such as the Imperial core in Rome and central Italy (Antonio et al., 2019; Posth et al., 2021) and the Balkans (Olalde et al., 2023). The Iberian Peninsula also received this demographic input, which could have its proximal source in the Italian Peninsula, where this ancestry was already widespread, or in more eastern provinces where this ancestry has its distal origin. Unlike the Italian Peninsula or the Balkans, where individuals fully deriving from Eastern Mediterranean populations are documented, in Iberia, this ancestry tends to be always in admixed form. This could be a result of our transect largely missing the Republican period and the early Roman Empire between 1-200 CE, where this immigration might have been at its peak, but it could also relate to more complex processes where the incoming populations had heterogeneous ancestral origins. At the Coracho Basilica (Lucena) (4^th^ century CE), for instance, four individuals could not be modeled by using an East Mediterranean population as their ancestry source, yet they demanded more central Mediterranean sources.

**FIGURE 2.**
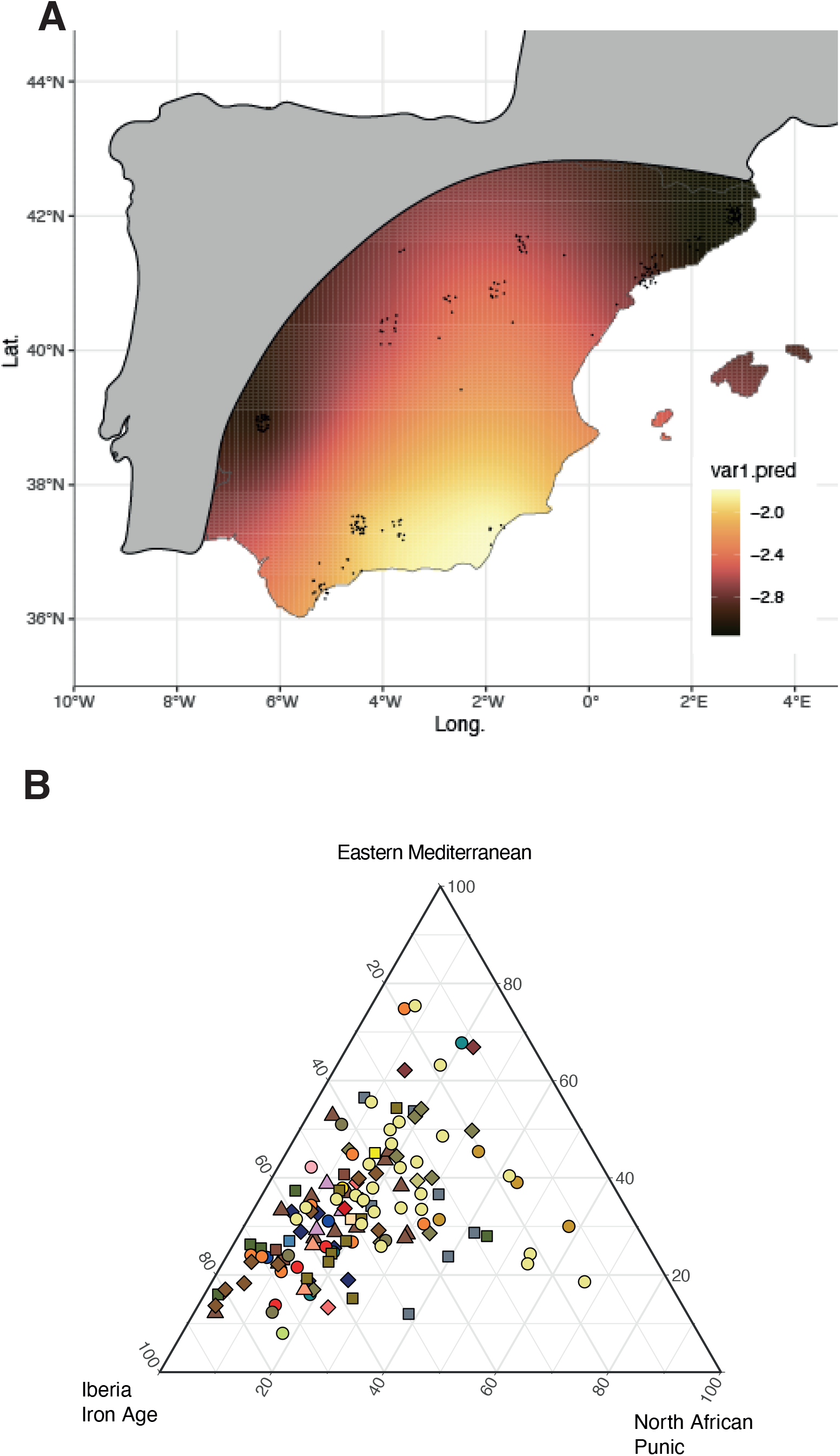
Ancestry modeling of the Roman and post-Roman Iberian population. A. Kriged North African ancestry proportions in the Late Antiquity Iberian Peninsula. Due to a lack of data points, the prediction of Western and Northern Iberia has been cropped. B. Ancestral proportions ancestral proportions of the Roman and post-Roman Iberian population, according to a three-way model.

We detect one individual who can be modeled as originating from the Eastern Mediterranean (I20275), unearthed from the Les Goges archaeological site in present-day Girona (Northeastern Spain). This individual dates from the 8^th^ century CE, suggesting that the close migratory links of Iberia with the rest of the Mediterranean persisted long after the decline of Roman rule. This later interconnection was plausibly mediated by the Eastern Roman (Byzantine) Empire, which still played a major role in the Iberian culture and economy even after direct Roman control ceased.

### Large-scale gene flow originating from North Africa

In addition to the Eastern Mediterranean influence, a North African genetic component became prominent in the Roman period, as reflected in a shift of sampled Iberian individuals toward the PCA space occupied by North Africans (Figure 1C; yellow polygon). Previous work detected North African ancestry during the Roman period in four individuals from one site in Granada (southeast Mediterranean area) (Olalde et al., 2019). We now document this signal ranging from 3-66% of ancestry in all the newly reported Roman-Period individuals (n=32) from this area, excavated at three different urban sites covering the coast (ancient *Malaca* and ancient *Baria*) and the inland (ancient *Acinipo*), and one rural inland site, Coracho Basilica close to present-day Lucena (Córdoba). This signal is also clearly present in nearly all individuals from the seven Late Antiquity sites in this area, with the highest proportions observed in coastal sites such as *Baria* on the eastern part and Arroyo Vaquero on the western part closer to the Gibraltar Strait.

North African ancestry is not restricted to the southeastern Mediterranean and is also detectable in other areas, although generally at lower proportions (Figure 2A). In the Central Iberian Plateau, we detect it in 6 Late Antiquity sites ranging from 4-85% (with only five individuals out of 29 without any Nort African signal, Table S5). At *Emerita Augusta*, one individual is entirely North African (95% ± 5%) out of 9 Roman Period individuals. At eight sites around *Tarraco* in the northeast area, North African ancestry is present in around two-thirds of the individuals (*n*=26 out of 36) ranging from 3-39% (Table S5), hinting that at least in this area, the signal was linked to human mobility attracted to the capital of the *Tarraconensis* province.

Uniparental markers support the autosomal results and provide information about the nature and proximal origin of this influx. In the four areas included in our study, we find mitochondrial and Y-chromosome lineages with a clear African origin, such as mitochondrial haplogroups L1, L2, L3, and the more North Africa-specific U6, or Y-chromosome haplogroup E1b-M81which reaches peak frequencies in present-day northwest Africa. These lineages coexist with others clearly related to the preceding pre-Roman populations, supporting the admixture of North African individuals with the local Iberian population.

Our results demonstrate that North African ancestry was not restricted to one site but was widespread in the southeast area during the Roman and Late Antiquity periods and even reached other regions. These results imply a large-scale demographic influx of both men and women from northwest Africa that brought the two shores of the Gibraltar Strait genetically closer than ever before.

Compared with other imperial regions, the North African gene flow was significantly more intense and widespread and was carried by a large portion of the analyzed individuals. In the Danubian frontier, only two individuals were found to harbor North African ancestry and 1 East African ancestry (Olalde et al., 2023). Similarly, only two individuals could be modeled with North African components in the city of Rome during imperial times (Antonio et al., 2019), and three individuals from central Italy dated to 300 BCE carried high African ancestry proportions (Posth et al., 2021).

The influx of North African ancestry into the Iberian gene pool during the period of Roman control was on a larger scale than in previous periods. Earlier civilizations (such as Phoenicians and Carthaginians) established trading colonies on the Iberian coast and also carried African ancestry. For example, individuals yielding African ancestry can be detected at the Villaricos site during the Punic rule of southeast Iberia (Ringbauer et al. in press). However, their demographic impact on the Iberian population as a whole was not as widespread and intense as the one we observed during the period of Roman rule.

#### Migration Period and Germanic Influence in Iberia

During the “Great Migration Period,” various Germanic-speaking (and other) people spread across Europe. The Iberian Peninsula saw the arrival of several groups, especially Buri, Suebi, Vandals, Alans, and Visigoths. Each has unique origin stories and migration routes, but we can hope to trace their movements by finding genetic signatures of Central - Northern European-like (CNE) ancestry in the autosomes or uniparental markers characteristic of CNE groups and largely absent in Iberia during preceding periods.

Our genetic evidence suggests the presence of a Germanic CNE-related signal in the Iberian Peninsula even before the Great Migration Period. We detect that individual I29631, dating to the early 3^rd^ century CE from ancient *Tarraco* (Manuel de Falla archaeological site), has ∼97% CNE-related ancestry (Table S5). Previous studies have documented sporadic human mobility from beyond the Empire’
ss frontiers before the Migration Period in regions such as the British Isles and the Danubian Frontier (Martiniano et al., 2016; Olalde et al., 2023), and our results extend these observations to Iberia.

The influence of CNE ancestry becomes stronger from the early 5^th^ century CE onwards at specific sites (Figure 3). The impact of Germanic migrants on the Iberian Peninsula is well-documented through both historical records and archaeology and is marked by the establishment of Germanic settlements and the introduction of new cultural elements. We sampled several key archaeological sites with cultural elements traditionally associated with Germanic influences, which provide insights into this process. One of them, Pla de l’Horta in the northeast Mediterranean region, was previously studied by Olalde et al., 2019 with nine individuals, and we now complement the dataset with 31 newly-reported individuals and 5 with higher coverage data. The site consists of a Roman *fundus* (suburban villa) that persisted into post-Roman times. It also includes a necropolis associated with Germanic cultural elements, used from the early 4^th^ century CE and until the early 8^th^ century CE. We count 35 individuals in our dataset who date to such a timeframe and cultural context. In this necropolis, we detect 14 individuals with 27-86% CNE-like ancestry (Table S5), but no individual can be modeled using this ancestry as the only source (the other 21 individuals do not present such ancestry signal). Unlike the Goths that entered the Roman Balkan peninsula in the 4^th^ century, we do not detect Sarmatian-like ancestry (Olalde et al., 2023). The absence of un-admixed CNE individuals is also striking, indicating that Germanic groups mixed with local people; for example, seven individuals carried 12-61% of the local Iberian Iron age-like signal, further implying admixture of the migrant people groups with the local population. Civil unions between Germanic migrants and local Hispano-Romans were not widespread in the years following their arrival. This was, however, overturned when the Visigoth King reared abjured Arianism, converted to Nicean Christianism (Catholicism), and allowed marriages between both groups (Collins, 2008). Yet, we detect that the remaining 21 individuals studied from this archaeological site do not present Germanic-like ancestry, carrying different amounts of the three basal ancestries of the local Hispano-roman population. While we observe that three individuals can be modeled as 100% local Iberian Iron Age, the East Mediterranean ancestry does not surpass 50%, and the North African signal does not surpass 30% among these individuals.

**FIGURE 3.**
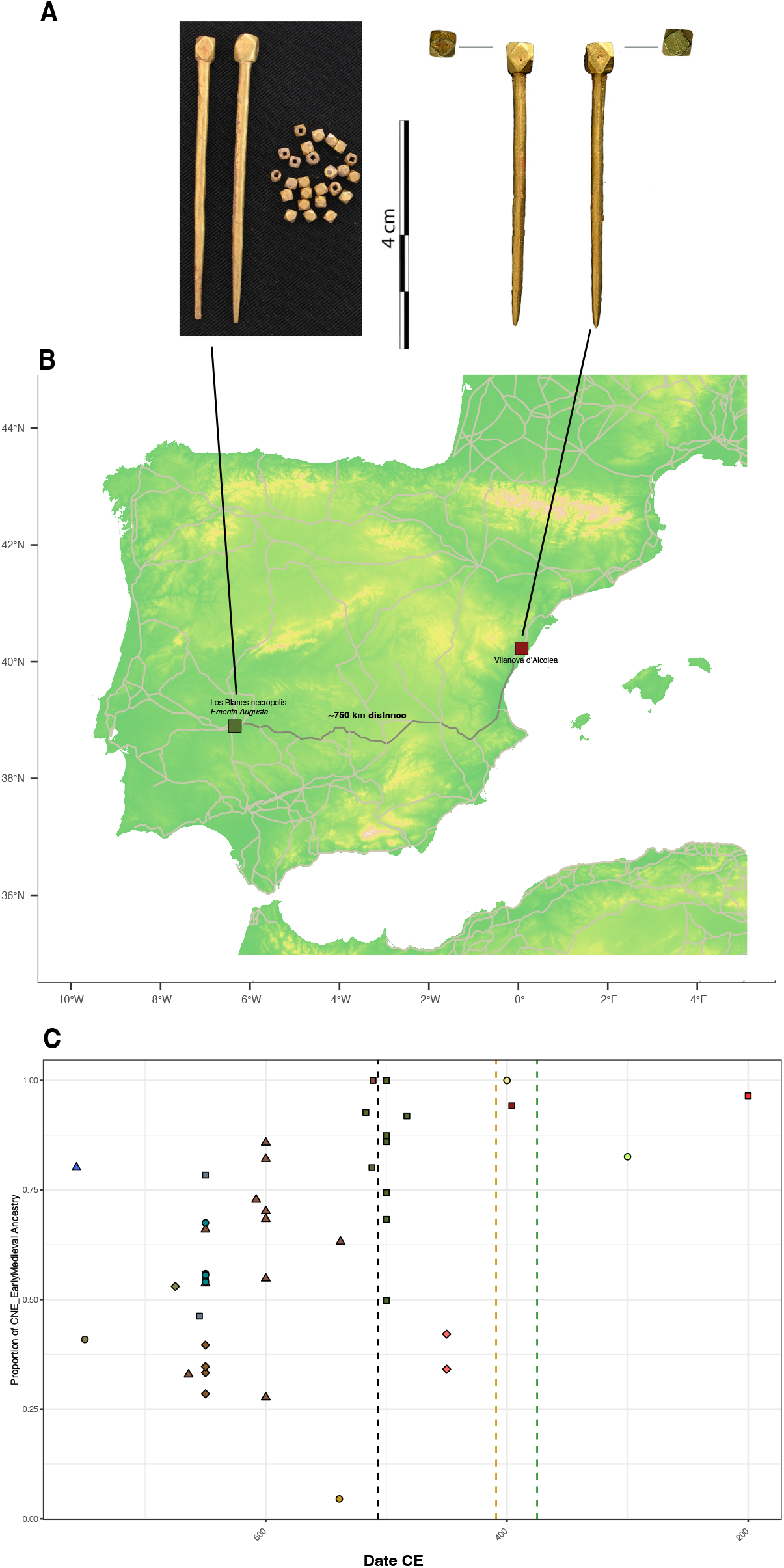
Long-distance connections in the Iberian Peninsula during the Germanic rule period. **A**. Golden pins of Danubian-Pontic tradition are found at Vilanova d’Alcolea and Corralón de Los Blanes (*Emerita Augusta*) archaeological sites. **B**. Location of Vilanova d’Alcolea and Corralón de Los Blanes (*Emerita Augusta*) archaeological sites, where two siblings were found. The shortest path between the two sites along the main Roman roads is indicated, which amounts to about 700 kilometers. **C**. Evolution of the proportion of Germanic-like ancestry across time, with the main Germanic arrivals marked: Arrival of Alans, Suebi, and Vandals (green dotted line), the establishment of the Suebian kingdom (orange dotted line), and the establishment of the Visigothic kingdom (black dotted line).

We also present data from 30 newly reported individuals from the Los Blanes archaeological site at ancient *Emerita Augusta*, the capital of Lusitania province, seven from the Roman period and 23 from the period of Suebian control during the 5^th^ century (Table S1). The later samples correspond to a unique context containing individuals with exceptionally rich grave goods. We find a clear presence of CNE-like ancestry during the Suebian period, with two individuals modeled as 100% CNE-derived and the rest of the individuals who yield such ancestry having 50-93%, while it was largely absent in the individuals dated to the Roman period. In total, 11 individuals dating to this period out of 20 (with enough recovered genetic material for genome-wide analysis) yielded CNE-like ancestry (Table S5). The Los Blanes necropolis is culturally and chronologically associated with the Suebian period. This group is recorded as having migrated directly from *Germania*, crossing Gaul in a matter of years, implying a different population history from the Goths that entered the Empire through the Danubian Limes (Olalde et al., 2023). Although, once again, we do not detect any Sarmatian-like ancestry, we do detect three individuals carrying between 29-50% Central Asian ancestry. According to historical sources, the Suebi entered the peninsula in the early 5^th^ century accompanied by the Alans, a Sarmatian people group, and our genetic data may be reflecting this history. Interestingly, several individuals from this necropolis dated to the Suebian period do not present any signal of CNE-like ancestry, suggesting local population continuity after the arrival of migrant groups. This continuity is also evident in two individuals with a predominant North African genetic legacy at *Emerita Augusta*, one in the period of Roman control and one in the Suebian period.

Overall, during the Great Migration period in Iberia, we detect three individuals who yield 100% CNE-like ancestry: I28358, I23283, and I23772, dated to the 4^th^-6^th^, 5^th^-6^th^, and 5^th^-6^th^ centuries CE respectively, and a further 11 individuals with >80% Germanic-like ancestry (Table S5). We also detect a further 52 admixed individuals who gather varying amounts of CNE-like ancestry, along with individuals without any CNE-like ancestry co-inhabiting in the same settlements. The persistence of Hispano-Roman individuals alongside Germanic-like ancestry points to a scenario of coexistence and gradual admixture rather than whole population replacement. Indeed, 27 individuals harbor both CNE-like and Iberian Iron Age-like ancestry. This indicates some level of intermarriage between the Germanic migrants and the local Hispano-Roman population. However, the overall demographic contribution of the migrant groups was relatively small compared to the enduring presence of the pre-existing population.

Our results suggest Germanic migrations associated with the genesis of the Visigothic kingdom were more the result of an elite replacement than a large-scale migration. Until the conversion of Recared’s king to the Nicean (Catholic) faith in 589 CE, the Arianism of the Visigoths likely acted as a social segregation mechanism. It is precisely under Recared’s rule marriages between Visigoths and locals were allowed. Afterward, the genetic dilution of the Germanic community into the much larger Hispano-Roman population accelerated (Collins, 2008).

#### Close relatives separated by large geographical distance

We find two siblings excavated approximately 700 kilometers apart, which represents by far the largest geographical separation between first-degree relatives in the entire archaeogenetic record (the next being two Copper Age individuals from southern England separated by 5 kilometers) (Table S3, S4). The brother was buried at the Los Blanes necropolis at *Emerita Augusta* and dated to 364-535 cal CE. His sister died 257-409 cal CE and was buried next the Roman *Via Augusta* road at Vilanova d’Alcolea (present-day Castelló, Northeast Spain) in a high-status burial with rich grave goods such as a crystal glass and two gold needles of Danubian-Pontic tradition (García-Borja et al., 2021; Kazanski, 1989), very similar to those found at Los Blanes (Figure 3A). Both siblings exhibited 92-95% CNE-related ancestry, and the brother belonged to the typically Germanic I1 Y-chromosome lineage. This finding highlights the high mobility of Germanic groups during Late Antiquity and their internal connectivity within the peninsula, enabled by the infrastructure established in Iberia during the Roman colonization. Both siblings were buried next to important Roman roads, and according to (https://orbis.stanford.edu/), a trip by foot would have taken ∼22 days (Figure 3B). While the exact migration path of these siblings remains unclear, it is plausible that the sister was born within the Germanic community at *Emerita Augusta*, where her brother was buried and where we have revealed many individuals with her same ancestral origins, and ended her days far away, buried in an isolated but conspicuous burial next to the *Via Augusta*.

## Conclusions

Unlike the limited colonies established by previous civilizations such as the Greeks and Phoenicians/Punics, Roman rule brought significant inflows of Eastern Mediterranean and North African ancestry into Iberia. Historical sources, archaeological, and epigraphic evidence document the arrival of Roman colonists who settled in highly Romanized areas, such as the Guadalquivir Valley starting in the early days of Roman presence in Iberia (Abulafia, 2011; Alvar-Ezquerra, 2007) and our data demonstrate that they precipitated a large-scale demographic change across the whole Peninsula with long-term continuation in the later periods. The contributions of migrants from these regions created a cosmopolitan society interconnected with the broader Mediterranean region. Interestingly, most studies to date have documented the influence of eastern Mediterranean groups on individuals from the central (Italy and the Balkans) and western Mediterranean (Iberia, as we show here). Future archaeogenetic studies in Roman North Africa are necessary to determine if the mobility between Iberia and North Africa was bidirectional.

The genomic analysis of Late Antiquity sites allowed us to identify groups with clear Central-Northern European ancestral origins and to establish close familial connections over hundreds of kilometers. The demographic impact of the Great Migrations has been genomically investigated in other regions of the former Empire, such as Great Britain, the Balkans, and the Italian Peninsula (Antonio et al., 2019; Gretzinger et al., 2022; Olalde et al., 2023; Posth et al., 2021). In England, the potential Anglo-Saxon contribution was estimated to be under 50% in central-south England and most likely in the range of 10%-40% from the analysis of modern genomes (Leslie et al., 2015). This estimate rises to 76% on average when analyzing early Medieval burials, mainly from eastern England (Gretzinger et al., 2022). Both figures point to a large-scale demographic input from continental North Sea regions. In modern Balkans, the potential contribution of the Slavic migrations was estimated to range from 40-60% in places such as modern Serbia to 20% in mainland Greece (Olalde et al., 2023). In the Iberian Peninsula, despite the clear presence of individuals with Germanic ancestry at specific sites, their demographic impact was relatively limited. The characteristic ancestry mixture and diversity typical of Hispano-Roman society remained, and although admixture occurred, the overall genetic landscape did not see radical change. The findings reported in this study contrast with the evidence observed in other European regions where the Migration period peoples’ impact was substantially higher, such as the Anglo-Saxons in the British Isles (Gretzinger et al., 2024), Germanic societies in Italy (Antonio et al., 2019; Posth et al., 2021), or the Slavs in the Balkans (Olalde et al., 2023). However, these findings are not the final piece in the genomic history of this key region. The Iberian Peninsula’s complex and dynamic history, shaped by Romanization, Visigothic rule, Islamic conquests, and subsequent conquest of the Christian kingdoms, presents a unique genetic landscape still to be fully revealed. Given the region’s position as a crossroads between Europe and Africa, the genomic history of Iberia remains an exciting frontier with many questions left to answer. Future genomic research will need to explore deeper layers of population interactions, including the role of local continuity, migration waves, and the influence of later historical events.

## Acknowledgments

We thank the funding agencies for this study: PGC2018-0955931-B-100 grant (MCIU/AEI/FEDER, UE) of the Spanish Ministry of Science of Innovation (C.L.-F), PID2021-124590NB-100 grant of the Spanish Ministry of Science of Innovation (C.L.-F), grant ‘‘Ayudas para contratos Ramón y Cajal’’ funded by MCIN/AEI/10.13039/501100011033 and by ‘‘ESF Investing in your future’’ (I.O.), FPI-2019 (Spanish Ministry of Science of Innovation, BDNS ID:476421) (P.C.). For the ancient DNA work we acknowledge support from NIH grant HG012287; from the Allen Discovery Center program, a Paul G. Allen Frontiers Group advised program of the Paul G. Allen Family Foundation; from John Templeton Foundation grant 61220; and from the Howard Hughes Medical Institute (HHMI). This article is subject to HHMI’s Open Access to Publications policy. HHMI lab heads have previously granted a nonexclusive CC BY 4.0 license to the public and a sublicensable license to HHMI in their research articles. Pursuant to those licenses, the author-accepted manuscript of this article can be made freely available under a CCBY 4.0 license immediately upon publication.

## Author Contributions

I.O., P.C., C.L-F., and D.R. conceived and designed the study.

P.C. wrote the initial manuscript draft

I.O. P.C., C.L-F., and D.R. wrote the manuscript and compiled supplementary materials with the input of all other co-authors.

J.M.J-A., N.C., D.V., J.A.V., A.C., D.B., M.B-Á., J.H-M., C.C., M.I.G-C., JQ., J.R., M.A.C.,

A.A., J.M., R.C., M.LC., J.F.R-P., M.D-G., T.P-R., S.V-Á., J.B., J.S., J.V., V.G-E., A.R., E.V.,

J.A.M., M.S., A.C., P.G-B., A.C., T.E., B.C-G., P.P-A, C.V-I, J.M-P, S.S-S, and P.C. assembled archaeological material and prepared the site descriptions.

N.R. performed laboratory work.

S.M. performed the Bioinformatic data processing

P.C. performed population genetic analyses. I.O., P.C., analyzed data.

I.O., C. L-F and D.R. secured the funding

