## Supplemental_information for "Disparate demographic impacts of the Roman Colonization and the Migration Period in the Iberian Peninsula"

1. Institut de Biología Evolutiva, CSIC-Universitat Pompeu Fabra, Barcelona, Spain
2. BIOMICs Research Group, Department of Zoology and Animal Cell Biology, University of the Basque Country UPV/EHU, Vitoria-Gasteiz, Spain
3. Ikerbasque—Basque Foundation of Science, Bilbao, Spain
4. Department of Human Evolutionary Biology, Harvard University, Cambridge, MA, USA
5. Departamento de Prehistoria y Arqueología, Universidad de Granada, Granada, Spain
6. Universitat de Girona, Girona, Spain
7. Institut Català de Paleoecologia Humana i Evolució Social (IPHES-CERCA), Tarragona, Spain
8. Universitat Rovira i Virgili, Departament d'Història i Història de l'Art, Tarragona, Spain
9. Museo Arqueológico y Etnológico de Lucena, Lucena, Spain
10. Archaeological Museum of Badajoz, Junta de Extremadura, Badajoz, Spain
11. Universidad Isabel I, Burgos, Spain
12. Facultad de Filosofía y Letras, Universidad de Granada, Granada, Spain
13. Dept. Geography, Prehistory and Archaeology, University of the Basque Country UPV/EHU, Vitoria-Gasteiz, Spain
14. BioArCh, Department of Archaeology, University of York, York, United Kingdom
15. University of the Basque Country UPV/EHU, Bilbao, Spain
16. Departamento de Ciencias Históricas, Universidad de Málaga, Málaga, Spain
17. Departamento de Patrimonio Histórico, Ayuntamiento de Estepona, Estepona, Málaga, Spain
18. Museo de Guadalajara, Guadalajara, Spain
19. Museo de Zaragoza, Zaragoza, Spain
20. Empresa CATARQUEOLEGS, SL, Spain
21. Independent Researcher
22. Departamento de Prehistoria, Historia Antigua y Arqueología, Universidad Complutense de Madrid, Madrid, Spain
23. Ministerio de Cultura, Madrid, Spain
24. Departament de Arqueologia, Universitat Autònoma de Barcelona, Barcelona, Spain

25. Departamento de Didáctica de la Matemática, Didáctica de las Ciencias Sociales y de las Ciencias Experimentales, Universidad de Málaga, Málaga, Spain
26. Ayuntamiento de Pizarra, Málaga, Spain
27. Departamento de Prehistoria y Arqueología, Universidad Nacional de Educación a Distancia, Madrid, Spain
28. Departamento de Protohistoria y Colonizaciones, Museo Arqueológico Nacional, Madrid, Spain
29. Department of Genetics, Harvard Medical School, Boston, MA, USA
30. Madrid Institute for Advanced Study - MIAS, Madrid, Spain
31. Departamento de Prehistoria y Arqueología, Universidad Autónoma de Madrid
32. Howard Hughes Medical Institute, Boston, MA, USA
33. Universitat de Barcelona, Barcelona, Spain
34. Centro asociado Alzira-València, Universidad Nacional de Educación a Distancia, València, Spain
35. Departamento de Conservación, Museo Arqueológico Nacional, Madrid, Spain
36. Departamento de Antigüedades Medievales, Museo Arqueológico Nacional, Madrid, Spain
37. CASTRVM patrimonio histórico S.L., Spain
38. Universitat Rovira i Virgili, Departament d'Història i Història de l'Art, Tarragona, Spain
39. Museo Arqueológico de Ronda, Ronda, Málaga, Spain
40. Department of Evolutionary Anthropology, University of Vienna, Vienna, Austria; Human Evolution and Archaeological Sciences, University of Vienna, Vienna, Austria
41. Natural Sciences Museum of Barcelona (MCNB), Barcelona, Spain
42. Broad Institute of MIT and Harvard, Boston, MA, USA

**Contents:**


1. Archeological Overview
2. Direct AMS 14C Dates
3. Ancient DNA laboratory procedures
4. Genomic Data bioinformatic processing
5. Uniparental Marker Calling
6. Kinship analysis and Identification of Identity by descent (IBDs) DNA fragments
7. Aneuploidies
8. PCA
9. F4-statistics

#### 1. Archeological Overview

##### 1.1 Almería, Cuevas del Almazora, Villaricos

<u>Contact person(s):</u> Alicia Rodero

**Brief Settlement history:** The Villaricos archaeological site contains a range of areas for habitation, burial, and industrial activities. This site spans from the Phoenician-Punic era to the Arab-Andalusian period and contains multiple cemeteries and different typologies of burial sites. In total, there have been 1,842 tombs excavated at the site (Almagro-Gorbea, 1984); however, the excavations have been poorly documented. Luis Siret published in 1906 "Villaricos y Herrerías. Antigüedades púnicas, romanas, visigóticas y árabes" with the description of around only 100 tombs.

While Villaricos is best known for its expansive, rock-hewn hypogea, mainly built during Punic times, some of these hypogea were re-used in antiquity. Therefore, amongst the materials held at the Museo Arqueológico Nacional (MAN), some burials were identified by radiocarbon dating as belonging to the Roman-Late Antiquity period.

**Excavation history:** Villaricos has a large burial site that was first explored in the late 1800s. Renowed archaeologist Luis Siret started the systematic excavations in January 1890. The last entry in his journals dates from June, 1914. After Siret, Miriam Astruc and later María José Almagro Gorbea continued the excavations and the study of the materials. Some of the findings were published in 1906 by the Real Academia de la Historia, and more details were given in books such as Astruc's 1951 publication (Astruc, 1951; Siret, 1905). The artifacts from Siret's work were sent to the Museo Arqueológico Nacional (MAN) in Madrid in 1935, where they are still kept (Pereira et al., 1996).

**Geographical Information:** Villaricos is an archaeological site located in the province of Almería, southeastern Spain, near the Mediterranean coast. It sits at the mouth of the Almanzora River.

**Summary of sampled materials:** We sampled five graves dated to the Roman and Late Antiquity period.

**Graves:**

- **Genetic Identifier:** I18195. **Grave Identifier:** 1935/4VILL/T67/1. **Grave Type:** untracable. **Skeletal information:** untracable.**Grave goods:** untracable. **Dating:** 436-581 cal CE (Direct on associated individual 1545±15 BP, PSUAMS-7737). **Ancestry summary:** Iberia_IA__+__WestAnatolia_Roman_Byzantine__+__North_African_Punic.

- **Genetic Identifier:** I18198. **Grave Identifier:** 1935/4VILL/T321/1. **Grave Type:** untracable. **Skeletal information:** untracable. **Grave goods:** untracable. **Dating:** 25-203 calCE (Direct on associated individual 1930±20 BP, PSUAMS-7738). **Ancestry summary:** North_African_Punic.

- **Genetic Identifier:** I18399. **Grave Identifier:** 1935/4VILL/T1159/1. **Grave Type:** untracable. **Skeletal information:** untracable. **Grave goods:** untracable. **Dating:** 432-551 calCE (Direct on associated individual 1570±15 BP, PSUAMS-7742) **Ancestry summary:** CNE_EarlyMedieval__+__North_African_Punic.

- **Genetic Identifier:** I18400. **Grave Identifier:** 1935/4VILL/T1263/2. **Grave Type:** untracable. **Skeletal information:** untracable. **Grave goods:** untracable. **Dating:** 261-416 calCE (Direct on associated individual 1690±20 BP, PSUAMS-7743). **Ancestry summary:** Iberia_IA__+__WestAnatolia_Roman_Byzantine__+__North_African_Punic.

- **Genetic Identifier:** I18401. **Grave Identifier:** 1935/4VILL/T1319-I/1. **Grave Type:** untracable. **Skeletal information:** untracable. **Grave goods:** untracable. **Dating:** 400-600 CE. Context (date on other dated individual(s) of the site). **Ancestry summary:** Iberia_IA__+__WestAnatolia_Roman_Byzantine__+__North_African_Punic.

##### 1.2 Barcelona, Castellar del Valles

Contact person(s): Maite I. Garcia Collado, Juan A. Quiros Castillo, Jordi Roig

**Brief settlement history:** The oldest archaeological evidence recovered at Castellar del Vallès dates to the Early and Middle Neolithic (Coll & Roig, 2006; Roig & Coll, 2010a, 2015, 2010b, 2011, 2018; Ruiz & Subirà, 2010; Subirá et al., 2004). Afterwards, the site was occupied during the Iron Age and the late Roman period (Roig & Coll, 2010b). The early medieval site is initially dated to the first half of the 6th century and it consisted of a small peasant settlement. It lasted until the 8th century with at least one short-distance displacement. The structures recorded included sunken-featured buildings, storage structures, a food preparation area with ovens, a press, a midden, and a possibly irrigated orchard area (Roig & Coll, 2008, 2010b). Between the end of the 6th century and the 7th century, the habitat was displaced to the north (Coll & Roig, 2006; Roig & Coll, 2010b).

The cemetery was located east of the first residential area (Roig, 2015, 2019). The necropolis had a linear outline arranged in a northwest-southeast direction. There were 21 burials and they were organised in three groups. Four types of burials were recorded: tegulae burials, burials in mixed materials, slabs burials, and simple pits. The total number of individuals identified was 33. Most of them were in primary deposits and supine positions. There were also secondary deposits as reductions and on tumuli on burials. No conventional grave goods were recovered and the only elements found in burials were eggs.

**Excavation History:** The archaeological site of Plaça Major de Castellar del Vallès was discovered in the late 1980s by J. M. Coll and J. Roig while they were doing the municipal inventory of archaeological heritage (Coll & Roig, 2006). Still, the first excavation did not take place until 2003, when the council of Castellar del Vallès developed the zone behind Can Torras, so far used as vegetable gardens. The excavation covered approximately 2200 $m^2$ (Coll & Roig, 2005, 2006; Roig & Coll, 2008). In 2005 an underground car park was to be built in the neighbouring plot, so 6750 $m^2$ were excavated as a mitigation measure (Roig & Coll, 2007, 2008). The next year a smaller area of 360 $m^2$ corresponding to a playground was intervened in order to join some of the structures dug in the previous campaigns (Roig, 2006). Finally, in 2008 the completion of the development of the area and the construction of a residential building led to the excavation of 4000 $m^2$ more (Molina & Roig, 2008; Roig & Coll, 2010a, 2010b). All these archaeological interventions were carried out by the company Arrago under the direction of J. M. Coll and J. Roig.

**Geographic Information:** Plaça Major de Castellar del Vallès is the name of an archaeological site located at the main square of the current homonymous town, in the region of western Vallès, province of Barcelona. Originally this site was considered to be two separate entities, so it was given two different names: Plaça Major de Castellar del Vallès and Horts de Can Torras. However, it was soon understood that the structures from both interventions were part of the same settlement. The UTM ETRS89 coordinates of the site are 424011 4607621 and it is 333 meters above sea level. Castellar del Vallès is 27 km northwest of Barcelona, 74 km southwest of Girona and 8 km northeast of the seat of the bishopric of Egara, present Terrasa. The site was situated at the foot of the Catalan Pre-Coastal mountain range, in the upper Ripoll valley, less than 1 km from the riverbed. In addition, the area is crossed by many seasonal watercourses originating in the nearby mountains, such as Canyelles torrent. This location would have provided plenty of fertile fields for agrarian activities and, thanks to the vicinity to the mountains, forest and other altitude resources would have been readily available too.

**Summary of sampled materials**: A minimum number of 32 individuals preserved skeletal remains for osteoarcheaological analysis (García-Collado, 2020). Macroscopic preservation was good. The subadult to adult ratio of the buried population was 0.35 and the male to female ratio was 1.75. Life expectancy at birth was calculated to be 26.5 years. Quality of the collagen recovered was also good. Carbon and nitrogen stable isotope ratios of 21 individuals revealed quite a homogeneous diet with some individuals exclusively eating C3 plants and others with small but consistent consumption of C4 crops. Animal protein intake was between moderate and abundant and there may have been some contribution of aquatic resources too. Sampling for ancient DNA analysis was aimed at representing all demographic categories and funerary rituals. Teeth, preferentially molars, were sampled.

**Graves:**

- **Genetic Identifier:** I19877. **Grave Identifier:** PMCV 14-1-109. **Grave Type:** Slabs burial. Individual in primary supine position, N-S orientation. **Skeletal information:** Adult sp (>20 years), undertermined sex. **Grave goods:** No grave goods. **Dating:** 500-800 CE. Context of archeological findings and archeological layer. **Ancestry summary:** Iberia_IA. **Additional information:** $\delta^{13}C$ = -18.8‰, $\delta^{15}N$ = 9.0‰, %C = 40.5, %N = 13.8, C/N = 3.4, %coll = 1.6

- **Genetic Identifier:** I19879. **Grave Identifier:** PMCV 19-2-141. **Grave Type:** Tegulae burial. Individual in reduction (secondary deposit). **Skeletal information:** Young adult (22-25 years), probably male. **Grave goods:** egg. **Dating:** 500-800 CE. Context of archeological findings and archeological layer. **Ancestry summary:** Excluded from genome-wide analysis due to low coverage.

- **Genetic Identifier:** I19880. **Grave Identifier:** PMCV 22-2-116. **Grave Type:** Tegulae burial. Individual in reduction (secondary deposit). **Skeletal information:** Adult sp (>20 years), probably male. **Grave goods:** No grave goods. **Dating:** 500-800 CE. Context of archeological findings and archeological layer. **Ancestry summary:** Iberia_IA. **Additional information:** $\delta^{13}C$ = -18.9‰, $\delta^{15}N$ = 8.8‰, %C = 40.1, %N = 13.3, C/N = 3.5, %coll = 0.9

- **Genetic Identifier:** I19881. **Grave Identifier:** PMCV 43-1-350. **Grave Type:** Slabs burial. Individual in primary supine position, NW-SE orientation. **Skeletal information:** Young adult (22-36 years), female. **Grave goods:** No grave goods. **Dating:** 500-800 CE. Context of archeological findings and archeological layer. **Ancestry summary:** Excluded from genome-wide analysis due to low coverage. **Additional information:** $\delta^{13}C$ = -18.9‰, $\delta^{15}N$ = 9.0‰, %C = 41.6, %N = 14.1, C/N = 3.4, %coll = 1.7

- **Genetic Identifier:** I19882. **Grave Identifier:** PMCV 44-1-469/470 #1. **Grave Type:** Tegulae burial. Individual in primary supine position, N-S orientation **Skeletal information:** Young adult (17-27 years), male. **Grave goods:** No grave goods. **Dating:** 500-800 CE. Context of archeological findings and archeological layer. **Ancestry summary:** Iberia_IA__+__WestAnatolia_Roman_Byzantine. **Additional information:** $\delta^{13}C$ = -18.8‰, $\delta^{15}N$ = 8.9‰, %C = 41.3, %N = 14.1, C/N = 3.4, %coll = 1.2

- **Genetic Identifier:** I19883. **Grave Identifier:** PMCV 45-1-412. **Grave Type:** Tegulae burial. Individual in primary supine position, NW-SE orientation. **Skeletal information:** Adult sp (>20 years), male. **Grave goods:** No grave goods. **Dating:** Radicarbon dated to 437-599 calCE (1535±20 BP, PSUAMS-10246). **Ancestry summary:** Iberia_IA__+__WestAnatolia_Roman_Byzantine. **Additional information:** male. $\delta^{13}C$ = -19.3‰, $\delta^{15}N$ = 9.0‰, %C = 41.6, %N = 14.8, C/N = 3.3, %coll = 2.5

- **Genetic Identifier:** I19884. **Grave Identifier:** PMCV 48-1-353. **Grave Type:** Slabs burial. Undetermined position. **Skeletal information:** Adult sp (>20 years), undertermined sex. **Grave goods:** No grave goods. **Dating:** 500-800 CE. Context of archeological findings and archeological layer. **Ancestry summary:** Excluded from genome-wide analysis due to low coverage. **Additional information:** $\delta^{13}C$ = -19.0‰, $\delta^{15}N$ = 9.1‰, %C = 30.7, %N = 10.8, C/N = 3.3, %coll = 1.6

- **Genetic Identifier:** I19885. **Grave Identifier:** PMCV 49-2-419. **Grave Type:** Tegulae burial. Individual in tumulus on grave (secondary deposit). **Skeletal information:** Adult sp (>20 years), undertermined sex. **Grave goods:** No grave goods. **Dating:** 500-800 CE. Context of archeological findings and archeological layer. **Ancestry summary:** Excluded from genome-wide analysis due to low coverage.

##### 1.3 Barcelona, Sant Llorenç del Munt, Cova Simanya Gran

Contact person(s): Juan Ignacio Morales, María Soto & Artur Cebrià.

**Brief settlement history:** Cova Simanya is a karstic complex made up of different galleries. The most important of these is Simanya Gran, which shows evidence of human occupation from the Middle Paleolithic to historical times. The remains included in this work come from levels P1 and W1 located in a sub-surface position in the "Main Gallery" and in the "West Gallery" of Simanya Gran respectively. The P1 level has a non-uniform distribution throughout the Main Gallery with a significant presence of human remains. These appear completely disarticulated and accumulated randomly, probably as a result of the periodic flow of water through the gallery and the progressive remobilization of the remains (Cebrià & Morales, 2023).

The W1 level of the "West Gallery", although it has only been partially excavated, has a higher density of remains than level P1. The presence of ceramic and vitreous fragments alongside human remains is noteworthy. Both the preliminary characteristics of the anthropological set and the spatial characteristics of the gallery suggest a possible burial chamber or burial area. The C14 datings from two different individuals, one from level P1 (431 ± 57 cal CE) and another from level W1 (450 ± 61 cal AD), place the burial time within the 5th century CE (Cebrià & Morales, 2023).

**Excavation History:** Excavations in Simanya Gran began in 2020 with the aim of reconstructing the general stratigraphic sequence of the site and were primarily focused on the study of the Pleistocene record. The sample from levels P1 and W1 was recovered during exploratory surveys during the first excavation campaign (Cebrià & Morales, 2023).

**Geographic Information:** By Complex Simanya (ETRS89 31N 417522.2(E) 4614082.4 (N), 875 m above sea level) we refer to a group of caves that open up aligned on the northwest slope of Montcau, and which are part of the same karstic system or share the same speleogenesis dynamics. The caves that make up this complex are the Triangle Cave (or Dry Cave), Simanya Petita, Simanya Gran, the Canal Cave (or Torrent Cave), and Angel's Cave. For the first three caves, there is clear evidence that they are part of the same original karstic system, although in the case of Simanya Petita the physical connection between galleries could not be reached by a few meters. This set is what we call Cova Simanya sensu lato, using the abbreviations ST (Simanya Triangle), SP (Simanya Petita) and SG (Simanya Gran) for the longitudinal galleries, as well as SPO (Simanya Goose Pass) for the cross gallery that connects them. The Canal and Angel's Caves currently appear as independent galleries, although they were likely part of the same drainage system that generated Cova Simanya (Cebrià & Morales, 2023).

**Summary of sampled materials:** The remains selected for DNA sampling are 3 dental pieces, belonging to three different individuals, recovered during the 2020 campaign. Two of the remains SG20.02.54 and SG20.02.50 come from level P1, while SG20.07.31 comes from level W1 (Cebrià & Morales, 2023).

**Graves:**

- **Genetic Identifier:** I28352. **Grave Identifier:** SG20.02.54. **Grave Type:** comingled remains from potential collecive inhumation. **Skeletal information:** undefined sex, 4.5-5.5 years old. **Grave goods:** unattributable. **Dating:** 377-474 CE IntCal20 on associated individual (1640 ± 30 BP, Beta-Beta-574953). **Ancestry summary:** Iberia_IA__+__WestAnatolia_Roman_Byzantine.

- **Genetic Identifier:** I28353. **Grave Identifier:** SG20.07.31. **Grave Type:** comingled remains from potential collecive inhumation **Skeletal information:** undefined sex, adult. **Grave goods:** unattributable. **Dating** 350-500 CE, Context (date on associated tooth (1640 BP, Beta-Beta-574953)**. Ancestry summary:** Iberia_IA__+__WestAnatolia_Roman_Byzantine.

- **Genetic Identifier:** I28354. **Grave Identifier:** SG20.02.50. **Grave Type:** comingled remains from potential collecive inhumation **Skeletal information:** undefined sex, adult. **Grave goods:** unattributable. **Dating:** Direct Radiocarbon dated to 377-474 CE, Context Beta-Beta-574953)**. Ancestry summary:** Iberia_IA__+__WestAnatolia_Roman_Byzantine__+__North_African_Punic.

#### 1.4 Castelló, La Vilanova d'Alcolea, L'Hostalot-Ildum

Contact person(s): Paz Calduch Bardoll, Pilar Ulloa Eres & Pablo García Borja.

**Brief settlement history:** The Roman remains of Hostalot were disclosed by J. Senent Ibáñez in 1923 in his study on the layout of the Via Augusta between the Sénia and Millars rivers, relating the place to the post of the Roman Empire called Ildum. Years later, in different works on the layout of the Via Augusta as it passes through Valencian lands, this initial proposal was strengthened, identifying the Hostalot with the imperial *mansio* known as Ildum in the Antonine itinerary, the vessels of Vicarello and the Anonymous of Ravenna (Arasa-i-Gil, 2013).

**Excavation History:** The first archaeological excavations were carried out in 1986 and 1987 under the direction of F. Arasa (Arasa-i-Gil, 2022) in order to document the perimeter walls of the best preserved building or Sector I. In 1992 a new archaeological excavation was carried out motivated by the discovery of a complete milestone in the course of the works for the expansion of the CV-10 highway. This action was directed by P. Ulloa and focused on an extension of 590 m² next to the route of the road (Ulloa-Chamorro & Grangel-Nebot, 1996). The latest interventions date from the years 2018, 2021 and 2022, linked to a project to enhance and study the enclave promoted by the local government of the municipality (García-Borja et al., 2021).

**Geographic Information:** The Hostalot-Ildum archaeological site is located in the municipality of Vilanova d'Alcolea (Castellón), 263 m above sea level. Most of the site extends along the southern margin of the Carrasqueta ravine, although the remains are scattered on both sides of the CV-10 highway which, in this place, is proposed to follow the same route as the Via Augusta and than the Camino Real between Barcelona and Valencia. It presents two moments of occupation linked to buildings that provided services to travelers passing through the adjacent roads, an older one from Roman times and a more recent one dated between the 16th and 19th centuries. For this reason, it receives the place name of the old hostel built on the Roman ruins.

**Summary of sampled materials:** In the course of the archaeological excavations carried out in 1992, the remains of a skeleton in anatomical connection corresponding to a primary burial located in the courtyard of a building identified as the *Hospidium* of the *mansio* were located. The body was located deposited inside a NE-SW orientation pit excavated in the natural geological sediment. The skeleton appeared in a supine position, with the head tilted slightly to the right, with the right arm stretched parallel to the body and the left arm flexed with the hand on the pubis(García-Borja et al., 2021; Ulloa-Chamorro & Grangel-Nebot, 1996).

**Graves:**

- **Genetic Identifier:** I22103. **Grave Identifier:** UE-3006. **Grave Type:** no visible burial construction, NE-SW orientation, supine position. **Skeletal information:** Adult, female. **Grave goods:** glass vase, placed next to her right hand, and two gold pins placed on her chest (García-Borja et al., 2021; Ulloa-Chamorro & Grangel-Nebot, 1996). **Dating:** Direct Radiocarbon 277-409 cal CE (1710± 20 BP, PSUAMS-8812) **Ancestry summary:** CNE_EarlyMedieval__+__North_African_Punic. Additional Information: Sister of sample I23273 buried in Mérida, Blanes necropolis.

##### 1.5 Córdoba, Lucena, Coracho Basílica

Contact person(s): Archeology: Daniel Botella Ortega. Anthropology: Juan Pablo Diéguez Ramírez & Ricardo Ortega

**Brief settlement history**: The archeological site consists of a martyrium basilica with different phases or cultural layers. It was founded during the Constantinian period, had changes during the Byzantine period, and was adapted for Mozarabic rituals in the 6th-7th centuries. The site was abandoned in the 8th century. It is surrounded by a large late-antiquity necropolis with burials in graves that are oriented from east to west. The graves may be covered with stones or flat or curved tiles, or they may have no covering at all. In some graves, the remains of several individuals have been found. The earlier bodies are often grouped either at the head or the feet of the most recently buried person. Few grave goods have been found (Diéguez-Ramírez, 2016).

**Excavation History**: The archeologica work was carried out due to the construction of the A-45 highway between Lucena and Antequera. The site is located at the northern edge of the Lucena Sur-Encinas Reales Norte section. Earlier, a surface survey was conducted in 2000, where this site was identified by the director of the project, Fernando Penedo Cobo. In March 2023, Francisca Casado Trenas and María Cabeza Liébana Sánchez were involved in the work. Due to disagreements in interpreting the findings between the excavation team and the municipal archaeology department of Lucena, the Provincial Heritage Commission of Córdoba, in a resolution issued in April 2004, authorized the relocation and reconstruction of the basilica remains on land provided by the Lucena City Council. This is where they can be seen today.

**Geographic Information**: The site is located in the southern area of the Lucena municipality, in a region called the "campiña alta cordobesa," the landscape consists of olive groves on Triassic marl and gypsum soils, shaped by the nearby Anzur River. The martyrium basilica and its associated necropolis were built on one of these rounded hills, very close to the old Benamejí road and east of the ancient Corduba-Malaca route.

**Summary of sampled materials:** The graves from this site are dated to the 4$^{rd}$ century CE. They contain the remains of individuals of various ages and sexes. Burial 3 contained various grave goods, such as blue and black glass beads, bone pieces, a perforated shell, jet beads, and other jewelry-related items. the majority of the other graves contain no grave goods. There are also subadult graves, like Burial 94, containing the remains of a 4-6-year-old female with no grave goods, and Burial 168, with a 10-12-year-old female, also without grave goods. Anthropological studies determined specific conditions in certain graves (Diéguez-Ramírez, 2016; Diéguez-Ramírez et al., 2022). For example, Burial 34 has osteomyelitis in the right tibia and osteophytes in the vertebrae (Ortega-Ruiz & Leggio, 2024).

**Graves:**

- **Genetic Identifier:** I35803. **Grave Identifier:** Burial 3 (CF 96), ind 1. **Grave Type:** Rectangular simple pit with double burial. One individual is lying on their back, with the bones of the second individual at their feet, first individual sampled. Orientation is E-W. **Skeletal information:** Adult > 45 years old, male. **Grave goods:** Blue glass beads for necklaces, black glass beads for bracelets, a bone piece that could be part of a necklace, a spindle-shaped bone bead, a fish-shaped bone bead, a perforated shell for necklaces, jet beads, amber bead, earring fragment, silver clasp chain, small black glass beads for necklaces, silver clasp chain, silver threads for bracelets. **Dating:** Archeologically dated to 300 – 400 CE. **Ancestry summary:** Iberia_IA__+__WestAnatolia_Roman_Byzantine__+__North_African_Punic.

- **Genetic Identifier:** I35804. **Grave Identifier:** Burial 4, ind. 1 **Grave Type:** Rectangular simple pit with double burial. One individual is lying on their back, with the bones of the second individual at their feet, first individual sampled. Orientation is E-W. **Skeletal information:** Young Adult, 18-29 years old, Female. **Grave goods:** no grave goods. **Dating:** Archeologically dated to 300 – 400 CE. **Ancestry summary:** Iberia_IA__+__WestAnatolia_Roman_Byzantine__+__North_African_Punic.

- **Genetic Identifier:** I35805. **Grave Identifier:** Burial 34, ind. 1 **Grave Type:** Oval simple pit with an individual lying on their back. Sandstone slab at the head as an indicator or cover. Orientation is E-W. **Skeletal information:** Adult, >30 years old, Male. **Grave goods:** no grave goods. **Dating:** Archeologically dated to 300 – 400 CE. **Ancestry summary:** Iberia_IA__+__WestAnatolia_Roman_Byzantine__+__North_African_Punic. **Additional information:** Osteomyelitis in right tibia. Osteophytes in vertebrae and herniated discs.

- **Genetic Identifier:** I35806. **Grave Identifier:** Burial 94, ind. 1 (subadult) **Grave Type:** Oval simple pit. **Skeletal information:** Child, 4-6 years old, Female. **Grave goods:** no grave goods**. Dating:** Archeologically dated to 300 – 400 CE. **Ancestry summary:** Iberia_IA__+__WestAnatolia_Roman_Byzantine__+__North_African_Punic**. Additional Information:** Shovel-shaped incisors and Wormian bones.

- **Genetic Identifier:** I35898. **Grave Identifier:** Burial 27, ind. 1 **Grave Type:** Rectangular simple pit with a double burial. One individual lying on their back, with the ossuary of the second individual at their feet. Orientation is E-W. **Skeletal information:** Adult, >30 years old, Male. **Grave goods:** no grave goods. **Dating:** Archeologically dated to 300 – 400 CE. **Ancestry summary:** Iberia_IA__+__WestAnatolia_Roman_Byzantine__+__North_African_Punic. **Additional information:** Joint pathologies.

- **Genetic Identifier:** I35899. **Grave Identifier:** Burial 37, ind. 1 **Grave Type:** Trapezoidal simple pit with a double burial. One individual lying on their back, with the ossuary of the second individual at their feet. Orientation is E-W. **Skeletal information:** Adult,20-25 years old, Undetermined sex. **Grave goods:** no grave goods. **Dating:** Archeologically dated to 300 – 400 CE. **Ancestry summary:** Iberia_IA__+__WestAnatolia_Roman_Byzantine__+__North_African_Punic.

- **Genetic Identifier:** I35942. **Grave Identifier:** Burial 5, ind. 2 **Grave Type:** Rectangular simple pit with double burial. One individual is lying on their back, with the bones of the second individual at their feet, second individual sampled. Orientation is E-W. **Skeletal information:** Adult, 25-35 years old, Female. **Grave goods:** no grave goods. **Dating:** Archeologically dated to 300 – 400 CE. **Ancestry summary:** Iberia_IA__+__WestAnatolia_Roman_Byzantine__+__North_African_Punic.

- **Genetic Identifier:** I35943. **Grave Identifier:** Burial 64, ind. 1 **Grave Type:** Oval simple pit, covered with well-squared sandstone slabs. One individual lying on their back. Orientation is E-W. **Skeletal information:** Infant, 2-3 years old, undetermined sex. **Grave goods:** no grave goods**. Dating:** Archeologically dated to 300 – 400 CE. **Ancestry summary:** Iberia_IA__+__WestAnatolia_Roman_Byzantine__+__North_African_Punic.

- **Genetic Identifier:** I35944. **Grave Identifier:** Burial 4, ind. 2 **Grave Type:** Rectangular simple pit with double burial. One individual is lying on their back, with the bones of the second individual at their feet, second individual sampled. Orientation is E-W. **Skeletal**

**information:** Adult, 25-35 years old, Male. **Grave goods: no grave goods. Dating:** Archeologically dated to 300 – 400 CE. **Ancestry summary:** Iberia_IA__+__WestAnatolia_Roman_Byzantine__+__North_African_Punic.

- **Genetic Identifier:** I35945. **Grave Identifier:** Burial 7, ind. 1 **Grave Type:** Rectangular simple pit with an individual buried lying on their back. Orientation is E-W. **Skeletal information:** Adult, >35 years old, Male. **Grave goods:** no grave goods. **Dating:** Archeologically dated to 300 – 400 CE. **Ancestry summary:** Iberia_IA__+__WestAnatolia_Roman_Byzantine__+__North_African_Punic.

- **Genetic Identifier:** I35946. **Grave Identifier:** Burial 86, ind. 2 **Grave Type:** Oval simple pit covered with sandstone slabs, including reused door hinge elements. Orientation is E-W. **Skeletal information:** Young Adult, 16-25 years old, Female. **Grave goods:** no grave goods. **Dating:** Archeologically dated to 300 – 400 CE. **Ancestry summary:** Italy_Sicily_Punic__+__North_African_Punic. **Additional information:** Osteoporosis in the hip, pregnancy marks and childbirth with injury.

- **Genetic Identifier:** I35947. **Grave Identifier:** Burial 231, ind. 2 **Grave Type:** Oval simple pit, covered with well-squared sandstone slabs and vertical brick walls, containing a double burial with individuals lying on their backs. Orientation is E-W. **Skeletal information:** Adult, 18-25 years old, Male. **Grave goods:** no grave goods. **Dating:** Archeologically dated to 300 – 400 CE. **Ancestry summary:** Iberia_IA__+__WestAnatolia_Roman_Byzantine__+__North_African_Punic. **Additional information:** *Cribra orbitalia*, lower body periostitis.

- **Genetic Identifier:** I35948. **Grave Identifier:** Burial 168, ind. 1 **Grave Type:** Simple square upper pit with vertical brick walls. Lower oval pit. Individual lying on their back, with the ossuary of the second individual at the feet. Orientation is E-W. **Skeletal information:** Child, 10-12 years old, Female. **Grave goods:** no grave goods. **Dating:** Archeologically dated to 300 – 400 CE. **Ancestry summary:** Iberia_IA__+__WestAnatolia_Roman_Byzantine__+__North_African_Punic.

- **Genetic Identifier:** I35949. **Grave Identifier:** Burial 27, ind. 2 **Grave Type:** Oval simple pit with triple burial, containing two inhumations and an ossuary at the foot of both. Individuals lying on their backs, with the ossuary of the third individual at the feet. Orientation is E-W. **Skeletal information:** Adult, 30-35 years old, Female**. Grave goods:** no grave goods. **Dating:** Archeologically dated to 300 – 400 CE. **Ancestry summary:** Iberia_IA__+__WestAnatolia_Roman_Byzantine__+__North_African_Punic.

- **Genetic Identifier:** I36195. **Grave Identifier:** Burial 103, ind. 1 **Grave Type:** Trapezoidal simple pit with burial in supine position. Orientation is E-W. **Skeletal information:** Adult, 35-45 years old, Male**. Grave goods:** no grave goods. **Dating:** Archeologically dated to 300 – 400 CE. **Ancestry summary:** Iberia_IA__+__WestAnatolia_Roman_Byzantine__+__North_African_Punic.

- **Genetic Identifier:** I36196. **Grave Identifier:** Burial 25, ind. 1 **Grave Type:** Rectangular simple pit with double burial. Orientation is E-W. **Skeletal information:** Adult, 30-40 years old, Male**. Grave goods:** no grave goods. **Dating:** Archeologically dated to 300 – 400 CE. **Ancestry summary:** Iberia_IA__+__WestAnatolia_Roman_Byzantine__+__North_African_Punic.

- **Genetic Identifier:** I36197. **Grave Identifier:** Burial 35, ind. 1 (juvenile) **Grave Type:** Oval simple pit with double burial, individual in supine position, with the skull next to the head of the other individual, while the rest of the body of the second individual is at the feet. Orientation is E-W. **Skeletal information:** Child, 6-10 years old, Undetermined sex. **Grave goods:** no grave goods. **Dating:** Archeologically dated to 300 – 400 CE. **Ancestry summary:** WestAnatolia_Roman_Byzantine__+__North_African_Punic.

- **Genetic Identifier:** I36198. **Grave Identifier:** Burial 169, ind. 2 **Grave Type:** Oval simple pit, with a flat rectangular sandstone slab over the head. Double burial with the first individual in a supine position, and the osseous remains of the second individual at the feet. Orientation is E-W. **Skeletal information:** Child, 9-20 years old, Undetermined sex**. Grave goods:** no grave goods. **Dating:** Archeologically dated to 300 – 400 CE.) **Ancestry summary:** No fitting Model.

- **Genetic Identifier:** I36199. **Grave Identifier:** Burial 84, ind. 1 **Grave Type:** Oval-shaped simple pit with a covering of well-squared sandstone slabs. Double burial, with the first individual in a supine position and the osseous remains of the second individual at the feet, although the skull appears next to the more recent one. Orientation is E-W. **Skeletal information:** Adult, 25-30 years old, Male. **Grave goods:** no grave goods. **Dating:** Archeologically dated to 300 – 400 CE. **Ancestry summary:** Italy_Sicily_Punic__+__North_African_Punic. **Additional information:** Schmorl's nodes.

- **Genetic Identifier:** I37206. **Grave Identifier:** Burial 257, ind. 1 **Grave Type:** Simple pit with a brick covering. Double burial, with individuals in a supine position. Orientation is E-W. **Skeletal information:** Adult, 20-30 years old, Female**. Grave goods:** no grave goods. **Dating:** Archeologically dated to 300 – 400 CE. **Ancestry summary:** Iberia_IA__+__WestAnatolia_Roman_Byzantine__+__North_African_Punic.

- **Genetic Identifier:** I37207. **Grave Identifier:** Burial 80, ind. 1 **Grave Type:** Oval simple pit. Double burial, with individuals in a supine position. Orientation E-W. **Skeletal information:** Adult, 25-30 years old, Male. **Grave goods:** no grave goods. **Dating: Archeologically dated to 300 – 400 CE. Ancestry summary:** Iberia_IA__+__WestAnatolia_Roman_Byzantine__+__North_African_Punic. **Additional information:** Schmorl's nodes, *ponticulus posticus.*

- **Genetic Identifier:** I37208. **Grave Identifier:** Burial 100, ind.1 **Grave Type:** Oval simple pit. Double burial, with individuals in a supine position. Orientation E-W. **Skeletal information:** Adult, 25-35 years old, Undetermined sex. **Grave goods:** no grave goods. **Dating:** Archeologically dated to 300 – 400 CE. **Ancestry summary:** Iberia_IA__+__WestAnatolia_Roman_Byzantine__+__North_African_Punic.

- **Genetic Identifier:** I37209. **Grave Identifier:** Burial 205, ind. 1 **Grave Type:** Trapezoidal simple pit with a covering of rectangular flat sandstone slabs resting on a wider and rectangular upper pit. It has fragments of bricks on the north side. The lower rectangular pit has rounded shorter sides. Double burial, with individuals in a supine position. Orientation is E-W. **Skeletal information:** Adult, 35-45 years old, Female. **Grave goods:** no grave goods. **Dating:** Archeologically dated to 300 – 400 CE. **Ancestry summary:** Iberia_IA__+__WestAnatolia_Roman_Byzantine__+__North_African_Punic.

- **Genetic Identifier:** I37210. **Grave Identifier:** Burial 104, ind. 5 (infant) **Grave Type:** Simple oval pit with multiple burials, an individual in a supine position without a skull, with the ossuary to the east, at the feet. Orientation is E-W. **Skeletal information:** Child, <12 years old, Undetermined sex. **Grave goods:** no grave goods. **Dating:** Archeologically dated to 300 – 400 CE. **Ancestry summary:** Italy_Sicily_Punic_Roman__+__North_African_Punic.

- **Genetic Identifier:** I37211. **Grave Identifier:** Burial 160, ind 1 **Grave Type:** A square simple pit, which is actually an ossuary, without orientation. **Skeletal information:** Adult, 35-45 years old, Undetermined sex. **Grave goods:** Ritual ceramic jar. **Dating:** Archeologically dated to 300 – 400 CE. **Ancestry summary:** CNE_EarlyMedieval.

- **Genetic Identifier:** I37212. **Grave Identifier:** Burial 223, ind. 1 **Grave Type:** A simple rectangular pit with an individual in supine position, oriented E–W. **Skeletal information:** Undetermined age and sex. **Grave goods:** no grave goods. **Dating:** Archeologically dated to 300 – 400 CE. **Ancestry summary:** Iberia_IA__+__WestAnatolia_Roman_Byzantine__+__North_African_Punic.

- **Genetic Identifier:** I37213. **Grave Identifier:** Burial 205, ind. 2 **Grave Type:** A simple trapezoidal pit with oval-shaped shorter sides, covered with well-squared sandstone slabs, supported by rubble in the north side. Double burial, individuals in supine position. Orientation E to W. **Skeletal information:** Adult, Female. **Grave goods:** no grave goods. **Dating:** Archeologically dated to 300 – 400 CE. **Ancestry summary:** Iberia_IA__+__WestAnatolia_Roman_Byzantine__+__North_African_Punic. **Additional information:** Possible osteomyelitis

- **Genetic Identifier:** I37250. **Grave Identifier:** Burial 33, ind. 1 **Grave Type:** Simple oval pit with a stone at the east end covering the feet. Individual buried in a supine position. Orientation E-W. **Skeletal information:** Child 9-10 years old, undetermined sex. **Grave goods:** no grave goods. **Dating:** Archeologically dated to 300 – 400 CE. **Ancestry summary:** Iberia_IA__+__WestAnatolia_Roman_Byzantine__+__North_African_Punic.

- **Genetic Identifier:** I37251. **Grave Identifier:** Burial 102, ind. 1 **Grave Type:** An oval simple pit, covered with rectangular sandstone slabs. Double burial, individual in supine position, with the second individual's ossuary at the feet. Orientation E-W. **Skeletal information:** Adult, >35 years old, Undetermined sex. **Grave goods:** no grave goods. **Dating:** Archeologically dated to 300 – 400 CE. **Ancestry summary:** Excluded of the Analysis due to low coverage.

- **Genetic Identifier:** I37252. **Grave Identifier:** Burial 226, ind. 1 **Grave Type:** An oval simple pit, covered with remains of tegulae and vertical brick walls. Rectangular sandstone slabs. Burial in supine position. Orientation from E-W. **Skeletal information:** Young Adult, 17-25 years old, Undetermined sex. **Grave goods:** no grave goods. **Dating:** Archeologically dated to 300 – 400 CE. **Ancestry summary:** Excluded of the Analysis due to low coverage.

- **Genetic Identifier:** I37253. **Grave Identifier:** Burial 25, in 2 **Grave Type:** Rectangular simple pit. Double burial, individuals in supine position. Orientation from east to west. **Skeletal information:** Adult, 30-40 years old, Male. **Grave goods:** no grave goods. **Dating:** Archeologically dated to 300 – 400 CE. **Ancestry summary:** Iberia_IA__+__WestAnatolia_Roman_Byzantine__+__North_African_Punic.

- **Genetic Identifier:** I37254. **Grave Identifier:** Burial 89, ind. 1 **Grave Type:** A rectangular simple pit with remains of tegulae covering. Individual burial in supine position. Orientation from northeast to southwest. **Skeletal information:** Adult, >45 years old, Female**. Grave** goods: no grave goods. **Dating:** Archeologically dated to 300 – 400 CE. **Ancestry summary:** Iberia_IA__+__WestAnatolia_Roman_Byzantine__+__North_African_Punic.

- **Genetic Identifier:** I37255. **Grave Identifier:** Burial 166, Ind. 1 **Grave Type:** A trapezoidal simple pit, covered with 4 well-squared rectangular sandstone slabs. Individual burial in supine position. Orientation from E-W. **Skeletal information:** Child, 8-10 years old, undetermined sex. **Grave goods:** 1 jug with red paint on the upper curve with a grid pattern and 2 bronze earrings. **Dating:** Archeologically dated to 300 – 400 CE. **Ancestry summary:** Iberia_IA__+__WestAnatolia_Roman_Byzantine__+__North_African_Punic.

- **Genetic Identifier:** I37256. **Grave Identifier:** Burial 94, ind. 2 (juvenile) **Grave Type:** A simple rectangular pit. Double burial, with the individual in supine position, and the second individual's ossuary at the feet. Orientation from E-W. **Skeletal information:** Child, 3-4 years old, Female. **Grave goods:** no grave goods. **Dating:** Archeologically dated to 300 – 400 CE. **Ancestry summary:** Iberia_IA__+__WestAnatolia_Roman_Byzantine__+__North_African_Punic. **Additional Information:** Shovel-shaped incisors

- **Genetic Identifier:** I37257. **Grave Identifier:** Burial 75, ind. 1 **Grave Type:** An oval-shaped simple pit, covered with four rectangular sandstone blocks. Individual buried in a supine position. Orientation from east to west. Head positioned to the west. **Skeletal information:** Child, 4-6 years old, undetermined sex. **Grave goods:** no grave goods. **Dating:** Archeologically dated to 300 – 400 CE. **Ancestry summary:** Iberia_IA__+__WestAnatolia_Roman_Byzantine__+__North_African_Punic.

##### 1.6 Girona, Pla de l'Horta

Contact person(s): Ana Costa, David Vivó and Neus Coromina

**Brief settlement history:** The location of the two necropolises arose from mechanical prospecting that followed, between 1970 and 1972, the discovery of the Roman villa. The progress of the works has made it possible to ensure that the two necropolises located were next to the last constructions of the great rural building (Agustí & Llinàs, 2012).

**Excavation History:** Here were carried three archaeological campaigns between 2004 and 2006 out on the corner of Avenida Jacint Verdaguer and Carrer de Sant Ferriol, located just 100 meters north of the excavated core of the Roman villa (Agustí & Llinàs, 2012).

**Geographic Information:** The archaeological site is located in Sarrià de Ter, 4.5 km from the current city of Girona. The necropolis is sheltered against the eastern slope of the Montagut hill and faces the small plain that opens to the west, occupying a gently sloping land that extends to the ford where one of the branches of the Via Augusta crossed the river Ter (Agustí & Llinàs, 2012).

**Summary of sampled materials:** We sampled human remains from most of the from Visigoth graves.

**Graves:**

- **Genetic Identifier:** I12030_enhanced. **Grave Identifier:** PH'06-1169 E58. **Grave Type:** Stone slab tomb. **Skeletal information:** Female, adult mature. **Grave goods:** No Grave goods. **Dating:** Archeologically dated to 500-600 CE. **Ancestry summary:** Iberia_IA__+__WestAnatolia_Roman_Byzantine__+__North_African_Punic. **Additional information:** published Olalde et al 2019.

- **Genetic Identifier:** I12031. **Grave Identifier:** PH'06-1172 E59. **Grave Type:** Simple pit with stones and tegulae**.** **Skeletal information:** Male, adult mature. **Grave goods:** two bronze buckle, a bronze square piece and an iron plate. **Dating:** Archeologically dated to 416-700 CE. **Ancestry summary:** CNE_EarlyMedieval__+__WestAnatolia_Roman_Byzantine__+__North_African_Punic. **Additional information:** published Olalde et al 2019.

- **Genetic Identifier:** I12032_enhanced. **Grave Identifier:** PH'06-1183 E63. **Grave Type:** Simple pit. **Skeletal information:** Male, adult young. **Grave goods:** two bottoms. **Dating:** Direct Radiocarbon date 420-556 calCE (1580±30BP; Beta-608905). **Ancestry summary:** CNE_EarlyMedieval__+__WestAnatolia_Roman_Byzantine__+__North_African_Punic. **Additional information:** published Olalde et al 2019.

- **Genetic Identifier:** I12033_enhanced. **Grave Identifier:** PH'06-1192 E66. **Grave Type:** Simple pit. **Skeletal information:** Male, adult young. **Grave goods:** two bronze buckle, a bronze ring, two iron knife and an iron punch **Dating:** Archeologically dated to 500-700 CE **Ancestry summary:** Iberia_IA__+__CNE_Early_medieval. **Additional information:** published Olalde et al 2019.

- **Genetic Identifier:** I12164_enhanced. **Grave Identifier:** NM 11; NPH; E54-1157. **Grave Type:** Simple pit. **Skeletal information:** Female, adult mature**.** **Grave goods:** A bronze buckle with glass, bronze fibula, a ring, a punch, an iron buckle and glass beads. **Dating:** Direct Radiocarbon date 330-433 calCE (1650 ± 30BP; Beta –541185) **Ancestry summary:** Excluded of the analysis due to low coverage. **Additional information:** published Olalde et al 2019

- **Genetic Identifier:** I13565. **Grave Identifier:** PH'06-1110 E38. **Grave Type:** Simple pit. **Skeletal information:** Female, adult young. **Grave goods:** A bronze buckle. **Dating:** Archeologically dated to 400-700 CE. **Ancestry summary:** Iberia_IA__+__WestAnatolia_Roman_Byzantine__+__North_African_Punic.

- **Genetic Identifier:** I13571. **Grave Identifier:** PH'06-1119 E41. **Grave Type:** Simple pit. **Skeletal information:** Male, adult mature. **Grave goods:** A bronze buckle. **Dating:** Archeologically dated to 400-700 CE. **Ancestry summary:** Iberia_IA__+__WestAnatolia_Roman_Byzantine.

- **Genetic Identifier:** I13572. **Grave Identifier:** PH'06-1136 E47. **Grave Type:** Stone slab tomb**. Skeletal information:** Female, adult mature. **Grave goods:** Two bronze bucke. **Dating:** Archeologically dated to 400-700 CE. **Ancestry summary: I**beria_IA__+__CNE_EarlyMedieval__+__WestAnatolia_Roman_Byzantine__+__North_ African_Punic.

- **Genetic Identifier:** I13573. **Grave Identifier:** PH'06-1086 E30. **Grave Type:** Simple pit with stones and tegulae. **Skeletal information:** Adult mature, sex inteterminated. **Grave goods:** No Grave goods. **Dating:** Archeologically dated to 500-700 CE. **Ancestry summary:** Iberia_IA__+__WestAnatolia_Roman_Byzantine.

- **Genetic Identifier:** I13574. **Grave Identifier:** PH'06-1042 E17. **Grave Type:** Simple pit with tegulae. **Skeletal information:** Female, adult young. **Grave goods:** A broze fibula, an iron ring and glass beads. **Dating:** Archeologically dated to 500-700 CE. **Ancestry summary:** Iberia_IA__+__WestAnatolia_Roman_Byzantine.

- **Genetic Identifier:** I13575. **Grave Identifier:** PH'06-1107 E37. **Grave Type:** Stone slab tomb**. Skeletal information:** Male, adult mature. **Grave goods:** A bronze buckle. **Dating:** Archeologically dated to 400-700 CE. **Ancestry summary:** Iberia_IA__+__WestAnatolia_Roman_Byzantine__+__North_African_Punic.

- **Genetic Identifier:** I20267. **Grave Identifier:** NM 4b; NPH; E7-1020. **Grave Type:** Rehused sarcophagus**. Skeletal information:** Female, adult indeterminated. **Grave goods:** No Grave goods. **Dating:** Direct InterCal20 radiocarbon dating 571-651 calCE (1450±30 BP, Beta-538218). **Ancestry summary:** Iberia_IA__+__CNE_Early_medieval__+__ North_African_Punic. **Additional information:** Tomb inside a possible mausoleum.

- **Genetic Identifier:** I20268. **Grave Identifier:** NM 15; NPH; E70-1204. **Grave Type:** Stone slab tomb. **Skeletal information:** Adult mature, sex inteterminated. **Grave goods:** No Grave goods. **Dating:** Archeologically dated to 400-700 CE. **Ancestry summary:** Iberia_IA__+__WestAnatolia_Roman_Byzantine__+__North_African_Punic.

- **Genetic Identifier:** I20269. **Grave Identifier:** NM 16; NPH; E35-1101. **Grave Type:** Simple pit. **Skeletal information:** Adult young, sex inteterminated. **Grave goods:** Some indeterminate iron pieces. **Dating:** Archeologically dated to 400-700 CE. **Ancestry summary:** Excluded from the whole-genome analysis due to contamination. **Additional information:** Revomed individual.

- **Genetic Identifier:** I20270. **Grave Identifier:** NM 17; NPH; E32-1092. **Grave Type:** Simple pit. **Skeletal information:** Adult young, sex inteterminated. **Grave goods:** Two bronze buckles, an iron knife, an some indeterminate iron pieces. **Dating:** Archeologically dated to 400-700 CE**. Ancestry summary:** CNE_EarlyMedieval__+__WestAnatolia_Roman_Byzantine.

- **Genetic Identifier:** I20271. **Grave Identifier:** NM 18; NPH; E29-1083. **Grave Type:** Simple pit with tegulae. **Skeletal information:** Male, adult mature. **Grave goods:** A bronze buckle, an iron buckle, 1 bottom and an iron needle. **Dating:** Archeologically dated to 400-700 CE. **Ancestry summary:** CNE_EarlyMedieval__+__ WestAnatolia_Roman_Byzantine__+__North_African_Punic.

- **Genetic Identifier:** I20273_d. **Grave Identifier:** NM 20; NPH; E53-1154. **Grave Type:** Simple pit. **Skeletal information:** Male, adult mature**. Grave goods:** A bronze buckle, an iron buckle. **Dating:** Archeologically dated to 400-700 CE. **Ancestry summary:** CNE_EarlyMedieval__+__WestAnatolia_Roman_Byzantine. **Additional information:** Radius pathology.

- **Genetic Identifier:** I20384. **Grave Identifier:** NM 3; NPH; E18-1045. **Grave Type:** Simple pit. **Skeletal information:** Adult young, sex inteterminated. **Grave goods:** A bronze buckle, a bronze piece, two iron knife, two nails, and few AE-4 coins. **Dating:** Direct IntCal20 radiocarbon dating to 435-587calCE (1545±20 BP, PSUAMS-10248)**. Ancestry summary:** Iberia_IA__+__WestAnatolia_Roman_Byzantine.

- **Genetic Identifier:** I20385. **Grave Identifier:** NM 5b; NPH; E7-1018. **Grave Type:** Rehused sarcophagus. **Skeletal information:** Male, adult young. **Grave goods:** No Grave goods. **Dating:** Direct IntCal20 radiocarbon dating to 605-772 cal CE (1370±30 BP). **Ancestry summary:** Iberia_IA__+__WestAnatolia_Roman_Byzantine__+__ North_African_Punic.

- **Genetic Identifier:** I20386. **Grave Identifier:** NM 6; NPH; E46-1133. **Grave Type:** Stone slab tomb. **Skeletal information:** Male, adult young. **Grave goods:** No Grave goods**. Dating:** Archeologically dated to 400-700 CE. **Ancestry summary:** Iberia_IA__+__WestAnatolia_Roman_Byzantine. **Additional information:** Possible reuse of the tomb.

- **Genetic Identifier:** I20388. **Grave Identifier:** NM 8; NPH; E20-1056. **Grave Type:** Simple pit with tegulae. **Skeletal information:** Adult, sex inteterminated. **Grave goods:** No Grave goods. **Dating:** Archeologically dated to 416-700 CE. **Ancestry summary:** Iberia_IA__+__WestAnatolia_Roman_Byzantine__+__North_African_Punic.

- **Genetic Identifier:** I20389. **Grave Identifier:** NM 9; NPH; E27-1078. **Grave Type:** Simple pit with tegulae. **Skeletal information:** Female, adult young. **Grave goods:** No Grave goods.**D ating:** Archeologically dated to 200-450 CE. **Ancestry summary:** Iberia_IA. **Additional information:** Associated to roman chronology.

- **Genetic Identifier:** I20390. **Grave Identifier:** NM 10b; NPH; E23-1075. **Grave Type:** Tegulae tomb. **Skeletal information**: Male, adult young. **Grave goods:** Pottery vase. **Dating:** Direct IntCal20 radiocarbon dating to 254-419 calCE (1700±30 BP; Beta-538220). **Ancestry summary:** Iberia_IA__+__WestAnatolia_Roman_Byzantine. **Additional information:** Associated to roman chronology.

- **Genetic Identifier:** I20391. **Grave Identifier:** NM 12; NPH; E11-1025b. **Grave Type:** Simple pit, multiple burial. **Skeletal information:** Male, adult young. **Grave goods:** No Grave goods. **Dating:** Archeologically dated to 200-450 CE. **Ancestry summary:** Iberia_IA__+__WestAnatolia_Roman_Byzantine__+__North_African_Punic. **Additional information:** Associated to roman chronology.

- **Genetic Identifier:** I20392. **Grave Identifier:** NM 13; NPH; E11-1025. **Grave Type:** Simple pit, multiple burial. **Skeletal information:** Male, adult senil. **Grave goods:** No Grave goods. **Dating:** Archeologically dated to 200-450 CE. **Ancestry summary:** Iberia_IA. **Additional information:** Associated to roman chronology.

- **Genetic Identifier:** I20393. **Grave Identifier:** NM 14; NPH; E16-1039. **Grave Type:** Simple pit and stone slab covered. **Skeletal information:** Female, adult mature. **Grave goods:** No Grave goods. **Dating:** Archeologically dated to 200-450 CE **Ancestry summary:** Iberia_IA__+__WestAnatolia_Roman_Byzantine__+__North_African_Punic. **Additional information:** Associated to roman chronology.

- **Genetic Identifier:** I20394. **Grave Identifier:** NM 21; NPH; E52-1151. **Grave Type:** Tegulae tomb. **Skeletal information:** Male, adult mature. **Grave goods:** A bronze buckle**.** **Dating** 542-605 calCE (1505±20 BP, PSUAMS-10249). **Ancestry summary:** Iberia_IA__+__WestAnatolia_Roman_Byzantine__+__North_African_Punic.

- **Genetic Identifier:** I20395. **Grave Identifier:** NM 22; NPH; E51-1147. **Grave Type:** Stone slab tomb**. Skeletal information:** Juvenile, sex inteterminated**. Grave goods:** An iron buckle. **Dating:** Archeologically dated to 400-700 CE. **Ancestry summary:** Excluded from the genome wide analysis due to low coverage.

- **Genetic Identifier:** I20396. **Grave Identifier:** NM 23; NPH; E67-1195. **Grave Type:** Simple pit. **Skeletal information:** Female, adult mature. **Grave goods:** No Grave goods. **Dating:** Archeologically dated to 416-700 CE. **Ancestry summary:** CNE_EarlyMedieval__+__WestAnatolia_Roman_Byzantine__+__North_African_Punic.

- **Genetic Identifier:** I20397. **Grave Identifier:** NM 24; NPH; E65-1189. **Grave Type:** Simple pit. **Skeletal information:** Female, adult senile. **Grave goods:** No Grave goods. **Dating:** Archeologically dated to 400-700 CE. **Ancestry summary:** CNE_EarlyMedieval__+__WestAnatolia_Roman_Byzantine__+__North_African_Punic.

- **Genetic Identifier:** I20398. **Grave Identifier:** NM 25; NPH; E45-1130. **Grave Type:** Simple pit. **Skeletal information:** Male, adult mature. **Grave goods:** An iron buckle. **Dating:** Archeologically dated to 400-700 CE. **Ancestry summary:** Excluded from the genome wide analysis due to low coverage.

- **Genetic Identifier:** I20400. **Grave Identifier:** NM 27; NPH; E43-1124. **Grave Type:** Simple pit. **Skeletal information:** Male, adult mature. **Grave goods: No Grave goods.Dating:** Archeologically dated to 400-700 CE. **Ancestry summary:** Iberia_IA__+__CNE_EarlyMedieval__+__WestAnatolia_Roman_Byzantine__+__North_African_Punic.

- **Genetic Identifier:** I20401. **Grave Identifier:** NM 28; NPH; E28-1076. **Grave Type:** Stone slab tomb**. Skeletal information:** Adult mature, sex inteterminated. **Grave goods:** No Grave goods. **Dating:** Archeologically dated to 400-700 CE. **Ancestry summary: SpainIA__+__**CNE_EarlyMedieval__+__WestAnatolia_Roman_Byzantine__+__North_African_Punic.

- **Genetic Identifier:** I20402. **Grave Identifier:** NM 29; NPH; E55-1160. **Grave Type:** Stone slab tomb**. Skeletal information:** Male, adult mature. **Grave goods:** No Grave goods. **Dating:** Archeologically dated to 400-700 CE. **Ancestry summary:** Iberia_IA__+__WestAnatolia_Roman_Byzantine__+__North_African_Punic.

##### 1.7 Girona, Sant Julià de Ramis, Les Goges

<u>Contact person(s):</u> Josep Burch, Jordi Sagrera, Jordi Vivo & Neus Coromina

**Brief settlement history:** The Goges necropolis is an intensive burial ground without an habitat site/village associated. There is litte información about the procedence from the people buried in that cementery (Agustí & Llinàs, 2012).

**Excavation History:** The excavations at the Les Goges necropolis took place in 1991 because public construction projects affected the site. After the excavation, the site was destroyed (Agustí & Llinàs, 2012).

**Geographic Information:** The Les Goges necropolis is located in the northeastern corner of the Iberian Peninsula. Specifically, it is on a terrace beside the Ter River, about 5 km east of the city of Girona (Agustí & Llinàs, 2012).

**Summary of sampled materials:** This is a burial necropolis. During the excavations, 207 graves were documented, most of them in simple pits. Very few of these graves contained burial goods. The timeline is from the early 5th century to the early 8th century (Agustí & Llinàs, 2012).

**Graves:**

- **Genetic Identifier:** I20274. **Grave Identifier:** NM 32; LG'91; E81. **Grave Type:** Simple pit. **Skeletal information:** Female, adult mature. **Grave goods:** No Grave goods. **Dating:** Direct IntCal20 Radiocarbon date to 426-575 calCE (1560±30 BP; Beta-507251). **Ancestry summary:** WestAnatolia_Roman_Byzantine__+__North_African_Punic.

- **Genetic Identifier:** I20275. **Grave Identifier:** NM 34; LG'91; E23. **Grave Type:** Simple pit. **Skeletal information:** Female, adult mature. **Grave goods:** No Grave goods. **Dating:** Direct IntCal20 Radiocarbon date to 607-774 calCE (1360±30 BP; Beta-507250). **Ancestry summary:** WestAnatolia_Roman_Byzantine.

- **Genetic Identifier:** I20276. **Grave Identifier:** NM 36; LG'91; E193. **Grave Type:** Simple pit. **Skeletal information:** Female, adult. **Grave goods:** No Grave goods. **Dating:** Direct IntCal20 Radiocarbon date to 607-774 calCE (1360±30 BP; Beta-507253). **Ancestry summary:** Iberia_IA__+__WestAnatolia_Roman_Byzantine__+__North_African_Punic.

- **Genetic Identifier:** I20278. **Grave Identifier:** NM 39; LG'91; E165. **Grave Type:** Simple pit. **Skeletal information:** Male, adult mature. **Grave goods:** No Grave goods. **Dating:** Direct IntCal20 Radiocarbon date to 584-658 calCE (1430±30 BP; Beta-507252). **Ancestry summary:** Iberia_IA__+__WestAnatolia_Roman_Byzantine__+__North_African_Punic8.

- **Genetic Identifier:** I20405. **Grave Identifier:** NM 33; LG'91; E108. **Grave Type:** Simple pit. **Skeletal information:** Adult, sex inteterminated. **Grave goods:** Coin. **Dating:** Archeologically dated to 400-700 CE. **Ancestry summary:** Excluded from the genome wide analysis due to low coverage.

- **Genetic Identifier:** I20406. **Grave Identifier:** NM 35; LG'91; E113. **Grave Type:** Stone slab tomb**. Skeletal information:** Juvenile, sex inteterminated. **Grave goods:** No Grave goods. **Dating:** Archeologically dated to 400-700 CE. **Ancestry summary:** Excluded from the genome wide analysis due to low coverage.

- **Genetic Identifier:** I20407. **Grave Identifier:** NM 38; LG'91; E164. **Grave Type:** Simple pit. **Skeletal information:** Adult, sex inteterminated.. **Grave goods:** No Grave goods. **Dating:** Archeologically dated to 400-700 CE. **Ancestry summary:** WestAnatolia_Roman_Byzantine__+__North_African_Punic.

- **Genetic Identifier:** I20408. **Grave Identifier:** NM 40; LG'91; E26. **Grave Type:** Simple pit. **Skeletal information:** Adult mature, sex inteterminated.**Grave goods:** No Grave goods. **Dating:** Archeologically dated to 400-700 CE. **Ancestry summary:** Iberia_IA__+__WestAnatolia_Roman_Byzantine__+__North_African_Punic.

##### 1.8 Granada, Colomera, Cortijo del Chopo

Contact person(s): Juan Manuel Jiménez Arenas, Elena Vallejo-Casas, Paula Chirosa Cañavate

**Brief settlement history:** The discovery of the necropolis took place in the late 1980s, when several of the funerary structures were uncovered and began to be looted. This activity, which endangered the integrity of the remains, led to an emergency archaeological intervention. The constant pillaging and the need to understand the dimensions of the late antique cemetery and its characteristics in greater detail, led to the subsequent continuation of excavation works at the site. A total of 168 graves were documented, most of them pit burials covered with slabs. These are arranged and orientated parallel to the slope of the terrain on which they are placed, surrounding it. Most of the individuals identified were found in primary deposits and in a supine position, as well as in reductions located at the foot or sides of the latest deceased. Although many of the tombs were partially or completely looted, half of them contained elements of personal adornment (rings, earrings, bracelets, pins, beads and buckles) and funerary deposits (pottery fragments and ceramic vases). In order to gain an in-depth knowledge of the recorded materials, different analytical techniques such as x-ray fluorescence, lead isotopes and infrared spectroscopy were carried out on a selection of the assemblage. It has been proposed that the site of Las Mesas, located in an extensive hilly area a short distance from Cortijo del Chopo, could be the settlement place related to the population buried in the necropolis.

**Excavation History:** The Cortijo del Chopo site was the object of three archaeological interventions carried out in 1986, 1988 and 1989. The first campaign made it possible to document the looted tombs and identified a total of 47 burials. During the interventions conducted in 1988 and 1989, 121 burials were exhumed, and various clothing items and funerary deposits, such as pottery vases, were also recovered. The results of the 1986 campaign were the subject of a study and have been published (Moyano et al., 1989), while the funerary structures and materials obtained in 1988 and 1989 were neither analysed nor made public. The bone remains recorded in 118 of the 168 graves excavated were not studied. The investigation of a selection of the anthropological collection, as well as of the entire assemblage of pieces, is a key focus of E. Vallejo-Casas's ongoing PhD thesis.

**Geographic Information:** The necropolis is located in the town of Colomera, in the province of Granada. Its UTM ETRS89 coordinates are 30T X: 437686 Y: 4138013. The site is situated on a rocky escarpment that interrupts the steep slope down to the Colomera River (Moyano et al., 1989). The tombs are arranged in the terrain between approximately 879 to 886 metres above sea level. To the southeast of the necropolis, the Mingarrón stream runs close to the foot of the Sierra de los Hornos, and to the southwest are Los Peñones and Las Mesas hill. The area also has several perennial and seasonal springs. The resources that sustained the economy of the community that inhabited the area would have been related to livestock, forests, and dry farming. It should be noted that Cortijo del Chopo is placed in the Colomera valley, which is on an important communication axis that connects Granada with the northern territory towards Jaén (Mattei, 2010).

**Summary of sampled materials:** After the interventions in the necropolis, the bone remains were deposited in the Archaeological and Ethnological Museum of Granada without undergoing any analysis. Currently, an anthropological study of the unpublished collection is being carried out gradually. As part of the research, radiocarbon analysis was conducted on five individuals from four prominent burials, whose results allow us to date them to between the 6th and early 8th centuries AD. Samples for ancient DNA analysis were taken from a smaller set of individuals studied, selected based on the characteristics of the materials they were associated with and because no signs of looting were found during their excavation.

**Graves:**

- **Genetic Identifier:** I32150. **Grave Identifier:** NCH-86, CI; T: XXXII (32) [3.200] (1). **Grave Type:** Pit burial covered with slabs. Individual in primary supine position. NNW-SSW orientation. (NMI in the grave: 6). **Skeletal information:** Infantile-juvenile, undetermined sex. **Grave goods:** bead and pottery vase**. Dating:** Archeologically dated to 500-700 CE. **Ancestry summary:** Iberia_IA__+__WestAnatolia_Roman_Byzantine__+__North_African_Punic.

- **Genetic Identifier:** I32152. **Grave Identifier:** NCH-86, CI; T: XXXIII (33) P:17. **Grave Type:** Pit burial covered with slabs. Individual in primary supine position, NNW-SSW orientation. (NMI in the grave: 1). **Skeletal information:** Infant, undetermined sex. **Grave goods:** earrings, beads **Dating:** Archeologically dated to 500-700 CE. **Ancestry summary:** Iberia_IA__+__WestAnatolia_Roman_Byzantine__+__North_African_Punic.

- **Genetic Identifier:** I32154. **Grave Identifier:** NCH-86, CI; T: XXXIV (34) Individuo 1 en posición, pieza No. 16 (3.400). (NMI in the grave: 3). **Grave Type:** Pit burial covered with slabs. Individual in primary supine position, NNW-SSW orientation. **Skeletal information:** Adult, undetermined sex. **Grave goods:** Bracelet, pottery vase. **Dating:** Archeologically dated to 500-700 CE. **Ancestry summary:** Iberia_IA__+__WestAnatolia_Roman_Byzantine__+__North_African_Punic.

- **Genetic Identifier:** I32155. **Grave Identifier:** NCH-86, CI; T34 XXXIV, Mandibula (2), (3.413), P:37. (NMI in the grave: 3). **Grave Type:** Pit burial covered with slabs. Individual reduced at the feet of the latest deceased, NNW-SSW orientation. **Skeletal information:** Adult, undetermined sex. **Grave goods:** Bracelet, pottery vase**. Dating:** Archeologically dated to 500-700 CE. **Ancestry summary:** Iberia_IA__+__WestAnatolia_Roman_Byzantine__+__North_African_Punic.

- **Genetic Identifier:** I32156. **Grave Identifier:** NCH-86, CI; T: XXXIV (35) Individuo 2 [3.503] P:27. (NMI in the grave: 4). **Grave Type:** Pit burial covered with slabs. Individual in primary supine position (lower), NNW-SSW orientation. **Skeletal information:** Adult, undetermined sex. **Grave goods:** earrings, bracelet. **Dating:** Archeologically dated to 500-700 CE. **Ancestry summary:** Iberia_IA__+__WestAnatolia_Roman_Byzantine__+__North_African_Punic.

- **Genetic Identifier:** I32157. **Grave Identifier:** NCH-86, CI; T: XXXV (35) Craneo 3 [3.505]. (NMI in the grave: 4). **Grave Type:** ). Pit burial covered with slabs. Individual reduced at the feet of the latest deceased, NNW-SSW orientation **Skeletal information:** Adult, undetermined sex. **Grave goods:** Earrings, bracelet. **Dating:** Archeologically dated to 500-700 CE. **Ancestry summary:** Iberia_IA__+__WestAnatolia_Roman_Byzantine__+__North_African_Punic.

- **Genetic Identifier:** I32158. **Grave Identifier:** NCH-86, CI; T: XXXV (35) Craneo 4 [3.506]. (NMI in the grave: 4). **Grave Type:** Pit burial covered with slabs. Individual reduced at the feet of the latest deceased, NNW-SSW orientation. **Skeletal information:** Adult, undetermined sex. **Grave goods:** Earrings, bracelet. **Dating:** Archeologically dated to 500-700 CE. **Ancestry summary:** Iberia_IA__+__WestAnatolia_Roman_Byzantine__+__North_African_Punic.

- **Genetic Identifier:** I32160. **Grave Identifier:** NCH-86, CI; Amontonam:IV, (40.005) (3), P:36. **Grave Type:** Simple pit burial. Individual in primary supine position, NNW-SSW orientation. **Skeletal information:** Juvenile, undetermined sex. **Grave goods:** Rings, earrings, bracelets, beads. **Dating:** Archeologically dated to 500-700 CE. **Ancestry summary:** Iberia_IA__+__WestAnatolia_Roman_Byzantine__+__North_African_Punic.

- **Genetic Identifier:** I32162. **Grave Identifier:** NCH-86, CI; burial 47 (previously notated as "Amontonamiento IV), (40.009) (2), P:45. NMI in the grave: 2). **Grave Type:** Simple pit burial. Individual in position not possible to determine. NNW-SSW orientation. **Skeletal information** Juvenile, undetermined sex. **Grave goods:** rings, earrings, bracelets, beads**.** **Dating:** Archeologically dated to 500-700 CE. **Ancestry summary:** Iberia_IA__+__ North_African_Punic.

#### 1.9 Granada, Montefrío, El Castillón

Contact person(s): Juan Manuel Jiménez Arenas

**Brief settlement history:** The necropolis of El Castillón (Montefrío, Granada) has been known since the second half of the 19th century. It was documented in Manuel de Góngora's book "Antigüedades Prehistóricas de Andalucía" (1868). Subsequently, it was cited by Manuel Gómez Moreno and Miguel Tarradell. All the information gathered by these scholars points to a Visigothic context for this necropolis. It was not until 1977 that six excavations, alternating necropolis with settlement, unearthing several domestic building structures, and 115 tombs (1977, 1980, 1983). However, despite the importance of this site, publications about it are exceedingly rare.

**Excavation History:** During the 1977 campaign, 32 graves were excavated. The following campaign carried out in 1980 increased the number of graves by 27. From these two campaigns, there is information on numerous grave goods, of which only a part is preserved in the Archaeological Museum of Granada. The 1983 campaign added 54 new graves (9 looted by clandestine excavators and the rest containing only skeletal remains).

All the tombs follow the same general construction process. They consist of a pit dug into the geological substrate, with the walls lined with vertical stone slabs and covered with horizontal stone slabs as a roof.

The tombs of El Castillón are organized in more or less parallel rows, or forming streets. The orientation of the graves is N-S, with some variations following the orography. Within the necropolis, a wall has been located in the southern part, near a tomb whose function is unknown. It may have served to divide the necropolis into two sectors or been part of a non-preserved structure.

Regarding the burial ritual, it is inhumation, with the graves being individual, double, or triple, and sometimes reused. Infant tombs were also documented. During the old excavations, it was noted that many of them were sterile (Torres, 1977, 1981). However, during the consolidation and restoration work of the necropolis between 2001-2002, 72 graves contained human remains (Afonso & Ramos, 2005). Additionally, two of the tombs (VII and XV) show evidence of fire inside.

The grave goods found in some tombs of the necropolis, especially the metal items and some ceramic jars, as well as the typology of the graves, point to the 6th and 7th centuries CE, coinciding partly with the Byzantine occupation of this part of the Iberian Peninsula.

**Geographic Information:** The necropolis is located on the hill of Castillón (Montefrío, Granada), occupying the gentle western slope of it (37°20'01.9"N, -3°58'32.2"W), very close to the Late Antiquity settlement, and relatively near emblematic sites of the Recent Prehistory of the southern Iberian Peninsula (Los Castillejos de la Peña de los Gitanos).

**Summary of sampled materials:** The grave goods are notable for the presence of ceramic jars, some of which include simple geometric decorations (parallel lines, comb patterns, zigzags, etc.), and metal adornments. Among these, earrings stand out, decorated with lines or dots or left plain, made of bronze. One of them features a pendant in the shape of a glass paste bead. There are also bracelets, rings, pins, and a needle, all made of bronze. Particularly striking are the 106 necklace beads made of jet, glass paste, and clay found in tomb LI. Finally, it is important to mention the 4 oval buckles with bronze pins and, most notably, the unique belt buckle found in tomb XIX, which features two rampant animal figures in the center of the buckle, holding a cup with green almond-shaped decorations that, according to the discoverers, would be a Byzantine motif. Sample re-sequenced from (Olalde et al., 2019).

**Graves:**

- **Genetic Identifier:** I3583_enhanced. **Grave Identifier:** Sepultura LXXX. **Grave Type:** Pit dug into the geological substrate, with the walls lined with vertical stone slabs and covered with horizontal stone slabs as a roof. **Skeletal information:** Undetermined sex and age. **Grave goods:** Absent. **Dating:** Archeologically dated to 400-600 CE. **Ancestry summary:** Iberia_IA__+__WestAnatolia_Roman_Byzantine__+__North_African_Punic. **Additional information:** Intact tomb.

##### 1.10 Granada, Plaza Einstein

Contact person(s): Juan Manuel Jiménez Arenas

**Brief settlement history:** The archaeological activity consisted of monitoring the earth movements associated with the construction project of Line 1 of the Granada Light Metro. This was done so that, in the event of detecting sedimentary and/or structural evidence of archaeological significance during the activity, these could be delimited and subsequently recovered and documented using manual methods in accordance with archaeological methodology. Previously, a Roman villa from the early Imperial period was documented in this area.Once it was abandoned, collapsed, and silted up, this space was used as a necropolis from the 4th century CE onwards.

**Excavation History:** The discovery of grave SEP-049 occurred within the framework of preventive archaeological interventions carried out during the construction of Line 1 of the Granada Light Metro, in its Camino de Ronda section. A series of structures and archaeological evidence were found, indicating an occupation of the peri-urban area of Granada from the 2$^{nd}$ and 3$^{rd}$ centuries CE to the 6$^{th}$ and 7$^{th}$ centuries CE, with notable remains associated with the city's Paleochristian period.

The excavation was carried out "in mine" focusing on the building structures and the graves that were detected during the monitoring of the earth movements. Levels have been documented from the 3$^{rd}$ century CE to the 6$^{th}$ century CE. Among the structures, the so-called Building I stands out, as well as the so-called Paleochristian structural complex (4th-6th centuries CE), which was reused as a cemetery area in a residual manner. Additionally, two Late Roman structural complexes dated by the presence of ceramics and a coin (3rd-4th centuries CE) and a structural complex from the High Imperial period (2nd-3rd centuries CE) were identified.

The SEP-049 grave was exhumed during the excavation of one of the accesses to an underground parking lot. It was found with a high degree of destruction due to constructions carried out in the present era, and its length was recorded as 1.40 m, a width of 0.90 m, and a maximum height of 0.55 m. Next to it, another grave (SEP-050) with a gabled roof and W-E orientation was documented (though not excavated).

From a functional point of view, it has been suggested that the oldest phase was linked to a production area associated with a villa or an agricultural production area. Subsequently, it was used as a dump. After this period, it was abandoned and reused as a cemetery. Later, the construction of a large building (interpreted as a mausoleum or pantheon, which evolved into a place of worship in the 5th century CE) caused the reorganization of the necropolis, intensifying its use as a burial space. It was during this time (5th century CE) that another less substantial building was constructed, which could have been a Martyrium.

**Geographic Information**: This grave was located on the access ramp to the underground parking lot situated on Camino de Ronda Street, PK 1.400 – 1.440, in the urban center of the city of Granada (37°10'17.6"N, 3°36'24.1"W).

**Summary of sampled materials:** In the grave where the sampled individual was found, no grave goods were discovered that could serve for social and/or temporal characterization. Individuals re-sequenced from (Olalde et al., 2019).

**Graves:**

- **Genetic Identifier:** I4055_enhanced. **Grave Identifier:** Tumba 49. **Grave Type:** Pit excavated into the geological substrate with walls made of medium-sized sandstone blocks and covered with the same construction material, all secured with earth. MNI: 1. Orientation W-E. **Skeletal information:** Male, Adult. **Grave goods:** No grave goods. **Dating:** Archeologically dated to 200-500 CE. **Ancestry summary:** Iberia_IA__+__WestAnatolia_Roman_Byzantine__+__North_African_Punic. **Additional information:** Undisturbed grave.

##### 1.11 Granada, Zafarraya, Necropolis de Las Delicias

Contact person(s): Juan Manuel Jiménez Arenas

**Brief settlement history:** Zafarraya settlement area was inhabited by Ek Argar peoples, and evidence of Phoenician and Roman camps have been found. It acted as an important location in the crossroads of Vélez, Alhama and Loja settlements.

**Excavation History:** There have been reports of late necropolises in the area since 1871, at which time a bronze earring, two copper earrings, a copper ear clasp, and a bronze buckle with decorations, among other metal objects, were entered into the Archaeological Museum of Granada. A total of two archaeological excavations were carried out during 1985 and 1986.

**Geographic Information:** The necropolis is located within the urban core of Zafarraya town (37°11'01"N, 4°39'00"W).

**Summary of sampled materials:** A total of 28 tombs with structures and another 9 that we have termed "piles" were found during 1985 and 1986 excavations. Two of the tombs were not excavated. Regarding the typology of the burials, the most common type consists of a box or cist made by placing vertical slabs to delimit an area. Some of them had a cover, while others did not. The absence of a cover may be due to looting, agricultural activities, or damage from the excavator, which piled up numerous slabs in the western sector. The orientation of these tombs is west-east (W-E).

The two most distinctive tombs represent a new type, not documented in the first campaign, characterized by a Roman burial covered with double-sloped tegulae tiles. These are tomb XIX and XXVI. The type of tomb, burial ritual, and the grave goods found suggest a Late Antiquity timeframe for these burials. Specifically, according to the archaeologists who excavated this necropolis, the presence of handmade ceramic bowls and glass bowls decorated with horizontal striations point to a possible Byzantine. The presence of three rectangular plates of fire-gilded copper found in three tombs, decorated with zigzag patterns, also point to Byzantine influence. On the other hand, the presence of a belt buckle with glass paste inlays and featuring a swastika in another tomb denotes a Ostrogothic influence. Similarly, the presence of oval buckles with pins, kidney-shaped buckles, shield-shaped belt fittings, and glass paste beads suggest a Germanic affiliation of the individuals buried with them.

Tomb XIX is an individual burial in a pit with a heavily deteriorated double-sloped tegulae tile cover. The corpse had its head resting on a well-fired red-colored roof tile, 50 cm long. The body was in a supine position. The burial goods are described in the corresponding section. The proposed chronology for this tomb and the other covered with tegulae tiles is the 4th century CE.

Based on all the previously mentioned evidence, it has been proposed that this necropolis was in use from the 3rd (presence of a bronze coin of Decius Emperor found in tomb XXIV) to the 7th century CE.

**Graves:**

- **Genetic Identifier:** I3584_enhanced. **Grave Identifier:** Tumba XIX. **Grave Type:** Pit dug into the geological substrate with a heavily deteriorated double-sloped tegulae tile cover. NMI: 1. Orientation W-E. **Skeletal information:** Male, Adult. **Grave goods:** A very rusted iron nail and a ceramic vessel of reddish ocher color, which bears a resemblance to form 45 described by Vegas, but we lean more towards comparing it with a vessel found in the necropolis of Aldea de San Esteban (4th century). **Dating:** Archeologically dated to the $4^{th}$ century CE. **Ancestry summary:** Iberia_IA__+__WestAnatolia_Roman_Byzantine__+__North_African_Punic. **Additional information:** Intact tomb.

#### 1.12 Guadalajara, Cubillejo de la Sierra

Contact person(s): Maria Luisa Cerdeño, Miguel Ángel Cuadrado

**Brief settlement history:** After surveys of nearby lands, the only settlement site that matches the necropolis is the villa of La Vega. This villa is at the foot of the hill where the cemetery is located. It has been greatly affected by farming activities. Over the years, fragments of mosaic, two column shafts, and ceramics such as black-glazed ware, Gallic terra sigillata, Hispanic terra sigillata, and T.S.H.T. have been found. Therefore, the villa's timeline likely extends from the late Republican period to late antiquity.

**Excavation History:** The discovery of the Late Antiquity cemetery of Cubillejo de la Sierra was unexpected but has opened new perspectives in the studies about the Late Antiquity in this region of Spanish Plateau. It was unexpectedly discovered during the work started in 2006 at the Celtiberian oppidum of Los Rodiles and its immediate surroundings, where several sites from different periods are concentrated (Cerdeño et al., 2008). A total of 22 graves have been excavated, but the evidence points to a much larger cemetery with more than one hundred graves (Cerdeño et al., 2015).

**Geographic Information:** This new cemetery is located in Cubillejo de la Sierra, near Molina de Aragón. The necropolis extends over the southern spur of Loma Gorda, on its flat surface and the east side of the hermitage. The hill reaches 1140 meters above sea level and is very eroded, with many limestone outcrops. At the base flows the Arroyo de la Vega, which forms the headwaters of the Piedra River upstream.

**Summary of sampled materials:** The dead are accompanied of graves goods not very rich as befits a rural society: some fibulae, buckles, belt clips and bronze rings; small iron knives and necklace beads of different materials. The preliminary study of the findings, including the burial shapes and few grave goods, suggests they date to the 6th and 7th centuries. We cannot be more precise about the dates. The bronze objects are similar to those from other graves, like those in the necropolis of Cacera de las Ranas in Aranjuez. There, shield-shaped appliqués were dated to the 6th century, and rigid plate belt buckles to the early 7th century. Therefore, we might place the Cubillejo necropolis in phases II (530-580 AD) and III (580-640 AD) of the Visigothic necropolises in Guadalajara (Daza & Catalán, 2009).

**Graves:**

- **Genetic Identifier:** I19665. **Grave Identifier:** EV-2006; Cata 8F; T-1. **Grave Type:** Precence of nails indicate the decesed was placed in a wooden coffin. Grave surrounded by large stone slabs (cist-like). E-W orientation (head towards sunrise). **Skeletal information:** male, <50 years old. **Grave goods:** a small rectangular bronze buckle, 20x10mm, with a pin and four hemispherical rivets at the corners, two bronze shield-shaped appliqués, fragments of an iron knife blade, and other small items. **Dating:** Archeologically dated to 500-700 CE. **Ancestry summary:** CNE_EarlyMedieval__+__WestAnatolia_Roman_Byzantine__+__North_African_Punic. **Additional information:** The grave was well-preserved, though the skeleton was incomplete.

- **Genetic Identifier:** I19666. **Grave Identifier:** EV'14; Cata 8F-8E; T-2. **Grave Type:** Precence of nails indicate the decesed was placed in a wooden coffin. Grave surrounded by large stone slabs (cist-like). E-W orientation (head towards sunrise). **Skeletal information:** woman, adult, over 25 years old. **Grave goods:** a belt plate with a rectangular heel, a shield-shaped base pin, and two perforated plates. **Dating:** Archeologically dated to 500-700 CE. **Ancestry summary:** Iberia_IA__+__WestAnatolia_Roman_Byzantine__+__North_African_Punic. **Additional information:** The surface soil had been disturbed.

- **Genetic Identifier:** I19667. **Grave Identifier:** EV'14; Cata 1E; T-4. **Grave Type:** Precence of nails indicate the decesed was placed in a wooden coffin. Grave surrounded by large stone slabs (cist-like). E-W orientation (head towards sunrise). **Skeletal information:** woman, adult, aged between 31 and 59 years. **Grave goods:** a 26mm bronze oval buckle with a curved pin, a 13mm diameter bronze ring with a small hanging loop, and a cylindrical ceramic bead. **Dating:** Direct InCal 20 radiocarbon dated to 569-643 calCE (1470±20 BP, PSUAMS-9708) **Ancestry summary:** CNE_EarlyMedieval__+__WestAnatolia_Roman_Byzantine__+__North_African_Punic. **Additional information:** The foot of the grave was disturbed when grave 6 was placed above it.

- **Genetic Identifier:** I19668. **Grave Identifier:** EV'14; Cata 1E-2E; T-5. **Grave Type:** Precence of nails indicate the decesed was placed in a wooden coffin. Grave surrounded by large stone slabs (cist-like). E-W orientation (head towards sunrise). **Skeletal information:** woman, adult, aged between 35 and 45 years. **Grave goods:** a 36mm diameter iron circular buckle with a fused pin, a deformed bronze ring with open ends, and various small bronze fragments. **Dating:** Archeologically dated to 500-700 CE. **Ancestry summary:** Iberia_IA__+__WestAnatolia_Roman_Byzantine__+__North_African_Punic. **Additional information:** Some stone slabs were slightly disturbed.

- **Genetic Identifier:** I19669. **Grave Identifier:** EV'14; Cata 1E-2E; T-6. **Grave Type:** Precence of nails indicate the decesed was placed in a wooden coffin. Grave surrounded by large stone slabs (cist-like). E-W orientation (head towards sunrise). **Skeletal information:** Male, Adult, elderly. **Grave goods:** a belt buckle with a rigid plate, a curved heel, and a shield-shaped base pin with perforated plates on the back, measuring 130x45mm and 2mm thick, a 20mm diameter bronze ring and a cylindrical blue glass bead with a white zigzag pattern. **Dating:** Archeologically dated to 500-700 CE. **Ancestry summary:** Iberia_IA__+__CNE_EarlyMedieval__+__WestAnatolia_Roman_Byzantine__+__North_ African_Punic. **Additional information:** The skeleton was incomplete, with the grave superimposed over T-4.

- **Genetic Identifier:** I19670. **Grave Identifier:** EV'17; Cata 2B-2C; T-16. **Grave Type:** Precence of nails indicate the decesed was placed in a wooden coffin. Grave surrounded by large stone slabs (cist-like). E-W orientation (head towards sunrise). **Skeletal information:** Male, adult, aged between 35 and 55 years. **Grave goods:** an 88mm iron chisel and a 10mm diameter iron rod. **Dating:** Direct InCal 20 radiocarbon dated to 561-641 calCE (1480±20 BP, PSUAMS-10215). **Ancestry summary:** Iberia_IA__+__WestAnatolia_Roman_Byzantine__+__North_African_Punic. **Additional**

**information:** held children's phalanges in his right hand and a vertebra in his left hand. The stone slabs were disordered, but the skeleton was complete and well-preserved.

- **Genetic Identifier:** I19671. **Grave Identifier:** EV'2014; Cata 3-9D; T-7. **Grave Type:** Precence of nails indicate the decesed was placed in a wooden coffin. Grave surrounded by large stone slabs (cist-like). E-W orientation (head towards sunrise). **Skeletal information:** Male, adult, about 20 years old. **Grave goods:** de two fragments of a small yellow glass container. **Dating:** Archeologically dated to 500-700 CE. **Ancestry summary:** Iberia_IA__+__North_African_Punic. **Additional information:** The grave was disturbed at the surface, and the skeleton was incomplete.

- **Genetic Identifier:** I19672. **Grave Identifier:** EV'17; Cata 4E; T-13. **Grave Type:** Precence of nails indicate the decesed was placed in a wooden coffin. Grave surrounded by large stone slabs (cist-like). E-W orientation (head towards sunrise). **Skeletal information:** *Male, Adult.* **Grave goods:** radial head of a bronze fibula with five molded appendages, and both decorated; the plate measures 33mm in diameter. **Dating:** Direct InCal 20 radiocarbon dated to 646-772 calCE (1350±20 BP, PSUAMS-9709). **Ancestry summary:** Iberia_IA__+__WestAnatolia_Roman_Byzantine__+__North_African_Punic. **Additional information:** The surface was disturbed, and the skeleton was complete but very deteriorated.

- **Genetic Identifier:** I19673. **Grave Identifier:** EV'17; Cata 4C-6C; T-19. **Grave Type:** Precence of nails indicate the decesed was placed in a wooden coffin. Grave surrounded by large stone slabs (cist-like). E-W orientation (head towards sunrise). **Skeletal information:** Woman, Adult, between 16-20 years old. **Grave goods:** No grave goods. **Dating:** Archeologically dated to 500-700 CE. **Ancestry summary:** No model found. **Additional information:** The grave was well-preserved, and the woman's height was estimated at 1.59 meters.

- **Genetic Identifier:** I19674. **Grave Identifier:** EV'17; Cata 7B; T-15. **Grave Type:** Precence of nails indicate the decesed was placed in a wooden coffin. Grave surrounded by large stone slabs (cist-like). E-W orientation (head towards sunrise). **Skeletal information:** Male, adult, around 20. years old. **Grave goods:** A highly deteriorated bronze coin measuring 20 mm in diameter and 2 mm in thickness. The ring of a buckle, 7 mm thick, with a notch where the pin would be inserted. A mini bronze buckle with an oval ring and a pin thickened at the tip. A bronze shield-shaped applique with a central rib, measuring 23 mm in length and 3 mm in thickness. A bronze washer with a diameter of 27 mm. A bronze rod with a preserved length of 61 mm. The distal bronze fitting of a knife sheath, with a preserved length of 20 mm. An iron knife, heavily oxidized, with a straight back and flat tang, measuring 140 mm in preserved length and 19 mm in blade width. Another iron knife, heavily oxidized, with a straight back and slightly curved edge, measuring 130 mm in preserved length and 20 mm in blade width. **Dating:** Archeologically dated to 500-700 CE. **Ancestry summary:** Excluded of the genome wide analysis due to coverage. **Additional information:** The grave was disturbed at the surface, and the skeleton was incomplete.

- **Genetic Identifier:** I19675. **Grave Identifier:** EV'17; Cata 12C; T-18a. **Grave Type:** Precence of nails indicate the decesed was placed in a wooden coffin. Grave surrounded by large stone slabs (cist-like). E-W orientation (head towards sunrise). **Skeletal information:** Woman, adult, between 18-20 years old. **Grave goods:** All grave goods were placed on the lower skeleton (T-18b / I19676). **Dating:** Direct InCal 20 radiocarbon dated to 594-650 calCE (1440±20 BP, PSUAMS-9710). **Ancestry summary:** Iberia_IA__+__WestAnatolia_Roman_Byzantine__+__North_African_Punic. **Additional information:** The upper skeleton is from a double burial, either superimposed or simultaneous.

- **Genetic Identifier:** I19676. **Grave Identifier:** EV'17; Cata 12C; T-18b. **Grave Type:** Precence of nails indicate the decesed was placed in a wooden coffin. Grave surrounded by large stone slabs (cist-like). E-W orientation (head towards sunrise). **Skeletal information:** Woman, adult, between 18-20 years old **Grave goods:** a fragment of a ceramic jug with a thickness of 6 mm. A radial-headed fibula with 5 appendages, a short bridge, and a long foot, with fabric attached, measuring 84 mm in length and 15 mm in height. A bronze earring with a barrel-shaped pendant, measuring 25 mm in diameter. 200 necklace beads made of various materials and shapes (bronze, Baltic amber, black paste, blue paste, green paste). **Dating:** Archeologically dated to 500-700 CE. **Ancestry summary:** Iberia_IA__+__WestAnatolia_Roman_Byzantine__+__North_African_Punic. **Additional information:** This is the lower skeleton from the previously described double burial.

- **Genetic Identifier:** I19677. **Grave Identifier:** EV'17; Cata 12C; T-18b. **Grave Type:** Precence of nails indicate the decesed was placed in a wooden coffin. Grave surrounded by large stone slabs (cist-like). E-W orientation (head towards sunrise). **Skeletal information:** Male, very incomplete, undetermined age. **Grave goods:** Two identical bronze buttons with conical heads decorated with incised petal-like lines and perforated tabs for attachment, measuring 13 mm in diameter and 15 mm in total length. A folded and fragmented bronze sheet, 4 mm wide. Two small tubular tuff beads, measuring 37 mm and 30 mm in length, and 10 mm in diameter. **Dating:** Archeologically dated to 500-700 CE. **Ancestry summary:** Excluded of the genome wide analysis due to contamination. **Additional information:** Presence of , an infant's talus bone. Stone slabs were disturbed, and the skeleton was incomplete.

##### 1.13 Guadalajara, Gualda

Contact person(s): Miguel Ángel Cuadrado Prieto

**Brief settlement history:** The settlement of El Tesoro-Carramantiel is a village and its necropolis, occupied in the late antique period, possibly during the 7th-8th centuries AD, based on the remains found in both areas.(Cuadrado-Prieto, 2002, 2022; Cuadrado-Prieto & Vallejo-Girvés, 1997).

**Excavation History:** The excavation began in 1992 after two tombs were illegally opened. Over five campaigns, 35 graves cut into sandstone were documented or excavated, some already looted in the past. Several houses from the nearby village, about 200 meters from the cemetery, were also documented.

**Geographic Information:** The remains are located in the Barranco Grande, a deep ravine that flows directly into the Tagus River in the La Alcarria region, in the south-central part of Guadalajara province.

**Summary of sampled materials:** The graves in this settlement mostly contain individual remains. However, some pits have been reused, with bones from the initial burial moved to the feet. Few grave goods were found: iron knife blades and personal adornments of silver and bronze. The anthropological study (Gómez-Pérez, 2006) identified the sex and age of some individuals and noted various pathologies and injuries. There is also a possibility that the bones were superficially affected by a fire at the site, as indicated by the dark coloration of some remains and other signs of indirect heat exposure on the bone surfaces.

**Graves:**

- **Genetic Identifier:** I19659. **Grave Identifier:** Tumba 4. **Grave Type:** Rectangular grave pit carbed in sandstone., NE-SW orientation. **Skeletal information:** Bones without anatomical connection due to clandestine removal; sex undetermined; around 20 years old; height 162.06 cm. **Grave goods:** incomplete iron knife blade with a triangular cross-section found mixed with the disturbed bones. **Dating:** Direct InCal 20 radiocarbon dated to670-774 calCE (1285±20 BP, PSUAMS-9703). **Ancestry summary:** Iberia_IA__+__WestAnatolia_Roman_Byzantine__+__North_African_Punic. **Additional information:** tomb raided during agost 1992.

- **Genetic Identifier:** I19660. **Grave Identifier:** Tumba 10. **Grave Type:** Rectangular pit carved in sandstone with a headrest carved at the top; slab partially covering the headrest; individual in primary supine position with arms extended along the body and head tilted to the left; orientation SE-NW. **Skeletal information:** possible male; 14-15 years old; height 164.62 ± 6.90 cm. **Grave goods:** no grave goods **Dating:** Archeologically dated to 500-800 CE**.** **Ancestry summary:** Iberia_IA__+__WestAnatolia_Roman_Byzantine__+__North_African_Punic. **Additional information:** Incomplete bilateral and symmetrical lumbarization of the first sacral vertebra; possible traumatic dislocation of the right shoulder joint; agenesis of the third

- **Genetic Identifier:** I19661. **Grave Identifier:** Tumba 11. **Grave Type:** Rectangular pit carved in sandstone with a headrest carved at the head end; slab partially covering the center, with smaller stones underneath; individual in primary supine position with arms extended along the body, head tilted on the chest; orientation SE-NW. **Skeletal information** Male, adult, 43.3-58.1 years; height 160.36 ± 6.90 cm. **Grave goods:** no grave goods. **Dating:** Archeologically dated to 500-800 CE. **Ancestry summary:** Iberia_IA__+__WestAnatolia_Roman_Byzantine__+__North_African_Punic. **Additional information:** Osteoarthritis.

- **Genetic Identifier:** I19662. **Grave Identifier:** Tumba 14. **Grave Type:** Rectangular pit carved in sandstone with a perimeter recess to fit a large covering slab; individual in primary supine position with arms crossed on the chest; orientation SE-NW. **Skeletal information:** Female, juvenile, 12 years ± 36 months. **Grave goods:** no grave goods. **Dating:** Archeologically dated to 500-800 CE. **Ancestry summary:** Iberia_IA__+__WestAnatolia_Roman_Byzantine__+__North_African_Punic. **Additional information:** Arthritic edges on the joint surfaces of the first two cervical vertebrae; possible fusion of two ribs.

- **Genetic Identifier:** I19663. **Grave Identifier:** Tumba 16. **Grave Type:** Rectangular pit carved in sandstone; individual in primary supine position with the only preserved arm on the abdomen; orientation SE-NW. **Skeletal information:** Female; 24-29 years; height 155.42 ± 5.92 cm. **Grave goods:** No grave goods. **Dating:** Direct InCal 20 radiocarbon dated to 772-974 calCE (1160±20 BP, PSUAMS-9707). **Ancestry summary:** Iberia_IA__+__WestAnatolia_Roman_Byzantine__+__North_African_Punic. **Additional information:** Osteoma on the skull; squatting marks; well-developed finger muscles, suggesting possible washerwoman.

- **Genetic Identifier:** I19664. **Grave Identifier:** Tumba 29. **Grave Type:** Rectangular pit carved in sandstone, part of a set of two on a promontory with a quadrangular perimeter recess to fit slabs over both; headrest carved at the head end and large broken slab covering the feet and another in the center; individual in primary supine position missing the skull with arms extended along the body; orientation NE-SW. **Skeletal information:** Male; 25.7-30.6 years; height 167.02 ± 6.90 cm. **Grave goods:** Half of a bronze liriform belt plate, decorated. **Dating:** Archeologically dated to 500-800 CE. **Ancestry summary:** Iberia_IA__+__WestAnatolia_Roman_Byzantine__+__North_African_Punic. **Additional information:** Head end looted with skull removed; possible trauma to the left shoulder during puberty.

#### 1.14 Guadalajara, Illana

Contact person(s): Miguel Ángel Cuadrado, José Martínez Peñarroya & Consuelo Vara Izquierdo.

**Brief settlement history:** The El Soto necropolis remained unknown until a trench for irrigation installation was dug in the namesake housing development, revealing a grave made of stone slabs. This development is on a promontory in Illana (Guadalajara), separate from the urban center. The remains of the Visigothic settlement of Recópolis are found around 20 km upstream, founded by Leovigildo in the mid-6th century CE.

This new addition to the map of early medieval settlements in the peninsula became known thanks to the property owner's decision to report the find to the cultural heritage authority in Guadalajara. Subsequent archaeological excavation of the grave revealed the existence of a necropolis, partly destroyed by a less responsible neighbor. The unique grave goods recovered provide valuable data for understanding the late period of the Visigothic monarchy in Hispania.

**Excavation History:** The archaeological work proceeded in two immediate phases. Initially, archaeological monitoring was conducted during the excavation of foundations for a new single-family home near the previously discovered burial. No evidence of prior habitation or burial was found, establishing the eastern limit of the necropolis. The foundation trench was ten meters square, intersected by another trench in its eastern half, both one meter deep, revealing a gravel layer beneath clayey sands.

The second phase involved excavating the funerary structure. After clearing spontaneous vegetation, the archaeological excavation began at parts of the structure exposed by the earlier irrigation trench. The trench had disturbed some of the skeletal remains, which were not fully filled in, according to the person who discovered it. The grave featured a mound of mortar debris, stones, and tile fragments. Inside, a denticulated lamina from a nearby deposit dating to the end of prehistory was found. The grave was made of gypsum stone slabs forming a cist, with a single-piece slab cover, broken during the trench excavation. This cover was sealed with mortar, remnants of which were found during the intervention.

**Geographic Information:** The current terrestrial communication system does not closely match the network that connected the population centers of the Visigothic Kingdom of Hispania. While recent times implemented a radial road system centered in Madrid, during the Visigothic period, the main axis connected cities like Zaragoza and Mérida in the northeast and southwest. Toletum, the capital, was centrally located on the plateau and along the Tagus River, which flows past the El Soto necropolis promontory.

This river course facilitated boat communications between the eastern and western landscapes of the southern plateau, nestled between clay-conglomerate landscapes and the extensive moors to the north and west, reaching Estremera in Madrid. To the east, the river flows between medium-sized elevations near Illana, intersecting with roads from Pastrana to the north and Tarancón to the south, in Cuenca province. This location served as a communication hub between terrestrial routes (Complutum-Segóbriga) and river routes (Recópolis-Toletum).

**Summary of sampled materials:** The archaeological record from this site was sparse but significant. Only one excavation unit, a grave with considerable grave goods at the time of burial and in optimal preservation, typically yields a substantial amount of material, including skeletal remains and artifacts. However, the circumstances of this find and the preservation state resulted in a limited but informative record. Key "index fossils" helped date the context and the partly destroyed necropolis on an adjacent plot.

Recovered bone fragments included eight skull pieces, about thirty limb fragments, fifty rib fragments, and fifteen vertebra fragments. In 25 cm of undisturbed sediment near the grave head, a gold tremissis was found. The obverse features a torso holding a cross with the legend "IND.IN.H.E.EGICA REX?" and upper interpunctions. The reverse shows a central cross with the legend "CORDOBAPATRICIA," separated by a cross. The well-preserved coin, with some edge wear and partial grafile, was minted in Córdoba between 687 and 700, during Égica's and Witiza's reigns. This dates the burial to the late 7th century CE.

**Graves:**

- **Genetic Identifier:** I19658. **Grave Identifier:** Tumba 1. **Grave Type:** Stone slab construction, SW-NE orientation. **Skeletal information:** unknown sex or age. **Grave goods:** Gold tremissis. **Dating:** Direct InCal 20 radiocarbon dated to 655-774 calCE (1325¬±20 BP, PSUAMS-9702). Archeological goods point towards $7^{th}$ century CE. **Ancestry summary:** Iberia_IA__+__WestAnatolia_Roman_Byzantine__+__North_African_Punic.

#### 1.15 Madrid, Boadilla

Contact person(s): Maite I. Garcia-Collado, Raúl Catalán

**Brief settlement history:** Boadilla, understood as the ensemble of the cemetery and residential areas, presents a wide occupational sequence. Before the early medieval phase, there is material evidence of Bronze (Calvo & Catalán, 2007; Dominguez-Fernández & López-Lancha, 2010), Iron (Dominguez-Fernández & López-Lancha, 2010) and Roman age occupations (Catalán et al., 2018). The foundation of the early medieval settlement is dated to the last quarter of the 5th century and it was inhabited probably until the middle of the 8th century. Domestic and productive areas were located north of Boadilla stream, while funerary areas were restricted to the south of the river. Residential areas were organised in family plots, each one composed of a main domestic building with sunken floors or based on stone banks, silos and other productive structures, such as water wells, cooking ovens or a kiln. Plots were divided by fences or wide empty areas interpreted as gardens, enclosures or spaces for other agrarian tasks.

The cemetery was made up of 181 burials, although it is likely that the original number was greater. It had a roughly rectangular outline. Burials were oriented predominantly in west-east direction and they never cut each other. Five different types of burials were recorded: simple pits, slabs burials, wall burials, burials made with reused building materials and burials in mixed materials. The total number of individuals identified was 226. The majority were found in primary deposits and in supine position, but there were also reductions, an ossuary and several undetermined secondary deposits. Around 20% of the individuals were buried with grave goods, including clothing items (fibulae, belt plates, buckles, buttons), jewellery (earrings, necklace beads, rings), tools (knives, tweezers, an arrowhead) and in one case a pottery container.

Finally, it is important to mention several items recovered in the residential area and the necropolis, as they reflect the access of some of the inhabitants to restricted goods, such as imported african pottery , amphorae or a rock crystal belt buckle, a very uncommon type usually related to high rank members such as the ones recovered in the frankish necropolis of Saint Diziers, France.

**Excavation History:** The first early medieval evidence in this area was a burial found in 1994 (Hernando & Iguácel, 1994) during the monitoring for the installation of a natural gas pipe for supplying the industrial park La Arboleda. During the first decade of the 21$^{st}$ century a new industrial park was promoted northeast of La Arboleda. This led to the discovery of the cemetery of Boadilla, covering about 3000 $m^2$, which was excavated between 2005 and 2008 by the company J. M. Rojas Arqueología under the direction of G. Garrido and J. Perera (Catalán & Rojas, 2009). Likewise, two new residential areas were promoted north of Boadilla stream. The district west of road A-42 is named Alameda del Señorío. Many of these plots revealed archaeological remains of different chronologies. Excavations were carried out in several of them, but only the results from plots R-23, R-24 and R-30 (approximately 2.5 ha), excavated between 2007 and 2008 (Calvo & Catalán, 2007), are public. East of road A-42 a vast area was surveyed for the development of a new residential zone in 2004. Immediately north of Boadilla stream the early medieval settlement of Cárcavas was discovered, including at least one burial (Dominguez-Fernández & López-Lancha, 2010).

**Geographic Information:** The archaeological site of Boadilla consists of a cemetery located north of the town of Illescas, in the province of Toledo, just 4 km south of the border with the region of Madrid. Its UTM ETRS89 coordinates are 30T 429705 4443432 and it is 585 meters above sea level. However, Boadilla belongs to a vaster settlement which extends up to 1 km around the cemetery. Boadilla is halfway between Toledo (35 km south) and Madrid (32 km north), 54 km southwest of the Roman city of Complutum (current Alcalá de Henares). The site is located in an area dominated by flatlands with a few small hills. The Boadilla stream flows in west-east direction and it divided the cemetery from the settlement. In the area there are many other seasonal water courses. This landscape would have been suitable for cereal fields. In the surroundings there are still some patches of holm oak meadows, which probably were greater in the past.

**Summary of sampled materials:** A minimum number of 172 individuals preserved skeletal remains for osteoarcheaological analysis (García-Collado et al., 2019). Macroscopic preservation was very poor and certainly biased any studies. The subadult to adult ratio of the buried population was 0.37, which is probably underestimating the proportion of subadults due to poor preservation, and the male to female ratio 0.25. Life expectancy at birth was calculated to be 35.2 years, although this data should be taken with caution due to preservation issues. In contrast, quality of the collagen recorvered was good. Carbon and nitrogen stable isotope ratios of 77 individuals revealed a very homogeneous diet predominantly based on C3 plants with small but consistent contribution of C4 crops, restricted but quite variable proportions of animal protein and no aquatic resources. Sampling for ancient DNA analysis was aimed at representing a variety of demographic categories, funerary rituals and diets. Teeth, preferentially molars, were sampled.

**Graves:**

- **Genetic Identifier:** I19888. **Grave Identifier:** BOA 017-1. **Grave Type:** Burial in mixed materials. Individual in primary supine position, W-E orientation. **Skeletal information:** Adult, >20 years old, undertermined sex. **Grave goods:** (No grave goods). **Dating:** Archeologically dated to 500-700 CE **Ancestry summary:** Excluded of the genome wide analysis due to unreliable mtDNA results, making it impossible to test mtDNA contamination. **Additional information:** $\delta^{13}C$ = -18.4‰, $\delta^{15}N$ = 10.2‰, %C = 34.0, %N = 12.9, C/N = 3.1, %coll = 3.4

- **Genetic Identifier:** I19889. **Grave Identifier:** BOA 018-1. **Grave Type:** Slabs burial. Individual in ossuary (secondary position). **Skeletal information:** Adult, >20 years, female. **Grave goods:** belt plate, necklace items. **Dating:** Archeologically dated to 500-700 CE **Ancestry summary:** CNE_EarlyMedieval__+__WestAnatolia_Roman_Byzantine. **Additional information:** ($\delta^{13}C$ = -18.0‰, $\delta^{15}N$ = 12.1‰, %C = 43.6, %N = 16.6, C/N = 3.1, %coll = 4.9

- **Genetic Identifier:** I19891. **Grave Identifier:** BOA 047-1. **Grave Type:** Burial made with reused building materials. Individual in primary supine position, W-E orientation. **Skeletal information:** Adult, >20 years, probably female. **Grave goods:** No grave goods. **Dating:** Archeologically dated to 500-700 CE **Ancestry summary:** Excluded of the genome wide analysis due to low coverage. **Additional information:** $\delta^{13}C$ = -18.3‰, $\delta^{15}N$ = 11.3‰, %C = 40.9, %N = 15.1, C/N = 3.2, %coll = 4.7

- **Genetic Identifier:** I19892. **Grave Identifier:** BOA 049-1. **Grave Type:** Simple pit burial. Individual in primary supine position, W-E orientation. **Skeletal information:** Adult, >20 years old, undertermined sex. **Grave goods:** No grave goods. **Dating:** Archeologically dated to 500-700 CE **Ancestry summary:** Excluded of the genome wide analysis due to low coverage.

- **Genetic Identifier:** I19894. **Grave Identifier:** BOA 053-1. **Grave Type:** Simple pit burial. Undetermined position. **Skeletal information:** Adult sp, >20 years old, undertermined sex. **Grave goods:** No grave goods. **Dating:** Archeologically dated to 500-700 CE **Ancestry summary:** Excluded of the genome wide analysis due to low coverage.

- **Genetic Identifier:** I19895. **Grave Identifier:** BOA 068-1. **Grave Type:** Simple pit burial. Individual in primary supine position, W-E orientation. **Skeletal information:** Adult, >20 years old, probably female. **Grave goods:** No grave goods. **Dating:** Archeologically dated to 500-700 CE **Ancestry summary:** Iberia_IA__+__CNE_Early_medieval. **Additional information:** $\delta^{13}C$ = -18.8‰, $\delta^{15}N$ = 10.2‰, %C = 15.0, %N = 5.7, C/N = 3.1, %coll = 0.6

- **Genetic Identifier:** I19896. **Grave Identifier:** BOA 086-1. **Grave Type:** Simple pit burial. Individual in primary supine position, W-E orientation. **Skeletal information:** Adult, >20 years, probably female. **Grave goods:** No grave goods. **Dating:** Archeologically dated to 500-700 CE **Ancestry summary:** CNE_EarlyMedieval__+__WestAnatolia_Roman_Byzantine__+__North_African_Punic. **Additional information:** $\delta^{13}C$ = -20.5‰, $\delta^{15}N$ = 8.7‰, %C = 43.7, %N = 15.8, C/N = 3.2, %coll = 5.8

- **Genetic Identifier:** I19897. **Grave Identifier:** BOA 094-1. **Grave Type:** ). Simple pit burial. Individual in primary supine position, W-E orientation. **Skeletal information:** Adult, >20 years , probably female. **Grave goods:** ring. **Dating:** Archeologically dated to 500-700 CE **Ancestry summary:** Iberia_IA__+__WestAnatolia_Roman_Byzantine__+__North_African_Punic.

- **Genetic Identifier:** I19898. **Grave Identifier:** BOA 105-4. **Grave Type:** Simple pit burial. Individual in reduction (secondary deposit). **Skeletal information:** Adult, >20 years old, probably female. **Grave goods:** No grave goods. **Dating:** Direct InCal 20 radiocarbon dated to 605-662 calCE (1400±20 BP, PSUAMS-10247) **Ancestry summary:** Iberia_IA__+__WestAnatolia_Roman_Byzantine__+__North_African_Punic. **Additional information:** $\delta^{13}C$ = -18.6‰, $\delta^{15}N$ = 10.2‰, %C = 43.5, %N = 14.8, C/N = 3.4, %coll = 5.6

- **Genetic Identifier:** I19899. **Grave Identifier:** BOA 112-1. **Grave Type:** Simple pit burial. Individual in primary supine position, W-E orientation. **Skeletal information:** Adult sp (>20 years), female. **Grave goods:** No grave goods. **Dating:** Archeologically dated to 500-700 CE **Ancestry summary:** Iberia_IA__+__North_African_Punic. **Additional information:** $\delta^{13}C$ = -18.7‰, $\delta^{15}N$ = 10.8‰, %C = 39.8, %N = 15.6, C/N = 3.0, %coll = 8.2

- **Genetic Identifier:** I19900. **Grave Identifier:** BOA 131-1. **Grave Type:** Simple pit burial. Individual in primary supine position, W-E orientation. **Skeletal information:** Adult, >20 years, female. **Grave goods:** earrings. **Dating:** Archeologically dated to 500-700 CE **Ancestry summary:** Iberia_IA__+__WestAnatolia_Roman_Byzantine__+__North_African_Punic.

- **Genetic Identifier:** I19903. **Grave Identifier:** BOA 160-1. **Grave Type:** Slabs burial. Individual in primary supine position, W-E orientation. **Skeletal information:** Adult, 21-36 years, male. **Grave goods:** No grave goods. **Dating:** Archeologically dated to 500-700 CE

**Ancestry summary:** Excluded of the genome wide analysis due to possible contamination. **Additional information:** $\delta^{13}C$ = -18.4‰, $\delta^{15}N$ = 8.9‰, %C = 39.3, %N = 13.8, C/N = 3.3, %coll = 7.6.

- **Genetic Identifier:** I19904. **Grave Identifier:** BOA 166A-1. **Grave Type:** Simple pit burial. Individual in primary supine position, W-E orientation. **Skeletal information:** Adult, >20 years, probably male. **Grave goods:** No grave goods. **Dating:** Archeologically dated to 500-700 CE **Ancestry summary:** Iberia_IA__+__CNE_Early_medieval. **Additional information:** $\delta^{13}C$ = -18.3‰, $\delta^{15}N$ = 10.3‰, %C = 39.7, %N = 15.0, C/N = 3.1, %coll = 7.9

- **Genetic Identifier:** I19906. **Grave Identifier:** BOA 182-1. **Grave Type:** Simple pit burial. Individual in primary supine position, W-E orientation. **Skeletal information:** Adult, >20 years, undertermined sex. **Grave goods:** necklace items. **Dating:** Archeologically dated to 500-700 CE **Ancestry summary:** Excluded of the genome wide analysis due to low coverage. **Additional information:** $\delta^{13}C$ = -17.2‰, $\delta^{15}N$ = 11.1‰, %C = 38.6, %N = 13.6, C/N = 3.3, %coll = 2.1

#### 1.16 Málaga, Acinipo

Contact person(s): Juan Manuel Jiménez Arenas, José Manuel Castaño.

**Brief settlement history:** The Acinipo archaeological site in Ronda, Málaga, has been known to local scholars since the 17th century. It was one of the first Roman cities in Baetica identified by humanist authors using its original name. The presence of a Roman theater in excellent condition attracted the attention of scholars and researchers early on. However, it wasn't until the latter part of the 20th century that the first archaeological studies began at this site, initially focusing on the iconic theater and the cultural period in which it was built (Del Amo, 1982). From this point, further excavations aimed to uncover more Roman-era landmarks in this ancient city, such as the forum and the baths (Puertas-Tricas, 1982; Puertas-Tricas & Aguayo-de-Hoyos, 1985).

During one of these excavations, older levels were documented, first identified from surface materials, which ultimately led to the creation of a research project focused on the pre-Roman phases of the settlement (Aguayo-de-Hoyos et al., 1987). Within this project, three excavation campaigns were carried out, mainly on the eastern spur of the Ronda la Vieja plateau, allowing the construction of an archaeological sequence from the late Copper Age to the early Roman period, including the late Bronze Age and the Iberian era, which were significant precursors to the Roman phase (Aguayo-de-Hoyos, 1997).

The available data suggest that this settlement was continuously inhabited from the Recent Prehistory (3200-2200 BCE) to the late Antiquity (5th-7th centuries CE), with the classical stages being when the city took shape, with its various internal and external elements, such as suburbs and necropolises. However, none of these research phases emphasized the areas beyond the topographic limits of the Mesa de Ronda la Vieja, where, based on indirect references, Roman-era necropolises were located—one to the north and another to the south (Castaño-Aguilar & Nieto-Gonzalez, 2008).

**Excavation history:** In 2004, a group of graves was discovered in the southern necropolis of Acinipo during agricultural work on land near the site. Most of these burials consisted of cremation urns and vessels, along with some inhumations, with a total of 48 graves documented in an excavated area of about 500 square meters. The preservation of the remains varied widely, affected not only by agricultural activities but also by the natural movement of the hillside, facilitated by the clay-rich geological substrate.

**Geographical information:** The southern necropolis of the Roman city of Acinipo is located on the southeastern slope of the Mesa de Ronda la Vieja, a plateau geologically formed from algal limestones and reaching an average elevation of 950 meters above sea level. This is the highest point among the depressions that make up the so-called Surco Intrabético. The necropolis is situated below this level, at about 900 meters above sea level, and its geology is mainly composed of sandstones and clays from the La Mina formation, with the limestones of the Las Mesas formation above them. This is where the archaeological site is located. Currently, it's about 20 km from the main town of the region, the city of Ronda, but in ancient times this distance was less significant because Acinipo was the primary urban center of the area.

**Summary of sampled material:** Among excavated graves, three types of burials were identified, likely indicating different rituals involving cremation and inhumation. Cremations were placed in painted ceramic urns of clear Iberian tradition, and in stone urns made from the limestone found in the area. The two inhumations documented (T-13 and T-37, from which the sample originated) were simple graves dug into the clay substrate, covered with flat tegulae tiles. The characteristics of these inhumed individuals suggest they were female, based on the grave goods found in the two burials, including bronze tweezers and a mirror. These objects were placed in wooden coffins or stretchers, of which only iron nails used in their construction remain. The inhumations were oriented east to west, with the deceased's head at the eastern end.

Apart from these burial types, there were other units identified by patches of ash, charcoal, bone fragments, and other materials (such as nails), with some covered by pitched tegulae tiles. These were interpreted as the remains of funeral pyres where some bodies were cremated (bustum or ustrinum). Based on the type of urns and some other material culture elements that provided chronological information, this southern sector of the Acinipo necropolis was dated to the 1st to the 2nd century CE. This dating was further supported by the distance of the graves from the southern gate and the lack of overlaps from later phases (Castaño-Aguilar et al., 2005).

**Graves:**

- **Genetic Identifier:** I23156. **Grave Identifier:** AC-04-N5; Tumba 13; 039. **Grave Type:** rave with flat tegulae cover and possible burial in a coffin or wooden platforms (presence of nails). **Skeletal information:** Possibly female, undetermined age. **Grave goods:** tongs and mirror. **Dating:** Archeologically dated to 1- 200 CE. **Ancestry summary:** Iberia_IA__+__WestAnatolia_Roman_Byzantine__+__North_African_Punic.

#### 1.17 Málaga, Estepona, Necrópolis Arroyo Vaquero

Contact person(s): Ildefonso Navarro Luengo, José Suárez Padilla, Juan Manuel Jiménez Arenas, Juan Manuel Jiménez Arenas

**Brief settlement history:** The Arroyo Vaquero site (Estepona, Málaga, Spain) has been known for a long time, although the first archaeological activity took place during the construction of a housing development. The excavation was carried out around two geographically and functionally defined areas (one consisting of building structures and the other of funerary structures). The building structures are made up of walls, some of which enclose a room paved with opus signinum. In another area, the remains of a poorly preserved mosaic were found, beneath which a burial was documented. The entire site has been considered a villa from the Late Roman period, although the information contained in the publication by Garrido Luque and Cisneros Franco (1987) is scarce.

Regarding the second area, a total of 33 tombs were documented, 24 of which were excavated. The construction techniques, burial ritual, and the few grave goods found suggested that it was a Hispano-Visigothic necropolis. Lastly, it is worth noting the documentation of knapped stone tools and handmade ceramics, pointing to the Late Bronze Age.

During 2001, due to the expansion of the highway N-340, a small intervention was carried out in the northern sector of the site, which uncovered a series of masonry structures and a pool filled in the second half of the 6th century or the first decades of the 7th century AD, according to the documented materials: several fragments of TSA D, in the form of Hayes 99, an oil lamp of the Atlante X C4 type (Bonifay type 56, n.22), and numerous slow-wheel-turned cooking pots typical of contexts from this period.

In 2006, a new sector of the Early Middle Age necropolis was documented, from which a total of 69 graves were excavated. Immediately west of the necropolis boundary, several large pits were excavated, filled with abundant fragments of coarse pottery, both wheel-turned and slow-wheel-turned, dating to the late 6th and early 7th centuries AD. The submission of a new construction project led to a new archaeological intervention, which consisted of cleaning and clearing the entire area excavated in previous interventions.

**Excavation History:** The necropolis of Arroyo Vaquero consists of 95 documented graves from the different excavations carried. It predominantly features simple graves, totaling 84, along with other more complex structures: one lined with bricks, eight lined with rubble, and two delineated by vertical slabs. The 2006 excavation permitted the knowledge of 69 new graves. Although there is no clear data on the organization of the occupied space, it is possible to interpret a certain grouping of burials in the northern sector, where five graves lined with rubble and two with vertical slabs stand out from the rest of the simple graves. These concentrations are common in late antique necropolises (Román Punzón, 2004). Although we are not certain whether this is due to spatial, temporal, or other factors, some researchers suggest a possible family organization.

Anthropological analysis leads to the suggestion that these are “family pantheons”, as no patterns were found concerning the sex or age of the buried individuals, with the same burial space shared by adults of both sexes and inmatures with adults.

The proposed chronological framework focuses on the 6th and 7th centuries AD, within the sphere of the Hispano-Visigothic world. This is mainly based on the typological affinity of the graves and the objects found in them with necropolises and settlements in the western Mediterranean, as well as with other bibliographical parallels. The review of all the information related to the necropolis of Arroyo Vaquero suggests linking their origin to a widely observed phenomenon during the 6th century, the appearance of cemeteries around a Christian place of worship that would have emerged as a re-adaptation of a sector of the pars urbana of a late antique villa, while their abandonment, equally linked to that of the settlement, could extend, at least by the type of graves used, until the 9th century, according to more recent excavations that date up to the 8th century.

**Geographic Information:** The Arroyo Vaquero Necropolis is located in the municipality of Estepona between Arroyo Vaquero and Arroyo Enmedio, borders the A-7 to the north, a private road to the south, the access road to the Arroyo Vaquero urbanization to the west, and other neighboring properties to the east (36°24'13.4"N - 5°11'37.6"W). Access is via the PK 155 of the A-7, from which the access road to the Arroyo Vaquero urbanization departs.

The geological substrate consists of Pliocene levels and Jurassic limestones. In general, it is a semi-compact sandy sediment of a beige-greenish color. This substrate is overlain by an alluvial level of reddish-yellow color with a sandy matrix containing numerous gravel and small stone inclusions. Immediately above the geological layer lie the remains of the Roman occupation.

**Summary of sampled materials:** Among the burial ritual objects, finds include two small ceramic jugs: one in G15, made of yellowish clay, with a single handle (handle not preserved), slightly convex base, and pear-shaped body, which could be classified as a variant of Izquierdo's Form 16 (1977); and another in H, made of whitish clay, also with a single handle (handle from the rim to the middle of the body), slightly convex base, and pear-shaped body. In both cases, the jugs were found at the pelvis level of the deceased. The most widely accepted hypothesis regarding their ritual use is that of a possible post-mortem baptism. Their presence has contributed to dating the cemetery to around the 6th and 7th centuries AD.

Regarding personal adornment objects, 14 simple bronze earrings were recovered in UEs E12 (2), I12 (2), G10 (2), I14 (1), G33 (3), I7 (2), and F16 (2); a double bronze bracelet in G10; three small bronze pins in F8; a bronze bracelet in I7; three bronze rings in N23, G8, and G10; an iron ring in I12; and a gold ring in G10.

**Graves:**

- **Genetic Identifier:** I23157. **Grave Identifier:** ARVI-06 No. 143; UE 9-3; Sondeo F; Tumba 19. **Grave Type:** Simple pit dug into the geological substrate and partially delineated by masonry. Remains of inhumations and an ossuary located at the feet. Minimum MNI: 4. Orientation W-E. **Skeletal information:** Unknown sex and age. **Grave goods:** Absent. **Dating:** Archeologically dated to 416-700 CE. **Ancestry summary:** Iberia_IA__+__WestAnatolia_Roman_Byzantine__+__North_African_Punic. **Additional information:** Intact tomb.

- **Genetic Identifier:** I23158. **Grave Identifier:** ARVI-06 No. 272; UE 17; Sondeo E; Tumba 12-2. **Grave Type:** Simple pit dug into the geological substrate. Remains of inhumations and an ossuary located at the feet. MNI: 4. Orientation W-E. **Skeletal information:** Male (?), Unknown age. **Grave goods:** Absent. **Dating:** Archeologically dated to 416-700 CE.

**Ancestry summary:** Excluded from the genome wide analysis due to low coverage. **Additional information:** Possibly excavated previously.

- **Genetic Identifier:** I23159. **Grave Identifier:** ARVI-06 No. 430; UE 6; Sondeo G; Tumba 29. **Grave Type:** Simple pit dug into the geological substrate, covered with stone slabs, filled with construction material and small stones. Remains of inhumations and an ossuary. MNI: 3. Orientation W-E. **Skeletal information:** Female, Adult. **Grave goods:** Absent. **Dating:** Archeologically dated to 416-700 CE. **Ancestry summary:** Excluded from the genome wide analysis due to contamination. **Additional information:** Intact tomb.

- **Genetic Identifier:** I23162. **Grave Identifier:** ARVI-06 No. 456; UE 15; Sondeo G; Tumba 38. **Grave Type:** Simple pit dug into the geological substrate, covered with stone slabs, although some are missing. Remains of inhumations and an ossuary at the feet. MNI: 3. Orientation W-E. **Skeletal information:** Male, Subadult. **Grave goods:** Absent. **Dating:** Direct InCal 20 radiocarbon dated to 600-652 calCE (1430±20 BP, PSUAMS-10251). **Ancestry summary:** Iberia_IA__+__WestAnatolia_Roman_Byzantine. **Additional information:** Possible looted tomb.

- **Genetic Identifier:** I23163. **Grave Identifier:** ARVI-06 No. 457; UE 15; Sondeo G; Tumba 35. **Grave Type:** Simple pit dug into the rock, covered with stone slabs, although some are missing. Inhumation and an ossuary at the feet. NMI: 3. Orientation W-E. **Skeletal information:** Male, Adult. **Grave goods:** Absent. **Dating:** Archeologically dated to 416-700 CE. **Ancestry summary:** Iberia_IA__+__WestAnatolia_Roman_Byzantine. **Additional information:** Bony callus on the right fibula. Possible looted tomb.

- **Genetic Identifier:** I23165. **Grave Identifier:** ARVI-06 No. 475; UE 18; Sondeo G; Tumba 41. **Grave Type:** Simple pit dug into the geological substrate, covered with stone slabs mixed with construction materials: bricks and decorated tiles. Remains of an inhumation. NMI: 1. Orientation W-E. **Skeletal information:** Male, young Adult. **Grave goods:** Absent. **Dating:** Archeologically dated to 416-700 CE. **Ancestry summary:** Iberia_IA__+__WestAnatolia_Roman_Byzantine__+__North_African_Punic. **Additional information:** Intact tomb.

- **Genetic Identifier:** I23166. **Grave Identifier:** ARVI-06 No. 482; UE 23; Sondeo G; Tumba 46. **Grave Type:** Simple pit dug into the geological substrate, covered with stone slabs mixed with construction materials: bricks and decorated tiles. Remains of multiple inhumations. MNI: 3. Orientation W-E. **Skeletal information:** Female, senile. **Grave goods:** Absent. **Dating:** Archeologically dated to 416-700 CE. **Ancestry summary:** Iberia_IA__+__WestAnatolia_Roman_Byzantine__+__North_African_Punic. **Additional information:** Intact tomb.

- **Genetic Identifier:** I23167. **Grave Identifier:** ARVI-06 No. 492; UE 6; Sondeo I; Tumba 74. **Grave Type:** Simple pit dug into the geological substrate, covered with large stone slabs Remains of an inhumation. MNI: 1. Orientación W-E. **Skeletal information:** Male, adult. **Grave goods:** Absent. **Dating:** Archeologically dated to 416-700 CE. **Ancestry summary:** Iberia_IA__+__WestAnatolia_Roman_Byzantine__+__North_African_Punic. **Additional information:** Intact tomb.

- **Genetic Identifier:** I23168. **Grave Identifier:** ARVI-06 No. 517; UE 9; Sondeo G; Tumba 32-3. **Grave Type:** Simple pit dug into the geological substrate, covered with large stone slabs and smaller stones. Remains of multiple inhumations. Orientation W-E. **Skeletal information:** Male(?), Subaldult. **Grave goods:** Absent. **Dating:** Archeologically dated to 416-

700 CE. **Ancestry summary:** Iberia_IA__+__WestAnatolia_Roman_Byzantine__+__North_African_Punic. **Additional information:** Intact tomb.

- **Genetic Identifier:** I23169. **Grave Identifier:** ARVI-06 No. 527; UE 6; Sondeo D; Tumba 2-1. **Grave Type:** Simple pit dug in the geological substrate. Remains of inhumation and an ossuary located at the feet. NMI: 2. Orientation W-E. **Skeletal information:** Male, young Adult. **Grave goods:** Absent. **Dating:** Archeologically dated to 416-700 CE. **Ancestry summary:** WestAnatolia_Roman_Byzantine__+__North_African_Punic. **Additional information:** Intact tomb.

- **Genetic Identifier:** I23170. **Grave Identifier:** ARVI-06 No. 532; UE 25; Sondeo G; Tumba ?. **Grave Type:** Simple pit dug into the geological substrate, without the cover preserved. Remains of inhumation. MNI: 1. Orientation W-E. **Skeletal information:** Unknown sex, senile. **Grave goods:** Absent. **Dating:** Archeologically dated to 416-700 CE. **Ancestry summary:** Excluded from the genome wide analysis due to low coverage. **Additional information:** Possible excavated previously.

- **Genetic Identifier:** I23171. **Grave Identifier:** ARVI-06 No. 542; UE 6; Sondeo G; Tumba 29-2. **Grave Type:** Simple pit dug into the geological substrate, covered with stone slabs mixed with construction material and small stones. Inhumations remains and an ossuary. NMI: 3. Orientation W-E. **Skeletal information:** Male, young Adult. **Grave goods:** Absent. **Dating:** Archeologically dated to 416-700 CE. **Ancestry summary:** WestAnatolia_Roman_Byzantine__+__North_African_Punic. **Additional information:** Intact tomb.

- **Genetic Identifier:** I23173. **Grave Identifier:** ARVI-06 No. 556; UE 23; Sondeo G; Tumba 26. **Grave Type:** Simple pit dug into the geological substrate, covered with stone slabs mixed with construction materials: bricks and decorated tiles. Remains of inhumations. NMI: 3. Orientation W-E. **Skeletal information:** Male (?), Unkown age. **Grave goods:** Absent. **Dating:** Archeologically dated to 400-600 CE. **Ancestry summary:** Iberia_IA__+__WestAnatolia_Roman_Byzantine__+__North_African_Punic. **Additional information:** Intact tomb.

- **Genetic Identifier:** I23174. **Grave Identifier:** ARVI-06 No. 557; UE 23; Sondeo G; Tumba 46-2. **Grave Type:** Simple pit dug into the geological substrate, covered with stone slabs mixed with construction materials: bricks and decorated tiles. Remains of inhumations. MNI: 3. Orientation W-E. **Skeletal information:** Male (?), Unknown age. **Grave goods:** Absent. **Dating:** Direct InCal 20 radiocarbon dated to 432-561 calCE (1565±20 BP, PSUAMS-10325). **Ancestry summary:** Iberia_IA__+__WestAnatolia_Roman_Byzantine__+__North_African_Punic. **Additional information:** Intact tomb.

- **Genetic Identifier:** I23175. **Grave Identifier:** ARVI-06 No. 572; UE 9; Sondeo G; Tumba 32. **Grave Type:** Simple pit dug into the rock, covered with large slabs and smaller stones. Remains of inhumations and an ossuary located at the feet. MNI: 3. Orientation W-E. **Skeletal information:** Male, unknown Age **Grave goods:** Absent. **Dating:** Archeologically dated to 416-700 CE. **Ancestry summary:** Iberia_IA__+__WestAnatolia_Roman_Byzantine__+__North_African_Punic. **Additional information:** Intact tomb.

- **Genetic Identifier:** I23179. **Grave Identifier:** ARVI-06 No. 604; UE 8; Sondeo G; Tumba 31. **Grave Type:** Simple pit dug into the geological substrate, covered with stone slabs mixed with construction materials. Remains of inhumations. MNI: 3. Orientation W-E. **Skeletal information:** Unknown sex and age. **Grave goods:** Bronze ring and a fragment of glassware in the ossuary. **Dating:** Archeologically dated to 416-700 CE. **Ancestry summary:** Iberia_IA__+__WestAnatolia_Roman_Byzantine__+__North_African_Punic. **Additional information:** Intact tomb.

#### 1.18 Malaga, Calle Rosarito

Contact person(s): Juan Manuel Jiménez Arenas & Sonia López Chamizo

**Brief settlement history**: The city of *Malaca* is situated at the western end of the Mediterranean Sea, south of the Iberian Peninsula, inheriting the Phoenician city of MLK. This continuity led Strabo (III, 4, 2) in the 1st century BC to assert that it had a Phoenician plan.

**Excavation History:** The excavation at Calle Rosarito in the municipality of Malaga (Spain) was performed by Arqueosur (Estudio de Arqueología S.L. enterprise during 2011.

**Geographic Information:** The necropolis is located on the western bank of the Guadalmedina River, serving as a liminal space between the realm of the living and the dead.

**Summary of sampled materials:** Tomb No. 6, discovered on Rosarito Street, is believed to belong to the late Roman phase of one of the necropolises in the city of Malaga. The Western Necropolis of Malaca is a space with a long funerary tradition that dates back to the 6th century BC and persists until the 5th century AD. Over almost 1000 years of sacred use, it incorporated different burial rites and afterlife beliefs, with its endurance not occurring by chance. The burial exhibits distinctive architectural features that allow us to explore both the internal structural aspects and monument-like signaling elements .

**Graves:**

- **Genetic Identifier:** I26172. **Grave Identifier:** Calle Rosarito. **Grave Type:** Evenly spaced nails found within the pit suggest wooden coffin burial. Tegulae added. E-W orientation, **Skeletal information:** Adult, >60 years old, Female. **Grave goods:** four oil lamps, a Levantine jug, four glass pieces, a handled ceramic plaque, a shell, and ritualistic metal nails. **Dating:** Direct InCal 20 radiocarbon dated to 126-232 calCE (1865±20 BP, PSUAMS-9811). **Ancestry summary:** Iberia_IA__+__WestAnatolia_Roman_Byzantine__+__North_African_Punic. **Additional information:** The woman was interred inside a wooden coffin, indicated by the evenly spaced nails found within the pit. Tegulae with a gabled roof were placed on the coffin, covered with soil, and later, a mound made of rubble masonry was added. The careful funeral ritual bestowed upon "Rosarito" by her community likely involved body preparation, perhaps through purification and dressing in a highly significant attire. While fabric remnants haven't been preserved, numerous glass elements have been documented, ranging from the neck to the feet, serving as decorative "strass" enriching the attire. Similarly, the treatment of the body included hairstyling, with a bun held by 3 *accus crinalis*. Within the coffin, various items were placed to accompany the deceased on her journey to the afterlife. These pieces were arranged in specific locations, suggesting adherence to particular ritual procedures. Altogether, four oil lamps, a Levantine jug, four glass pieces, a handled ceramic plaque, a shell, and ritualistic metal nails were arranged. Outside the coffin, two stones, faunal bone remains, a mortar, and two more oil lamps were positioned. The dating of the burial is supported by the chronologies of ceramic pieces, placing the ensemble between the late 2nd century and early 3rd century AD. Osteological study of the skeleton reveals that the individual was an elderly woman above 60 years old. Throughout her life, she experienced various traumatic events leaving traces on her bones. Notable are signs of malnutrition or poor nutrient absorption, congenital-idiopathic diseases, and potential infectious disease from zoonosis (López-Chamizo, 2023).

#### 1.19 Malaga, Pizarra, Castillejos de Quintana

Contact person(s): Francisca Rengel Castro, Enrique Viguera, Virgilio Martínez Enamorado & Juan Manuel Jiménez Arenas

**Brief settlement history:** This is a fortified hilltop settlement with two walled areas built in the 9th century. It was inhabited from the 7th to the 10th century. Known in Arabic sources as Santa María de Bobastro, this name is part of a political strategy naming various southern Bobastro fortifications after saints: Santa María, Santa Eulalia, and San Pedro. Initial settlement traces back to the Visigothic period (revealed by pottery found in situ in one of the graves: T5). The site saw renewed activity during the rebellion (fitna) of ʿUmar ibn Ḥafṣūn and his sons against the Umayyads (880-927), linking it directly to the counter-power base at Bobastro/Las Mesas de Villaverde (Ardales, Málaga).

**Excavation History:** Archaeological work began in 2020, focusing on cleaning prehistoric tombs in the Gibralmora mountains (a few kilometers from the early medieval necropolis) and some visible structures in the Visigothic-Andalusian hilltop village of Castillejos de Quintana, including the mentioned necropolis. Four rock-cut tombs oriented east were uncovered after repeated looting. Further excavation in 2022 revealed 13 more tombs of various sizes and other elements like walls, houses, and possibly a mosque.

**Geographic Information:** Los Castillejos de Quintana is located in the Gibralmora mountains within the municipality of Pizarra, Málaga province. The mountains rise over 400 meters above the Guadalhorce River. This site lies in the heart of Algarbía, the historical region of western Málaga known in Andalusian times as Algarbía. All architectural elements are carved into rock, made of Miocene calcarenite and conglomerates, creating a unique landscape similar to Bobastro, the rebellion center of Ibn Ḥafṣūn against the Umayyads.

**Summary of sampled materials:** The archaeological record includes significant ceramic finds from two phases (Visigothic and Andalusian). Both traditions are present at the site, but Visigothic pieces (7th-8th centuries) dominate the necropolis, indicating pre-Muslim occupation (before 711). Andalusian ceramics include various early types (until the 10th century), with no caliphal period references (post-929), suggesting abandonment after Córdoba's troops conquered the site in 921. Other materials found in the tombs include a few Visigothic buckles and other metal items.

**Graves:**

- **Genetic Identifier:** I29760. **Grave Identifier**: Grave I, Individual 1. **Grave Type**: Grave excavated in the rock layer. **Skeletal information**: Female. Age between 40 and 49 years. **Grave goods**: Funeral jar**. Dating**: Archeologically dated to 400-600 CE. **Ancestry summary**: Iberia_IA__+__WestAnatolia_Roman_Byzantine__+__North_African_Punic. **Additional information**: Caries and osteoarthritis. Raided tomb.

- **Genetic Identifier:** I29761. **Grave Identifier:** Tumba F, Individuo 1. **Grave Type:** Grave excavated in the rock layer. **Skeletal information:** Adult, between 40 and 52 years old, Male. **Grave goods:** Jar parts, amorphus ceramic pieces, roof tiles. **Dating:** Archeologically dated to 400-600 CE. **Ancestry summary:**

Iberia_IA__+__CNE_EarlyMedieval__+__WestAnatolia_Roman_Byzantine__+__North_African_Punic. **Additional information:** Enthesopathy. Raided tomb.

- **Genetic Identifier:** I29762. **Grave Identifier:** Tumba H, Individuo 1. **Grave Type:** Grave excavated in the rock layer. **Skeletal information:** Adult, between 35 and 51 years old, Female. **Grave goods:** Medieval metal ring. Ceramic and tile fragments **Dating:** Archeologically dated to 400-600 CE. **Ancestry summary:** Iberia_IA__+__WestAnatolia_Roman_Byzantine__+__North_African_Punic. **Additional information:** Caries. Raided tomb.

- **Genetic Identifier:** I29763. **Grave Identifier:** Tumba H, Individuo 2. **Grave Type**: Grave excavated in the rock layer. **Skeletal information**: Adult, between 20 and 35 years old, Female. **Grave goods**: Ceramic and tile fragments. **Dating:** Archeologically dated to 400-600 CE. **Ancestry summary**: Individual excluded from the genome wide analysis. **Additional information:** Raided tomb.

- **Genetic Identifier:** I29764. **Grave Identifier**: Grave I, Individual 2. **Grave Type:** Grave excavated in the rock layer. **Skeletal information**: Female. Age between 20 and 22 years old. Measurement between 1.53 and 1.60 meters. **Grave goods**: Ceramic fragments **Dating:** Archeologically dated to 400-600 CE. **Ancestry summary**: Iberia_IA__+__WestAnatolia_Roman_Byzantine__+__ North_African_Punic. Additional information: Orbital sieve. Raided tomb.

- **Genetic Identifier**: I29896. **Grave Identifier**: Grave G, M3. **Grave Type:** Grave excavated in the rock layer **Skeletal information:** Male. Age ranges from 20 to 41 years. Grave goods: Fragments of ceramics and tiles. **Dating:** Archeologically dated to 400-600 CE. **Ancestry summary:** Iberia_IA__+__WestAnatolia_Roman_Byzantine__+__North_African_Punic. **Additional information:** Raided tomb

#### 1.20 Mérida, Corralón de los Blanes Necropolis

Contact person(s): Macarena Bustamante Álvarez, Celia Chaves, Javier Heras Mora, Juan Manuel Jiménez Arenas

**Brief settlement history:** The city of Mérida is built on the ruins of Augusta Emerita, which was the westernmost capital of the Roman Empire. Shortly after its founding, around 25 BCE, Augustus placed it at the head of the newly created Province of Ulterior Lusitania. The city flourished over the centuries, becoming an important political, economic, and administrative center. It had two large forums, temples, baths, monumental aqueducts, and impressive entertainment buildings, like the Roman Theater, Amphitheater, and Circus. The governor Otho once ruled here, and later became the emperor of Rome; the Flavian dynasty's beautification trends and the policies of the Hispanic emperors also favored Emerita. Its continued prosperity was not interrupted by the military or economic crises of the 3rd century. On the contrary, the city was promoted administratively by the reforms of Emperor Diocletian. The governor of the Diocese of Hispania, which covered the entire Iberian Peninsula and parts of North Africa, resided here. Its role in this new administration brought significant social, economic, cultural, and religious development, giving Augusta Emerita a cosmopolitan character until the fall of the Roman Empire. Even then, Mérida was sought after by barbarian tribes who wanted its wealth and power, seeking to dominate Hispania (Heras 2018). The city eventually became the seat or residence of the court or oligarchy around the king of the "Suevi," a Germanic group that was part of the "Great Migrations" of the 5th century, which led to the military crisis that ended the Roman Empire (Heras, 2018; Heras et al., 2017; Heras & Olmedo-Gragera, 2015).

**Excavation History:** In 2005, excavations began at the site known as "Corralón de los Blanes," located at 41 Almendralejo Street in Mérida. This was a preventive excavation ahead of an urban development project that included building homes, commercial spaces, and underground parking on an area of half a hectare. The work stopped in 2007, with about 40% of the area still unexcavated. To this day, the project remains unfinished.

Despite this, the findings are exceptional (Heras et al., 2017). So far, the excavation has reached over 12 meters in depth and spans 900 years of history, from the earliest levels near the turn of the era, shortly after the founding of Augusta Emerita, to the 9th century, when part of the area was used to dig storage silos during the Islamic Emirate period.

During the Roman period, the first structures were monumental mausoleums, some preserved up to five meters in height. Successive archaeological phases also had a funerary nature, with cremations and inhumation burials continuing at least until the 3rd century. The use of the area as a necropolis likely ended in the 4th century, shifting to an extramural area that was eventually used as wasteland and dumps.

Each funerary or construction phase was preceded and followed by long periods when the land was used as a dump. Waste and soil acted as topographical agents, altering the original steep terrain, smoothing or reversing slopes over time.

This dynamic was especially significant during the second half of the 4th century, when urban waste accumulations were leveled to build a complex of industrial buildings, storage facilities, and possibly a "temple" for Eastern or mystery religions (Heras, 2011). This area was abandoned in the early 5th century and was violently and dramatically destroyed shortly after. People trapped in the rubble were never recovered for a proper burial. This event, dated through stratigraphy, material remains, and coins found on one of the bodies, is linked directly or indirectly to an assault on the city and the looting and destruction of areas outside the city walls.

On the ruins of these buildings and along the road leading from one of the city gates, new graves were dug for the most recent burials in this suburban area. Most of these were inhumation burials, similar in ritual and form to those of earlier centuries. However, some graves contained unique items like gold and silver jewelry or metal pieces that were likely part of the clothing worn by the deceased at their funeral. These bodies are of particular interest due to their origins, as they can be historically linked to the protagonists of the Great Migrations of the 5th century in Europe (Heras & Olmedo-Gragera, 2015).

**Geographic Information:** Mérida is currently the capital of Extremadura, the westernmost region of Spain. The region borders westernly with Portugal but the frontier is political as there is no geographical barriers. One of the main rivers on the Atlantic side of the Iberian Peninsula, the Guadiana River, flows through its urban core.

The Roman city was similar in size and covered a large area that spread across several promontories on the river's left bank. Beyond its walls, the city also stretched to the opposite river shore and into the surrounding plains. It reached the course of the Albarregas stream, a tributary that flows into the river at the city's base. The Roman-era burial grounds were located near the city walls, often close to the roads that led out of the city in all directions.

**Summary of sampled materials:** We sampled individuals from several archeological layers dating both to the Roman and Post-Roman period.

**Graves:**

- **Genetic Identifier:** I23272. **Grave Identifier:** BLA-ME-76; 8101. **Grave Type:** Inhumation in wooden box with perinatal between legs. W-E orientation. **Skeletal information:** adult female individual.. **Grave goods:** No grave goods found. **Dating:** Archeologically dated to 400-500 CE. **Ancestry summary:** Iberia_IA__+__CNE_Early_medieval__+__WestAnatolia_Roman_Byzantine. **Additional information:** Burial of a woman and another perinatal individual in fetal position between the legs. No material deposit and no remains of clothing. supine decubitus, with the arms extended on the hips and a perinatal -fetal- individual between the legs (8101_077), in poor state of preservation, although the skull can be seen with the face down. Lower jaw: second right premolar and first left premolar missing (closed jaw). It does not have wisdom teeth. Upper jaw with complete dentition, except for the wisdom tooth. It presents dental calculus.

- **Genetic Identifier:** I23273. **Grave Identifier:** BLA-ME-79; 8101. **Grave Type:** Burial in wooden box, unknown orientation. **Skeletal information:** Male, adult. **Grave goods:** Iron plaque at the head. **Dating:** Direct InCal 20 radiocarbon dated to 364-535 calCE (1645±20 BP, PSUAMS-8844). **Ancestry summary:** CNE_EarlyMedieval__+__WestAnatolia_Roman_Byzantine. **Additional information:**

Brother of Sample I22103 in Vilanova d'Alcolea site. Right arm over the chest. Left arm over the hip. Right leg over left leg. Retains the kneecaps. Nails in left clavicle, left arm and next to left tibia. Also next to right femur. Complete dentition. Wisdom teeth included. Very worn dental pieces.

- **Genetic Identifier:** I23276. **Grave Identifier:** BLA-ME-85; 8101. **Grave Type:** Burial in wooden box, W-E orientation. **Skeletal information:** Male, Adult. **Grave goods:** Iron plaque at the head**. Dating:** Archeologically dated to 400-500 CE. **Ancestry summary:** Iberia_IA__+__WestAnatolia_Roman_Byzantine__+__North_African_Punic__+__Kazakhstan_IA. **Additional information:** Right humerus on chest. Right hand under femur. Left hand on left femur. The patella is preserved. Lower jaw dislocated. Molar crowns very worn. Wisdom tooth present. Right tibia displaced to the left. Box nail next to the left humerus, left hand and right femur and iron plate at the head. Supine decubitus.

- **Genetic Identifier:** I23278. **Grave Identifier:** BLA-ME-95; 8101. **Grave Type:** Individual in an ossuary (MNI=3). **Skeletal information:** Unknown sex, adolescent. **Grave goods:** No grave goods. **Dating:** Archeologically dated to 200-300 CE. **Ancestry summary:** Iberia_IA__+__WestAnatolia_Roman_Byzantine. **Additional information:** Ossuary, without apparent grave, in which two human skulls and long bones (such as 6 femurs, 2 fibulas and 5 tibias, 2 ulnae and 2 radii) are identified, as well as a coxal bone, 3 hips, ribs. One of the skulls belongs to an adolescent individual, which preserves only the upper jaw - the wisdom tooth is missing - and the frontal bone. Secondary deposition

- **Genetic Identifier:** I23280. **Grave Identifier:** BLA-ME-103; 8101. **Grave Type:** Burial in wooden box with gabled tegulae cover. W-E orientation. **Skeletal information:** Male, Adult. **Grave goods:** No grave goods. **Dating:** Archeologically dated to 400-500 CE. **Ancestry summary:** CNE_EarlyMedieval__+__WestAnatolia_Roman_Byzantine. **Additional information:** Bone pieces displaced from their anatomical position. Skull turned to the left. Displaced lower jaw. Displaced ribs, arms and hands. Shoulder blades above the hip. No trousseau or dress.

- **Genetic Identifier:** I23283. **Grave Identifier:** BLA-ME-161; 8101. **Grave Type:** Burial in wooden box. **Skeletal information:** Undetermined sex, Adolsescent, between 13 and 16 years old. **Grave goods:** Aristocratic ornaments (gold earrings with prismatic top). **Dating:** Archeologically dated to 400-500 CE. **Ancestry summary:** Excluded of genome wide analysis due to low coverage. **Additional information:** Displaced bone remains. Lower jaw displaced with respect to the rest of the skull. Rib bones, vertebrae, hip bones piled up. Possible metabolic pathology in femurs (criba femoralis). Grave goods indicating aristocracy ("Suebian Princess").

- **Genetic Identifier:** I23284. **Grave Identifier:** BLA-ME-163; 8101. **Grave Type:** Inhumation in simple grave NE-SW Orientation. **Skeletal information:** Female, Adult between 30 and 45 years old. **Grave goods:** Aristocratic dress/dress (gold needles and beads). **Dating:** Archeologically dated to 200-550 CE. **Ancestry summary:** CNE_EarlyMedieval__+__WestAnatolia_Roman_Byzantine__+__North_African_Punic. **Additional information:** Arms arranged along the body. Right hand on right hip and left hand at the center of the hip. Skull turned to the right. Sternum is preserved. No patella. Ribs in bad condition, as well as femurs. Age between 38-39 years. Simple fossa with the particularity of a recess or "flange" in his feet. Supine decubitus.

- **Genetic Identifier:** I23288. **Grave Identifier:** BLA-ME-312; 8101. **Grave Type:** Inhumation in simple grave. N-S orientation. **Skeletal information:** Undetermined sex and age. **Grave goods:** No grave goods. **Dating:** Archeologically dated to 400-500 CE. **Ancestry summary:** Iberia_IA__+__WestAnatolia_Roman_Byzantine__+__North_African_Punic. **Additional information:** Fragmented skull turned to the right. Fragmented mandible, both lower and upper jaw. Arms along the trunk, with left hand on the hip. Legs extended. Epiphyses not attached to the diaphysis.

- **Genetic Identifier:** I23289. **Grave Identifier:** BLA-ME-1091; 8102. **Grave Type:** Partially obliterated inhumation. Unknown oreintation. **Skeletal information:** Undetermined sex and age. **Grave goods:** No grave goods. **Dating:** Archeologically dated to 400-500 CE. **Ancestry summary:** Excluded of genome wide analysis due to low coverage. **Additional information:** Burial next to the NE corner of building A4, affected by the mechanical cuts. Only the trunk and extremities are preserved, not the skull and left arm. It presents welded vertebrae. The epiphyses of the femur (head and knee) are still unwelded. No trousseau or dress.

- **Genetic Identifier:** I23752. **Grave Identifier:** BLA-ME-190; 8101. **Grave Type:** Inhumation in simple grave with tegulae cover. W-E Orientation. **Skeletal information:** Undetermined sex and age. **Grave goods:** No grave goods. **Dating:** Archeologically dated to 400-500 CE. **Ancestry summary:** Iberia_IA__+__WestAnatolia_Roman_Byzantine. **Additional information:** Possible skull fracture. Head turned to the left. Arms at the sides of the trunk. Left forearm higher than the right. No wisdom teeth. Very worn dental crowns. One of the molars has caries on the side. The sutures of the skull, especially a parietal, seem to have suffered a fracture. Supine decubitus. No trousseau or dress.

- **Genetic Identifier:** I23753. **Grave Identifier:** BLA-ME-211; 8101. **Grave Type:** Burial in a simple grave. W-E Orientation. **Skeletal information:** Undetermined sex and age. **Grave goods:** No grave goods. **Dating:** Direct InCal 20 radiocarbon dated to 364-535 calCE (1645±20 BP, PSUAMS-8844). **Ancestry summary:** CNE_EarlyMedieval__+__WestAnatolia_Roman_Byzantine. **Additional information:** Legs parallel. Skull turned to the right. Right ulna over the left humerus. Radius and ulna arched under the hip, as well as the hand. Right hand on hip. Left foot up and right foot turned to the left. No wisdom teeth and very worn teeth. Indeterminate adult (field). Supine decubitus.

- **Genetic Identifier:** I23754. **Grave Identifier:** BLA-ME-226; 8101. **Grave Type:** Roof-roofed interment with two-column signage. W-E Orientation **Skeletal information:** Undetermined sex and age. **Grave goods:** No grave goods. **Dating:** Direct InCal 20 radiocarbon dated to 408-537 calCE (1625±20 BP, PSUAMS-10253). **Ancestry summary:** CNE_EarlyMedieval__+__North_African_Punic. **Additional information:** Arms along the trunk. Right leg flexed to the left. Sternum and patellae preserved. Right hand under right leg. Fragmented skull. Lower dentition missing two right molars (closed socket). Left wisdom tooth. First molar extracted. Very worn dentition. Indeterminate adult (field). Supine decubitus. No trousseau or dress.

- **Genetic Identifier:** I23755. **Grave Identifier:** BLA-ME-233; 8101. **Grave Type:** Burial in wooden box, delimited with ceramic and marble fragments.NE-SW Orientation. **Skeletal information:** Undetermined sex and age. **Grave goods:** No grave goods. **Dating:** Direct InCal 20 radiocarbon dated to 259-415 calCE (1695±20 BP, PSUAMS-10254). **Ancestry summary:** Iberia_IA__+__WestAnatolia_Roman_Byzantine. **Additional information:** Arms along the body, with hands on the femur. Skull very deteriorated, turned to the left. Supine decubitus. No trousseau or dress.

- **Genetic Identifier:** I23757. **Grave Identifier:** BLA-ME-249; 8101. **Grave Type:** Burial in a single grave with a gable roof. NE-SW Orientation. **Skeletal information:** Male, Adult. **Grave goods:** No grave goods. **Dating:** Archeologically dated to 200-300 CE. **Ancestry summary:** Excluded of genome wide analysis due to low coverage. **Additional information:** Very worn teeth. Supine decubitus

- **Genetic Identifier:** I23758. **Grave Identifier:** BLA-ME-279; 8101. **Grave Type:** Burial in double simple grave. W-E orientation. **Skeletal information:** Undetermined sex and age. **Grave goods:** No grave goods. **Dating:** Archeologically dated to 400-500 CE. **Ancestry summary:** Iberia_IA__+__WestAnatolia_Roman_Byzantine__+__Kazakhstan_Sarmatian_IA. **Additional information:** Supine decubitus. No trousseau or dress.

- **Genetic Identifier:** I23761. **Grave Identifier:** BLA-ME-287; 8101. **Grave Type:** Burial in a single grave with a gable roof.W-E orientation. **Skeletal information:** Undetermined sex, adult. **Grave goods:** No grave goods. **Dating:** Archeologically dated to 400-500 CE. **Ancestry summary:** CNE_EarlyMedieval__+__WestAnatolia_Roman_Byzantine__+__North_African_Punic. **Additional information:** Arms parallel to the trunk with hands on hips. Left leg flexed. Crowns very worn. Wisdom tooth. Adult. Orientation: W-E. Supine decubitus. No trousseau or dress.

- **Genetic Identifier:** I23762. **Grave Identifier:** BLA-ME-301; 8101. **Grave Type:** Burial in wooden box. Orientation: NE-SW.: **Skeletal information:** Male, Adult. **Grave goods:** No grave goods. **Dating:** Archeologically dated to 200-300 CE. **Ancestry summary:** Iberia_IA__+__WestAnatolia_Roman_Byzantine__+__North_African_Punic. **Additional information:** Head turned to the left. Curved spine. Preserved up to the start of the left femur. Hands on the hip. Some nails have appeared, some with remains of wood. The right hip is not preserved. Right radius and ulna not complete. Degenerative process of the head of the left humerus. Part of the sternum is preserved. All the molars of the lower jaw are missing (closed sockets). No trousseau or dress.

- **Genetic Identifier:** I23763. **Grave Identifier:** BLA-ME-305; 8101. **Grave Type:** Inhumation in simple grave. SE-NW oreintation. **Skeletal information:** Male, adult. **Grave goods:** No grave goods. **Dating:** Archeologically dated to 400-500 CE. **Ancestry summary:**

Iberia_IA__+__North_Africa_Punic. **Additional information:** The head rests on fragments of brick. Very worn dentition. No trousseau or dress.

- **Genetic Identifier:** I23764. **Grave Identifier:** BLA-ME-308; 8101. **Grave Type:** Inhumation possibly in wooden box (*caligae* in lateral leg). NW-SE Orientation. **Skeletal information:** Male, Adult, between 45 and 50 years old **Grave goods:** *caligae* or military sandals on side of leg. **Dating:** Archeologically dated to 400-500 CE. **Ancestry summary:** Iberia_IA__+__WestAnatolia_Roman_Byzantine__+__North_African_Punic. **Additional information:** Head turned backwards. Arms parallel to the trunk and hands on femurs. Feet together turned to the right. Two nails, two iron objects and an "iron plate with spikes". Iron object parallel to the left tibia (*caligae*). Fragmented brick under the spine. Very worn and decayed dentition. Traumatic pathology in left hand. Possible military (*caligae* or military sandals on side of leg).

- **Genetic Identifier:** I23765. **Grave Identifier:** BLA-ME-310; 8101. **Grave Type:** Simple semi-deep burial. **Skeletal information:** Undetermined sex and age **Grave goods:** No grave goods. **Dating:** Archeologically dated to 200-300 CE. **Ancestry summary:** Iberia_IA. **Additional information:** No trousseau or dress.

- **Genetic Identifier:** I23766. **Grave Identifier:** BLA-ME-331; 8101. **Grave Type:** Burial in wooden box inside tegulae cist with flat cover. W-E orientation. **Skeletal information:** Undetermined sex and age **Grave goods:** No grave goods. **Dating:** Direct InCal 20 radiocarbon dated to **Ancestry summary:** CNE_EarlyMedieval__+__WestAnatolia_Roman_Byzantine. **Additional information:** Skull and trunk displaced from their normal anatomical position and "stacked". The legs are in their original position, parallel. The diaphyses are not joined. No trousseau or dress.

- **Genetic Identifier:** I23768. **Grave Identifier:** BLA-ME-347; 8101. **Grave Type:** Single grave burial, partially preserved. **Skeletal information:** Undetermined sex and age **Grave goods:** No grave goods. **Dating:** Archeologically dated to 200-300 CE. **Ancestry summary:** Iberia_IA__+__North_African_Punic. **Additional information:** Partially missing because cut by a later burial. No trousseau or dress.

- **Genetic Identifier:** I23770. **Grave Identifier:** BLA-ME-1147; 8102. **Grave Type:** Burial in a simple grave. NE-SW orientation.. **Skeletal information:** Undetermined sex and age **Grave goods:** No grave goods. **Dating:** Archeologically dated to 400-500 CE.**Ancestry summary:** Iberia_IA__+__WestAnatolia_Roman_Byzantine__+__North_African_Punic. **Additional information:** Inhumation in south corner of building A4. Right arm above the hip. In general, very deteriorated remains. In a pit excavated in the collapse of the building. No trousseau or dress.

- **Genetic Identifier:** I23771. **Grave Identifier:** BLA-ME-1189; 8102. **Grave Type:** Burial in a wooden box in a simple grave. W-E orientation. **Skeletal information:** Female, Adult,between 30 and 40 years old.. **Grave goods:** Aristocratic ornament/dress (necklace and gold needles. **Dating:** Archeologically dated to 400-500 CE. **Ancestry summary:** Iberia_IA__+__CNE_Early_medieval__+__WestAnatolia_Roman_Byzantine. **Additional information:** Skeleton with a large part of the pieces displaced from their original anatomical position, with only the legs in their normal location. Skull displaced and turned to the right (south), as well as the jaw - upside down. Arms arranged approximately next to the trunk and under the pelvis. Nails at the head, waist and feet. Fairly complete dentition, although with significant signs of wear. Rounded and open hip (female). Rounded ridge under the neck/skull. Adult due to cranial sutures and dental wear. Burial under road. Skeletal height: 1.51 m.. Supine decubitus. Allophyseal. Grave goods indicating aristocracy ("Suebian Princess").

- **Genetic Identifier:** I23772. **Grave Identifier:** BLA-ME-1222; 8102. **Grave Type:** Inhumation in box in grave with tegulae cover.NW-SE orientation. **Skeletal information:** Undetermined sex, adult, between 35 and 45 years old. **Grave goods:** No grave goods. **Dating:** Archeologically dated to 400-500 CE. **Ancestry summary:** CNE_Early_medieval. **Additional information:** Trench burial with three tegulae protecting the body. The skull faces SW, mandible displaced. Arms parallel to the trunk and hands under the hips. Legs straight and parallel. Supine decubitus. Allophysis. Inhumation without grave goods or clothing, next to I23771.AG. BLA-ME-1189; 8102.

- **Genetic Identifier:** I23773. **Grave Identifier:** BLA-ME-1348; 8102. **Grave Type:** Inhumation in simple grave (incomplete**).** Unknown orientation. **Skeletal information: Undetermined sex and age. Grave goods:** No grave goods. **Dating:** Direct InCal 20 radiocarbon dated to 215-326 calCE (1800±20 BP, PSUAMS-10256). **Ancestry summary:** Iberia_IA__+__North_African_Punic.

- **Genetic Identifier:** I24953. **Grave Identifier:** BLA-ME-1151; 8102. **Grave Type:** Ossuary or secondary deposit in ceramic container. **Skeletal information:** Undetermined sex, adult, advanced age. **Grave goods:** No grave goods. **Dating:** Archeologically dated to 400-500 CE.**Ancestry summary:** Iberia_IA__+__WestAnatolia_Roman_Byzantine__+__North_African_Punic. **Additional information:** Secondary inhumation of which a ceramic vessel (amphora) is part, where the smaller bones are collected. There is wear in teeth and vertebrae and robust bones (advanced age?). The skull is not preserved; the hip is very fragmented and, due to its size, it is not included in the container. No trousseau or dress.

- **Genetic Identifier:** I24954. **Grave Identifier:** BLA-ME-1186; 8102. **Grave Type:** Inhumation in stone cist. E-W orientation. **Skeletal information:** Age range 18-45 (probably between 22-29 years),.**Grave goods:** No grave goods. **Dating:** Archeologically dated to 400-500 CE. **Ancestry summary:** Excluded of genome wide analysis due to possible contamination. **Additional information:** Burial on collapsed building in north corner, delimited with stone blocks supported on a floor of amortized signinum. Incomplete skeleton. The upper extremities are missing (affected and altered by roots) and the lower extremities of both legs (cut by contemporary constructions). Slight green stain on ribs (remains of bronze object). The skull appears disarticulated with respect to the vertebrae, of which the cervical ones are missing. The mandible is also missing. In the preserved ones the wear of the crowns is appreciated. There are fractures in dry bone, in right radius, in epiphysis, left ulna in distal epiphysis; right femur in external condyle and tibia. Skeletal height: 1.30 m.

Orientation: E-W. Adult female. Age range 18-45 (22-29 years), from the degree of closure of the cranial sutures and the characteristics of the symphysis pubis. No trousseau or dress.

- **Genetic Identifier:** I28351. **Grave Identifier:** 8101-UE 285. **Grave Type: Burial in a simple grave. W-E** Orientation. **Skeletal information:** Undetermined sex, indeterminate subadult. **Grave goods:** No grave goods. **Dating:** Archeologically dated to 400-500 CE. **Ancestry summary:** Excluded of genome wide analysis due to low coverage. **Additional information:** Skull bones fragmented and "piled up". Missing hip and left femur. Wisdom tooth in socket. Interior calculus. Very worn. No trousseau or dress.

- **Genetic Identifier:** I28455. **Grave Identifier:** 8101-UE 160. **Grave Type:** Inhumation in simple grav, N-S orientation. **Skeletal information:** Female, infant, between 8 and 10 years old. **Grave goods:** Aristocratic dress (fibulae, plates, pendant) and burial aristocratic ornamental deposit. **Dating:** Archeologically dated to 400-500 CE. **Ancestry summary:** Excluded of genome wide analysis due to low coverage. **Additional information:** Subadult burial, arms along the body. Hands under the pelvis. Legs arched. Good state of preservation. Unattached patella. Part of the sternum is preserved. Skull very deteriorated. Supine decubitus.

#### 1.21 Segovia, Necrópolis de Castiltierra

Contact person(s): Andrés Carretero, Teresa Espinosa, Sergio Vidal Álvarez, Paula Pagés Alonso & Beatriz Campderá Gutiérrez

**Brief settlement history:** The Late Antiquity necropolis of Castiltierra is one of the largest and better known to date within ths cronology in Iberia. Its chronological extension goes from the late 5th century until the beginning of the 8th. It is located between two cities that were important during Roman times, Confluencia and Tiermes. During the 4th century crisis and later centuries, the population remained in small rural centers and bury their dead in necropolises such as Castiltierra or the nearby Duratón (Balmaseda & Arias, 2017).

**Excavation History:** The necropolis was discovered by chance during the construction of a road in the 1920s. This led to a time of pillaging and destruction of the graves. During the 1930s the excavation campaigns began, codirected by Emilio Camps and Joaquín María Navascués (1932, 1934 and 1935). Later, after the Spanish Civil War, there were two other campaigns, a short one in 1940 and a longer one directed by Julio Martínez Santa-Olalla in 1941.

**Geographic Information:** This necropolis is located within the municipality of Fresno de Cantespino, 1km away from the modern village of Castiltiera, in the northeast of the province of Segovia. The topography is quite flat but it is at a high altitude due to its proximity to the Sierra de Ayllón. The necropolis' area is quite large (according to J. Werner, the necropolis would cover an area of at least of 800m x 300m. The extent of the 1941 campaing by Santa-Olalla, would cover about 1700$m^2$) and it is on clay soil (Balmaseda & Arias, 2017)

**Summary of sampled materials:** The sampled materials came from the Santa-Olalla excavation from 1941. Unfortunately skeleton sampling, handling and storage practices didn't follow a scientific procedure. The individuals reported in this study were lower jaws from this site which were stored in a box without much archological context.

**Graves:**

- **Genetic Identifier:** I18408. **Grave Identifier:** 1973/58/CAS/C42/1688a: 30 L-F (4le). **Grave Type:** untracable. **Skeletal information:** untracable. **Grave goods:** untracable. **Dating:** Archeologically dated to 500-600 CE. **Ancestry summary:** Iberia_IA__+__WestAnatolia_Roman_Byzantine__+__North_African_Punic.

- **Genetic Identifier:** I18411. **Grave Identifier:** 1973/58/CAS/C42/1695: 30 L-F (4le). **Grave Type:** untracable. **Skeletal information:** untracable. **Grave goods:** untracable. **Dating** Archeologically dated to 500-600 CE. **Ancestry summary:** Iberia_IA__+__WestAnatolia_Roman_Byzantine__+__North_African_Punic

#### 1.22 Tarragona, Amposta, Mas Cabiscol-Finca Jornet

Contact person(s): Josep Maria Vergès (sampling and dating)/ Samuel Sardà Seuma (Project manager)

**Brief settlement history:** The Mas Cabiscol-Finca Jornet site has traditionally been interpreted as a particularly interesting burial area. This is because, based on documented characteristics such as the coexistence of inhumations and cremations, tumulus graves, ceramic types, etc., it seemed to belong to the late Bronze Age and the early Iron Age (roughly between 800 and 600 BCE) (Esteve, 2000; Garcia-Rubert, 2005; Noguera-Guillén, 2006). As a result, it was understood and explained as a reference point for funeral rituals that illustrate the transition from inhumation to cremation in the region near the ancient mouth of the Ebro River. However, the interest in the site and the possible issues or inaccuracies arising from the 1968 excavation by a group of local enthusiasts prompted a reevaluation of the ceramic materials. This led to its inclusion in the project: Settlement, Funeral Rituals, and Social Evolution at the Mouth of the Ebro (1st Millennium BCE) (CLT009/18/00088), granted by the Department of Culture (Generalitat de Catalunya). The project is led by Samuel Sardà (GRESEPIA-URV).

The re-examination of the ceramic materials has provided important insights into the final stage of the necropolis, which was traditionally defined by three small fragments of Phoenician wheel-made pottery, suggesting a date around 600 BCE. However, the re-study of these materials has pushed this date to 500 BCE or later, based on the presence of ancient B-type pottery (Py 1990: 534), likely an amphora A-MGR1. The rim, specifically, corresponds to the A-MGR bd1 or bd2 shape, more likely the first, suggesting a chronology between 600-500 BCE (Sourisseau, 1993).

As part of this re-evaluation and through the collaboration of Josep Maria Vergés (IPHES-URV), the site was included in an the present study. In October 2020, bone samples (mostly dental pieces) were selected for radiocarbon dating to refine the chronology of the documented inhumations and to determine the specific period of this burial area. The analyses by Beta Analytic Laboratory (Miami, USA) and Penn State Radiocarbon Laboratory yielded surprising results, indicating these were roman and late-antiquity inhumations, with dates 129-244 cal CE and 574-657 cal CE (95.4% confidence interval). With these radiocarbon dating results, it is now known that the Mas Cabiscol-Finca Jornet inhumations (Amposta) correspond to burials carried out during the Roman and Late Antiquity periods. This suggests that the area was originally used as a burial space during the Early Iron Age, as evidenced by the funerary urns and other ceramic vessels housed at the Museum of the Terres de l'Ebre (Amposta), and was later reused as a burial site during more recent periods.

**Excavation History:** The site was identified and excavated in March 1968 by the archaeologist Francesc Esteve Gálvez and his team of local collaborators. They documented a tumulus necropolis with inhumation and cremation graves. The site was identified during agricultural work when a tractor's plow disturbed stones that seemed to be from ancient constructions, along with a skull and other human bones. A total of 11 graves were excavated, one of which was defined as a funerary structure with stones forming circles (grave number 7). Among the documented structures, grave number 4 stands out because a significant layer of burnt soil was found, suggesting that a primary cremation might have taken place at the burial site (Esteve, 2000).

**Geographic Information:** The Mas Cabiscol-Finca Jornet site (Amposta, Montsià, Tarragona) is located at an altitude of 101 meters and covers an area of nearly 500 square meters (Noguera-Guillén, 2006). It's on the right bank of the Ebro River, about 3 km from the river's course, just west of the base of the Montsianell hill. From the site, there's a clear view of the natural corridor between the Sierra del Montsià and the Sierra de Freginals, which leads to the Ulldecona valley. This site could be the necropolis where the deceased from the neighboring Early Iron Age settlement of Lo Tapà (Amposta, Montsià), located about 500 meters southwest, were buried (Sardà-Seuma, 2021).

**Summary of sampled materials:** The bone remains recovered from the site are all stored at the Museu del Montsià (Amposta) with registration numbers 1141 and 1144. They belong to at least three individuals. Based on data from the excavators, they all seem to come from what is known as Grave 7 (Esteve, 2000). Specifically, the selected samples are all dental pieces, two with registration number 1141 and one with number 1144.

The documented and described funerary structures from the 1968 excavation correspond to graves typical of the Late Bronze Age and the Early Iron Age in the Ebro Valley. Specifically, Grave 7 has the typical format of flat circular mounds, with stones arranged horizontally and a single vertical stone in the center to mark the location of the urn. The funerary urns were handmade and, in some cases, had grooved decoration. They were placed in a simple pit or loculus without a cist. Some ceramic pieces found outside the graves were interpreted as offering vessels(Noguera-Guillén, 2006). The general characteristics of these funerary structures are similar to those found in Early Iron Age necropolises such as Sebes (Flix), Els Castellons (Flix) (Belarte-Franco et al., 2013, 2018; Noguera-Guillén, 2006), Coll del Moro (Gandesa) (Rafel & Hernández, 1992), and La Pedrera (Vallfogona de Balaguer-Térmens) (Vázquez et al., 2012).

The descriptions of the excavation of Grave 7 (from which the dental pieces for the current genomic study originate) are particularly important. It was a mound grave with evidence of inhumations along with fragments of various handmade ceramic vessels, typical of the Early Iron Age (Esteve, 2000). These findings supported the idea of inhumations in a mound structure possibly erected during the Late Bronze Age, similar to known cases like Tumulus 27, a collective inhumation grave in the Castellets II necropolis (Mequinenza) (Royo, 1996). However, the radiocarbon date obtained in 2020 suggest that this might be a reused mound structure serving as a tomb or funerary deposit in the Roman Period.

**Graves:**

- **Genetic Identifier:** I27078. **Grave Identifier:** Finca Jornet (Amposta); FJ-M2 (FJ-1144). **Grave Type:** Slabs burial in a reused tumulus grave. **Skeletal information:** Adult, >20 years old, undertermined sex. **Grave goods:** No grave goods. **Dating:** Direct InCal 20 radiocarbon dated to 129-244 cal CE (1840±20 BP, PSUAMS-13134). **Ancestry summary:** Iberia_IA.

#### 1.23 Tarragona, Reus / La Canonja, Antigons

Contact person(s): Josep María Vergès

**Brief settlement history:** This site consists of a Roman villa, founded around the mid-2nd century BCE. It likely belonged to a significant person or family from *Tarraco*. It has three distinct parts. The central section features a luxurious residential area with a magnificent nymphaeum decorated with marble statues of the goddess Cybele and the god Bacchus, among others. In the northern section, there's a pottery workshop with two large kilns, where amphorae, jars, and dolia were produced. Also, the presence of construction materials like roof tiles and antefixes, indicate this was produced here too. The southern section is a service and storage area with rooms, dolia, and small deposits and cisterns (some related to winemaking), The sewage system for wastewater is also found in this section. After the 3rd century CE (likely between the 4th and 5th centuries CE) some southern sector rooms and sewers were abandoned or repurposed, and the area began to be used as a necropolis, with various types of inhumation graves. This use continued until the 6th century CE.

**Excavation History:** The first references to this site are found in the book "Tarragona Monumental," published in 1849 by Joan Francesc Albiñana and Andreu de Bofarull, mentioning the discovery of different architectural remains, tombs, and archaeological material. Almost a century later, on October 19, 1951, Salvador Vilaseca reported to the Tarragona Monuments Commission the discovery, during agricultural work, of a Roman-era necropolis. In 1968, Ramon Capdevila made new findings, including millstone remains, and in 1970, along with Salvador Vilaseca, they found mosaic tesserae and a column fragment. In the summer of 1976, Rodolfo Cortés, a professor at the University Delegation of Tarragona at the University of Barcelona, and a group of Archaeology students excavated two Late Roman tombs. Between 1977 and 1978, during the construction of the NANTA feed factory, a salvage excavation was conducted with members of the Archaeological Research Group of the Reus Museum, directed by Lluïsa Vilaseca. During these interventions, various inhumations were excavated, with some in amphorae, and others in tombs made with tegulae, bricks, or stone slabs. These are the graves from which the analyzed remains in this study derive.

Between November 2008 and February 2009, during the expansion of the Mercat del Camp facilities, the Cota 64 company conducted an archaeological intervention in two phases, one directed by Iban Cabrelles and the other by Gema Ortega. This intervention allowed for a better understanding of the northwestern section of the villa's perimeter.

An underground water channel from the early imperial period, built with opus caementicium, was discovered. They also found a ceramic firing kiln, two silos, and 21 graves. The rustic elements fell into disuse around the 3rd-4th centuries, and the area was used as a necropolis during the 5th-6th centuries CE.

The 21 graves found during the excavation in 2009 were classified into four types: inhumations in simple pits with double-pitched tegulae covers (UF 1003, 1034, 1035, 1040, and 1042); inhumations in simple pits with irregular stone covers (UF 1012, 1013, 1014, 1017, 1021, and 1032); inhumations in simple pits with no cover (UF 1015, 1018, 1027, and 1033); and inhumations in cists built with stone slabs (UF 1007, 1009, 1010, 1011, and 1029).

The anthropological study of six individuals exhumed from UF 1009, 1012, 1013, 1014, and 1017, conducted by Joaquin Baixarias, indicates that five were male and one was female, with ages ranging from 30 to 50 years (except for UF 1009, which contained two children aged 3.5 and 4 years). All the adults showed various pathologies (such as osteoarthritis, ossifying myositis, bone wear, and degenerative joint disease) associated with physically demanding work, suggesting that they were likely individuals of low social rank who worked in the rural areas of Tarraco.

**Geographical information:** The Roman villa of Antigons is located in the Camp de Tarragona plain, 4 kilometers from the current coastline and 6 kilometers west of the city of Tarragona. The site is on a small hill between 40 and 50 meters above sea level, on the left bank of the Boella ravine, a seasonal watercourse about 500 meters to the west. This area has traditionally been used for dry farming, primarily vineyards.

**Summary of sampled materials:** The remains studied come from the excavations of 1977-1978, deposited in the Museum of Reus. Unfortunately, the lack of references in the materials deposited in the museum, from where the samples were taken, does not allow us to relate the remains to the scarce information (planimetric and photographic) available from the excavations of 1977-1978.

The data obtained during the 2009 intervention, during the excavation of another sector of the same necropolis, give us a certain context about the typology of the tombs and the anthropological profile of the buried individuals. These remains have not been included in this study because they were not accessible.

**Graves:**

- **Genetic Identifier:** I19795. **Grave Identifier:** MR-3424-A. **Grave Type:** untraceable) **Skeletal information:** untraceable. **Grave goods:** untraceable **Dating:** Archeologically dated to 300-500 CE. **Ancestry summary:** Iberia_IA__+__CNE_EarlyMedieval__+__WestAnatolia_Roman_Byzantine__+__North_African_Punic.

- **Genetic Identifier:** I19796. **Grave Identifier:** MR-3424-B. **Grave Type:** untraceable **Skeletal information:** untraceable. **Grave goods:** untraceable **Dating:** Direct InCal 20 radiocarbon dated to 245-378 calCE (1745±20 BP, PSUAMS-10234). **Ancestry summary:** Iberia_IA__+__WestAnatolia_Roman_Byzantine__+__North_African_Punic.

- **Genetic Identifier:** I19797. **Grave Identifier:** MR-3424-C. **Grave Type:** untraceable **Skeletal information:** untraceable. **Grave goods:** untraceable **Dating:** Archeologically dated to 300-500 CE. **Ancestry summary:** Excluded of genome wide analysis due to low coverage.

- **Genetic Identifier:** I19798. **Grave Identifier:** MR-3424-D. **Grave Type:** untraceable **Skeletal information:** untraceable. **Grave goods:** untraceable **Dating:** Archeologically dated to 300-500 CE. **Ancestry summary:** Iberia_IA__+__WestAnatolia_Roman_Byzantine.

- **Genetic Identifier:** I19799. **Grave Identifier:** MR-3424-E. **Grave Type:** untraceable **Skeletal information:** untraceable. **Grave goods:** untraceable **Dating:** Archeologically dated to 300-500 CE. **Ancestry summary:** Iberia_IA__+__CNE_EarlyMedieval__+__WestAnatolia_Roman_Byzantine__+__North_ African_Punic.

- **Genetic Identifier:** I35495. **Grave Identifier:** MR-3424-H. **Grave Type:** untraceable. **Skeletal information:** untraceable. **Grave goods:** untraceable **Dating:** Direct InCal 20 radiocarbon dated to 594-650 calCE (1440±20 BP, PSUAMS-10235). **Ancestry summary:** Iberia_IA__+__WestAnatolia_Roman_Byzantine__+__North_African_Punic.

#### 1.24 Tarragona, Cambrils

##### 1.24.1 Els Masos

Contact person(s): Josep Maria Vergès

**Brief settlement history:** The area where the site is located has traditionally been used for dry farming. The remains of a residence were found, which likely served as the center of an agricultural estate. It had a luxurious residential section with a peristyle, private baths, and decorations including paintings, mosaics, and marble stonework. The settlement was founded in the mid-1st century CE and lasted until the last quarter of the 3rd or first quarter of the 4th century, at which point the villa was abandoned.

**Excavation History:** In October 1999, under the direction of Ester Ramon from CODEX-Arqueologia i Patrimoni, an archaeological intervention took place during the construction of a water pipeline to supply the Ponent residential areas (Camarasa, 2002; Ramon, 1999). During this intervention, several architectural structures were identified, highlighting the significance of the settlement. In 2005 and 2006, during the construction of the A-34 highway, another archaeological intervention was conducted, directed by Pèir Còts from SICOM INFORMÀTICA SL, involving the excavation of an area of about 20,000 square meters (Cóts, 2005b, 2005a).

**Geographic Information:** It's located on the coastal plain, about 20 kilometers southwest of Tarragona, on the left bank of the Riudecanyes stream, 2 kilometers from the current coastline, within the municipality of Cambrils (Baix Camp).

**Summary of sampled materials:** We sampled three individuals, deposited in the Museum of Cambrils, who correspond to a father-mother-daughter trio. Unfortunately, the excavation report has not been made available, which is why the details of the grave(s) are unknown.

**Graves:**

- **Genetic Identifier:** I29524. **Grave Identifier:** 3012. **Grave Type:** untraceable. **Skeletal information:** untraceable. **Grave goods:** untraceable.**Dating:** Archeologically dated to 200 BCE - 500 CE. **Ancestry summary:** Iberia_IA__+__WestAnatolia_Roman_Byzantine__+__North_African_Punic.

- **Genetic Identifier:** I29525. **Grave Identifier:** 3076. **Grave Type:** untraceable.**Skeletal information:** untraceable. **Grave goods:** untraceable. **Dating:** Archeologically dated to 200 BCE-500 CE. **Ancestry summary:** Iberia_IA__+__WestAnatolia_Roman_Byzantine__+__North_African_Punic.

- **Genetic Identifier:** I29526. **Grave Identifier:** 3256. **Grave Type:** untraceable. **Skeletal information:** untraceable. **Grave goods:** untraceable. **Dating:** Archeologically dated to 200 BCE-500 CE. **Ancestry summary:** Iberia_IA__+__WestAnatolia_Roman_Byzantine__+__North_African_Punic.

##### 1.24.2 La Mineta

Contact person(s): Josep Maria Vergès

**Brief settlement history:** During the Roman period, the inmediate surroundings of this rescue excatvations are located inside of the limints of the Aeger Tarraconensis (territory under the influence of the socio-economics limits of *Tarraco*). This Late Antiquity necropolis where 33 graves have been recovered. The inhumations were done in amphorae, double-pitched tegulae, stone slab covers and simple pits. Were mostly oriented NW-SE. It is likely that this necropolis is related to the remains of a villa located nearby, for which we do not have precise data, as it has not been excavated.

**Geographic Information:** La Mineta is located on the coastal plain, at 30 meters above sea level and 1.5 kilometers from the current coastline, in an area bordered on either side by the Alforja stream and the Mare de Déu del Camí ravine, near the town of Cambrils (Baix Camp), about 17 kilometers southwest of Tarragona. The site's name comes from the presence of an old water mine in the area. However, the site has traditionally been in a region dedicated to dry farming.

**Excavation History:** In the year 2000, during the excavation of a stormwater drain, 15 Late Antiquity graves were uncovered. In 2004, due to a new phase of urbanization in the area, an archaeological survey was conducted by digging trenches. During this process, 18 new graves were found.

The graves excavated during the 2000 excavation were simple pits with double-pitched tegulae structures or stone slabs and other reused materials. The skeletons were oriented NW-SE (head to foot) and had no grave goods. It is likely that this necropolis is related to the remains of a villa located nearby.

The 18 graves uncovered during the 2004 excavation were mostly oriented NW-SE. The inhumations were done in amphorae in two cases (UF5 and UF7), with double-pitched tegulae in one case (UF18), with stone slab covers in eight cases (UF1, UF2, UF8, UF9, UF10, UF15, UF16, UF17), and in simple pits in seven cases (UF6, UF11, UF12, UF13, UF14) (García, 2004).

**Summary of sampled materials:** The samples studied come from the 2000 intervention, for which we do not have an excavation report, which prevents us from knowing in which type of tomb each individual was found.

**Graves:**

- **Genetic Identifier:** I29575. **Grave Identifier:** UE223. **Grave Type:** untraceable. **Skeletal information:** untraceable. **Grave goods:** untraceable. **Dating:** Archeologically dated to 200 BCE-500 CE. **Ancestry summary:** Iberia_IA__+__WestAnatolia_Roman_Byzantine__+__North_African_Punic.

- **Genetic Identifier:** I29628. **Grave Identifier:** UE211. **Grave Type:** untraceable. **Skeletal information:** untraceable. **Grave goods:** untraceable. **Dating:** Archeologically dated to 200 BCE-500 CE. **Ancestry summary:** Iberia_IA__+__WestAnatolia_Roman_Byzantine__+__North_African_Punic.

- **Genetic Identifier:** I29629. **Grave Identifier:** UE224. **Grave Type:** untraceable. **Skeletal information:** untraceable. **Grave goods:** untraceable.**Dating:** Archeologically dated to 200 BCE -500 CE. **Ancestry summary:** Iberia_IA__+__WestAnatolia_Roman_Byzantine__+__North_African_Punic.

##### 1.24.3 Voltants de la Torre Bargallona

Contact person(s): Josep Maria Vergès

**Brief settlement history:** The site, although it shows evidence of activity from the 14th to the 18th centuries, is notable for a large field of silos, likely used to store agricultural surplus, decommissioned between the 5th and 8th centuries CE. It also contains sand extraction pits, water wells, combustion structures, and 10 human inhumations. The field of silos and most structures are probably connected to a Roman settlement located about 250 meters away. The settlement is known but has not been excavated.

Nearly 300 excavated structures were found, divided into three main groups based on their shape. The first group, with irregular shapes, was interpreted as sand extraction pits. The second group, with a pear-like shape, had openings between 60 and 150 cm in diameter and a depth between 50 and 180 cm, suggesting they were silos. The third group, likely wells, had openings about 70 cm in diameter on average and indefinite depth (excavated only to a depth of 2.30 meters).

**Excavation History:** Between April and October 2006, due to the construction project of the A-34 highway in the Cambrils - Vilaseca - Salou section (from PK 1143 to PK 1151), a preventive archaeological intervention was carried out, directed by Montse Corominas Vidal from ATICS SL. In the first phase, the area of archaeological interest was defined through archaeological survey trenches. After marking the boundaries and dividing the site into three areas (Camp 1, 2, and 3), an extensive excavation was conducted (Vidal, 2007).

**Geographic Information:** The site known as Voltants de la Torre Bargallona is located on the coastal plain, about 15 kilometers southwest of Tarragona and 2 kilometers from the current coastline, within the municipality of Cambrils (Baix Camp). The area where the site is located has traditionally been used for dry farming.

**Summary of sampled materials:** The site's chronology can be divided into two distinct periods. The first period is represented by 143 silos containing fragments of African amphora (Keay 62A and Late Roman C), African Red Slip Ware (Hayes 91 and Hayes 61), and some local imitations of imported ceramics, indicating the site's abandonment between the 5th and 8th centuries CE. The second period is comprised of 10 structures with fragments of Catalan blue and green and manganese pottery, and medieval gray ceramics, suggesting that these structures were abandoned between the 14th and 18th centuries. A total of 41 structures have not been dated due to a lack of material evidence. The remains of 10 individuals were found, all inhumations. Two of the inhumations located in Camp 1, UE 23.01 and UE 23.02, correspond to a perinatal individual and an infant (3-6 months), respectively, and were buried in amphorae. Individual 23.01 was placed inside an Eastern amphora, LRA, dated between 380 and 650 CE. The amphora of individual 23.02, found in a supine position and oriented NW-SE, is dated to the late Roman period (Vidal, 2007).

In Camp 1, a skull (TBAR R-30 Camp 1) belonging to a 10-13-year-old individual was found. The skull was discovered in the filling of a circular cut, measuring 1.46 meters in diameter and 2.15 meters deep, along with the skeletons of five dogs, construction ceramic materials, and fragments of common ware with reduced firing.

In Camp 2, two graves were found in simple pits (graves 23 and 41). Grave 23, oriented N-S, contained the remains of two individuals (23A, aged 13-14 years, and 23B, aged 15-16 years), one on top of the other in a supine position. When individual 23A was placed, the skull and jaw of 23B were shifted, suggesting that 23B was already in a skeletal state. Grave 41, oriented SW-NE, contained the remains of three individuals (41A, aged 25-35 years, 41B, of undetermined age, and 41C, aged 1-1.5 years). The largest bones from the first individual were collected and placed between the legs of the second individual to make space, while the third individual was placed on top of the second, both in a supine position. In Camp 3, two individuals were found: TBAR IND 15A-I, aged 14-16 years, in a prone position without any apparent pit or funerary structure, and TBAR 15B.01 (corresponding to TBAR IND.2), aged 14-15 years, in a supine position, buried in a silo. The adults and subadults showed enthesopathies on their forearms and legs, likely from continued physical labor. The anthropological study of individuals from Camp 1, except for TBAR R-30 Camp 1, and those from Camp 2 was carried out by Thaïs Fadrique from Antropòlegs. The individuals from Camp 3 and the previously mentioned TBAR R-30 Camp 1 were analyzed by Thaïs Fadrique, Alba Núñez, and Assumpció Malgosa from the Osteobiography Research Group (GROB) of the Autonomous University of Barcelona (Vidal, 2007).

**Graves:**

- **Genetic Identifier:** I29537. **Grave Identifier:** Ind15A-1. **Grave Type:** prone decubitus position without any apparent pit or funerary structure. **Skeletal information:** Probably female, between 14-16 years. **Grave goods:** No grave goods **Dating:** Archeologically dated to 400 CE-700 CE. **Ancestry summary:** Iberia_IA__+__WestAnatolia_Roman_Byzantine.

- **Genetic Identifier:** I29538. **Grave Identifier:** Ind23A. **Grave Type:** doble burial in simple pit, oriented S-N. Individual in supine decubitus positioned on top of another body deposited some time before. **Skeletal information:** Female, 13-14 years **Grave goods:** She wore an earring **Dating:** Archeologically dated to 400-700 CE. **Ancestry summary:** Excluded of the genome wide analysis due to low coverage. **Additional information:** dental hypoplasia

- **Genetic Identifier:** I29539. **Grave Identifier:** Ind23B. **Grave Type:** doble burial in simple pit, oriented S-N. individual in supine decubitus. The skull was displaced when the second individual was placed. **Skeletal information:** Female, 15-16 years **Grave goods:** No grave goods **Dating:** Archeologically dated to 400-700 CE. **Ancestry summary:** Iberia_IA__+__WestAnatolia_Roman_Byzantine__+__North_African_Punic. **Additional information:** slight dental hypoplasia.

- **Genetic Identifier:** I29540. **Grave Identifier:** Ind41A. **Grave Type:** triple burial in simple pit, oriented SW-NE. The skeletal remains of this individual, the first to be buried, were collected and placed on top of a second corpse. **Skeletal information:** Male, adult, 25-35 years **Grave goods:** No grave goods **Dating:** Archeologically dated to 400-700 CE. **Ancestry summary:** Iberia_IA__+__WestAnatolia_Roman_Byzantine__+__North_African_Punic. **Additional information:** enthesopathies in the forearms and legs, resulting from continued heavy physical exertion

- **Genetic Identifier:** I29634. **Grave Identifier:** TBAR'06; R-30, Camp I. **Grave Type:** isolated skull in a pit **Skeletal information:** undetermined sex, 10-13 years. **Grave goods:** No grave goods **Dating:** Archeologically dated to 400-700 CE. **Ancestry summary:** WestAnatolia_Roman_Byzantine__+__North_African_Punic. **Additional information:** dental hypoplasia and *criba orbitalia.*

#### 1.25 Tarragona, La Canonja, Villa Romana dels Castellets

Contact person(s): Moisés Díaz García, Jordi Morera, Tatiana Piza Ruíz, Josep F. Roig Pérez,

**Brief settlement history:** The archaeological works revealed a coastal villa, with an area of about 7.500 $m^2$. We identified several Roman structures, with a chronology from the first century AD to the late 7th century CE.

**Excavation History:** The archaeological excavations were undertaken in two moments, first between 2013-2014 and later in 2017-2018. During the first excavation most of the structures (pars urbana, pars rustica, pars fructuaria and a little burial place) were identified. During the second excavation, structures associated with a funerary area and a torcularium were discovered.

**Geographic Information:** The site is currently located in the Chemical Park of Bayer Material Science, S. L. Previously it was in an agricultural land. In the roman period the villa was in a plain fertile soil with mineral water, near the sea and the capital city of Tarraco.

**Summary of sampled materials:** Archaeological work confirmed a *villae* with an urban and architectural evolution that spanned from the 1st century AD to the end of the 7th century AD. The foundation of the *uilla* should be placed, in the Augustus era. Even so, it'll not be until the end of the second century AD that this coastal *villae* will be equipped with all those architectural and residential elements that will define it. It'll be at this time that a residential area will be created with a distribution courtyard on which the different documented areas will pivot luxurious rooms with mosaic flooring, *cubiculae* and a *balneum* for private use with the *praefurnium* outside the building. Another area that is being built at this time corresponds to a wine production area, which was characterized by containing all those spaces that define it: work areas, press area and a wine cell occupied by a series of *dolia defossa*. The splendor of this settlement ends abruptly around 260-270 AD, probably linked to the period of the Frankish raids. A magnificent waste dump is a silent witness to this moment. Following this moment, the *uillae* falls into a process of abandonment that will last well into the 5th century AD, at which time it will be subject to structural reforms and reuse of materials and architectural areas, which will result in a new residential *uilla* with a central courtyard or garden and three possible distribution arms that separate functional spaces. Of all these, the construction of a new *balneum* stands out, as well as other areas of representation paved in opus signinum and with parts decorated with paintings. At this time there is also a production area with pipelines and tanks lined with opus signinum, which inform us about the exploitation of a dependent fundus. The reuse of materials and architectural areas will be a constant in later chronological phases, but new ones will also be created, as evidenced by the documentation of different wall structures, a fireplace and the construction of a canal kiln with a chronology of the second half of the 7th century AD. At this time, the construction of silos is also noteworthy, which would indicate that we are facing a rural settlement, where cereal production played an important role. We date the last moment of occupation to the end of the 7th century AD.
As for the exhumed individuals, two (UE's 2576 and 2578) were identified inside individual silos; two more (UE 3195 and 4244) were located in isolation; and the rest shared a small necropolis composed of slab cists. It is worth saying that individuals UE 4216 and 4217 shared the same burial space.

**Graves:**

- **Genetic Identifier:** I28364. **Grave Identifier:** UE 10039; UF 3. **Grave Type:** unknown **Skeletal information:** Unkown sex or age. **Grave goods:** no grave goods. **Dating:** Archeologically dated to 300-500 CE. **Ancestry summary:** Iberia_IA__+__WestAnatolia_Roman_Byzantine__+__North_African_Punic.

- **Genetic Identifier:** I29542. **Grave Identifier:** LCB13-UE4217. Grave Type human found inside a container of stone slabs. **Skeletal information**: Adult, 40-45 years old, Female. Lower jaw. **Grave goods:** no grave goods. **Dating:** Archeologically dated to 200-500 CE. **Ancestry summary:** Iberia_IA__+__WestAnatolia_Roman_Byzantine__+__North_African_Punic. **Additional information:** Cervical and dorsal arthritis. Schmorl's nodes on thoracic vertebrae D4-D12 and lumbar vertebrae L1-L4. Calcification of yellow ligaments in the dorsal vertebrae. Osteochondritis dissecans in the right mandibular condyle.

- **Genetic Identifier:** I29543. **Grave Identifier:** LCB13-2576. **Grave Type:** inside a silo (with I29545, without anatomical connection). **Skeletal information:** Adult, 20-25 years old, Male. Lower jaw. **Grave goods:** no grave goods. **Dating:** Archeologically dated to 200-500 CE. **Ancestry summary:** Iberia_IA__+__WestAnatolia_Roman_Byzantine__+__North_African_Punic. **Additional information:** Missing limbs. Infectious process in the palate. Periostitis on tibiae and fibulae. Possible osteoma on the lambda.

- **Genetic Identifier:** I29545. **Grave Identifier:** LCB13-2578. **Grave Type:** inside a silo (with I29543, without anatomical connection) **Skeletal information:** Adult, 25-35 years old, Male. Lower jaw. **Grave goods:** no grave goods. **Dating:** Archeologically dated to the sixth century CE. **Ancestry summary:** Iberia_IA__+__WestAnatolia_Roman_Byzantine__+__North_African_Punic. **Additional information:** Missing lower limbs. Signs compatible with tuberculosis (Pott's disease). Lesions on vertebrae D12, L2, L1. Fusion of vertebral arches D10-D11 and D12-L1. Calcification on an intercostal rib. Abnormal compression on the femur (possibly related to tuberculosis). Callus on the distal third of the left ulna. Callus on the proximal phalanx V of the right hand. Mixed human and faunal remains.

- **Genetic Identifier:** I29546. **Grave Identifier:** LCB13-3195. **Grave Type:** simple fossa with an elliptical plant. **Skeletal information:** Adult, 45-55 years old, Female. Lower jaw. **Grave goods:** no grave goods. **Dating:** Archeologically dated to the fifth century CE. **Ancestry summary:** Iberia_IA. **Additional information:** Considerably altered by construction processes from the so-called "second terms" dated around the mid-5th century CE.

- **Genetic Identifier:** I29547. **Grave Identifier:** LCB13-4171. **Grave Type:** human found inside a container of stone slabs. **Skeletal information:** Adult, >60 years old, male. Lower jaw. **Grave goods:** no grave goods. **Dating:** Archeologically dated to 200-500 CE. **Ancestry summary:** Iberia_IA__+__WestAnatolia_Roman_Byzantine__+__North_African_Punic. **Additional information:** Generalized arthritis in the spine, knees, shoulders, and feet. Trauma to the right parietal. Calcification of the Achilles tendon on both heels. Large fistula in dental piece 23. Almost all alveoli reabsorbed due to tooth loss during life. Mixed human and faunal remains.

- **Genetic Identifier:** I29548. **Grave Identifier:** LCB13-4212. **Grave Type:** Human found inside a container of stone slabs. **Skeletal information:** Adult, 60-70 years old, Male. Lower jaw. **Grave goods:** no grave goods. **Dating:** Archeologically dated to 200-500 CE. **Ancestry summary:** Iberia_IA__+__WestAnatolia_Roman_Byzantine__+__North_African_Punic. **Additional information:** Ossification of the costal cartilage of the first rib fused with the manubrium. Vertebral arthritis. Schmorl's nodes on D12, L1-L4. Arthritis in the hands (metacarpals I, II right and proximal phalanx III left). Calcification of the Achilles tendon on both heels. Arthritis in the hallux of the feet. Arthropathy in the knees and hip joints.

- **Genetic Identifier:** I29549. **Grave Identifier:** LCB13-4216. **Grave Type:** Undetermined. **Skeletal information:** Adult, 55-65 years old, Female. Lower jaw. **Grave goods:** no grave goods. **Dating:** Archeologically dated to 200-500 CE. **Ancestry summary:** Iberia_IA__+__WestAnatolia_Roman_Byzantine__+__North_African_Punic. **Additional information:** Costoclavicular sulcus on both clavicles. Strong calf and *gluteus minimus* medius insertion on femurs. Mid-insertion of the flexor digitorum longus on tibiae. Strong insertion of obliques, semimembranosus, and adductors on the pelvis. Osteophytes on the sacrum at S1. Cervical, dorsal, and L5 arthritis. Schmorl's nodes on D3, D7, and D8. Enthesopathy of the Achilles tendon on both heels. Possible trauma to the right parietal.

- **Genetic Identifier:** I29550. **Grave Identifier:** LCB13-4219. **Grave Type:** Human found inside a container of stone slab. **Skeletal information:** Adult, 35-45 years old, Male. Lower jaw. **Grave goods:** no grave goods. **Dating:** Archeologically dated to 200-500 CE. **Ancestry summary:** Excluded from the genome wide analysis due to low coverage. **Additional information** Costoclavicular sulcus on both clavicles. Sacralization of L5. Vertebral arthritis. Hernias on D6, D7, and D10. Schmorl's nodes on D8 and D9. Calcification of the Achilles tendon on both heels. Consolidated fracture on the distal phalanx V of the right foot. Secondary arthritis at the fracture site between the proximal and medial phalanx V of the right foot.

- **Genetic Identifier:** I29552_d. **Grave Identifier:** LCB13-4244. **Grave Type:** Human found inside a container of stone slabs. **Skeletal information:** Adult, 30-35 years old, Female. Lower jaw. **Grave goods:** no grave goods. **Dating:** Archeologically dated to 4th to 6$^{th}$ centuries CE. **Ancestry summary:** Iberia_IA__+__WestAnatolia_Roman_Byzantine__+__North_African_Punic. **Additional information:** Cribra femoralis on both femurs.

#### 1.26 Tarragona, La Selva del Camp, Barranc de Sales

Contact person(s): Josep Maria Vergès

**Brief settlement history:** At the Barranc de Sales site, remains of a *figlina* (ceramic production center) with an attached necropolis were found. The *figlina* began production after the first half of the 1st century BCE, or during the first century of the 1st century CE, and remained active until the late 3rd or early 4th century CE, with some minor production activity possibly continuing until the late imperial period. The first documented inhumations occurred at the end of the early imperial period, while the ceramic production center was still in full operation, and burials continued until the 5th century CE.

Among the structures associated with the ceramic production center, there were three rectangular-shaped settling tanks, with bases and walls made of tegulae, used for clay decantation. There was also a circular mixing trough, between 6 and 6.5 meters in diameter and 35 cm in preserved depth, with an opus signinum floor over a *rudus* base made of cobbles partially bound with lime mortar. Additionally, two large storage tanks for liquids were built with opus *caementicium* walls and lined with opus signinum. The site also had three kilns: one circular with a 3.5-meter diameter and two square-shaped kilns, with the best-preserved measuring about 4 x 4 meters. The production at this center focused on making construction elements such as tegulae, *imbrices, tubuli*, bricks with both negative and positive interlocking features, circular bricks, and antefixes.

The inhumation necropolis contained 48 funerary units, divided into two groups: Group A with 23 units and Group B with 20. There were also five isolated and scattered funerary units throughout the area. Group B is adjacent to the southern edge of the *figlina* structures, while Group A is in the same area but slightly farther, about 10 meters away. Both groups exhibited different orientations, with the most common being SW-NE, and varying burial types and ages of those buried. Regarding the types of funerary structures, the most common was inhumation in amphora containers (15 cases), followed by inhumation in double-pitched tegulae structures (8 confirmed cases and 6 possible), and pits with walls made of stones bound with lime mortar (4 cases). The only observable difference is that all four stone-built tombs were in Group B, suggesting, though not confirmed, a potential class distinction between the two groups.

Most of the graves are located outside the manufacturing area, except for inhumations 47 and 49, which are within the pottery zone, and inhumation 7, whose pit cuts through a floor related to the workshop area, indicating that this burial was done after the original manufacturing function had ended.

**Excavation History:** The existence of the site was first noted in a 1991 university paper by Andreu Ollé and Josep Vallverdú. Archaeological excavations took place in 2004 and 2006 during the expansion of the industrial area in La Selva del Camp. The first phase in 2004, mainly for delimitation, was conducted by Nemesis SCCL under Jaume Vilalta's direction. The second phase, from 2005 to 2006, completed the delimitation and carried out extensive excavation by Codex under Marta Bru's direction.

**Geographic Information:** The Barranc de Sales site is located one kilometers northeast of the town of Selva del Camp, on the edge of the Camp de Tarragona plain, at the foot of the first foothills of the pre-coastal mountain range, near the Selva stream, a seasonal watercourse.

**Summary of sampled materials:** The samples analyzed come from the funerary units (UF) 15 and 17, from sector A, and 44 and 48, found in sector B. The individual from UF15 was buried in an amphora container, UF44 is a grave with side walls made of *opus caementicium* and covered with tegulae, and individual of UF48 is covered with double-pitched tegulae. All three were found in decubitus position, and no grave goods were present. We have no information about the grave of UF17.

**Graves:**

- **Genetic Identifier:** I29531. **Grave Identifier:** BS-UE15. **Grave Type:** amphora container. **Skeletal information:** undetermined sex and age. **Grave goods:** No grave goods. **Dating:** Archeologically dated to 200 BCE-500 CE. **Ancestry summary:** Iberia_IA__+__WestAnatolia_Roman_Byzantine__+__North_African_Punic. **Additional information:** Individual in a supine position: legs extended and parallel, with arms stretched alongside the body or slightly bent, resting one or both hands on the pelvis or abdomen.

- **Genetic Identifier:** I29532. **Grave Identifier:** BS-UF17. **Grave Type:** untraceable. **Skeletal information:** undetermined sex and age. **Grave goods:** No grave goods. **Dating:** Archeologically dated to 200 BCE - 500 CE. **Ancestry summary:** Excluded from de genome wide analysis due to low coverage.

- **Genetic Identifier:** I29535. **Grave Identifier:** BS-UF44. **Grave Type:** Grave with side walls made of *opus caementicium* and covered with tegulae. **Skeletal information:** undetermined sex and age. **Grave goods:** No grave goods. **Dating:** Archeologically dated to 200 BCE -500 CE. **Ancestry summary:** Iberia_IA__+__WestAnatolia_Roman_Byzantine. **Additional information:** Individual in a supine position: legs extended and parallel, with arms stretched alongside the body or slightly bent, resting one or both hands on the pelvis or abdomen.

- **Genetic Identifier:** I29574. **Grave Identifier:** BS-UF48. **Grave Type:** covered with double-pitched tegulae. **Skeletal information:** undetermined sex and age. **Grave goods:** No grave goods. **Dating:** Archeologically dated to 200 BCE -500 CE. **Ancestry summary:** Excluded from de genome wide analysis due to possible contamination. **Additional information:** Individual in a supine position: legs extended and parallel, with arms stretched alongside the body or slightly bent, resting one or both hands on the pelvis or abdomen.

##### 1.27 Tarragona, Manuel de Falla

Contact person(s): Moisés Díaz García, Josep F. Roig Pérez, Josep María Vergès

**Brief settlement history:** The archaeological works revealed a several port warehouses, one via and another suburban structures, with a chronology from the fifth century AD to the late sixth century CE.

**Excavation History:** The archaeological excavations were undertaken in different moments, between 2017-2020. But was in the 2018 when a funerary area was discovered.

**Geographic Information:** The site is in south-west of Tarragona and is bounded to the north by the old Tabacalera building, to the east by number 12 of the same street, to the south by Torres Jordi street, and to the west by the Passeig de la Independència.

**Summary of sampled materials:** The oldest documented evidence would be related to the construction of large structures, which we can relate to port warehouses. Articulating these structures, it is identified at the southeast end of the site, part of a stretch of road in a north-east to south-west direction, reformed during the second half of the 5th century AD. The preserved part has a direction towards the west (towards the Tulcis River), and measures about 10 m in length by a width that oscillates between 1.50 m in the east and 0.80 m in the west.

Once the harbor ships of the previous phase had been amortized, a large building with a rectangular plan was built, of which only its southern side and part of the western boundary could be documented. This building, divided into three areas, had a minimum length of 14.23 m and a minimum width of 4.96 m. In posterity, the advanced second half of the 5th century AD, this building will be expanded.

After a period of abandonment and demolition (dated very late 5th century AD - early 6th century AD) a small cemetery area was identified with six Funeral Units - two of them despoiled - and all of simple type.

**Graves:**

- **Genetic Identifier:** I29631. **Grave Identifier:** UF1. **Grave Type:** Simple pit, filled with construction materials (tiles, dolia, stones). **Skeletal information:** Adult male, 40-45 years old. **Grave goods:** No grave goods. **Dating:** Archeologically dated to 200 BCE - 500 CE. **Ancestry summary:** CNE_EarlyMedieval__+__North_African_Punic. **Additional information:** Estimated height between 169.86 and 172.05 cm, based on the maximum length of the left femur. The skeleton is gracile, with marked muscular development in the medial third of the humeral diaphysis. Light signs of arthritis bilaterally affect the shoulder, elbow, and hip joints. Schmorl's nodes are present on lumbar vertebrae L2 and L3, and ligament calcification is present on the thoracic vertebrae. The right femur shows a third trochanter, and both tibiae exhibit non-pathological external torsion, possibly indicative of equestrian activity. The right tibia and first phalanx of the right foot show osteochondritis lesions. The acetabulum shows a small groove with porosity, possibly linked to the pelvic ligament system. Cervical vertebra C3 shows an abnormal trabecula, suggesting an extradural lesion. Dental examination reveals three carious lesions, generalized alveolar recession, and dental calculus buildup. The sagittal suture is almost entirely obliterated.

- **Genetic Identifier:** I29632. **Grave Identifier:** UF3. **Grave Type:** Simple pit filled with soil. **Skeletal information:** Partial skeletal remains of a non-adult individual aged 3-4 years. **Grave goods:** No grave goods. **Dating:** Archeologically dated to 200 BCE - 500 CE. **Ancestry summary:** Iberia_IA.

- **Genetic Identifier:** I29633. **Grave Identifier:** UF4. **Grave Type:** Adjacent to a wall on the east side of the plot, buried in a simple pit. The fill contained earth, glass fragments, and iron pieces. **Skeletal information:** Non-adult individual aged 12±1 years, probably female. **Grave goods:** No grave goods. **Dating:** Archeologically dated to 200 BCE - 500 CE. **Ancestry summary:** Iberia_IA. **Additional information:** The skeleton shows several notable alterations: periostitis with thickening, porosity, and lamellar layers affecting the diaphyseal and metaphyseal parts of the long bones (distal right humerus, proximal right and left ulna and radius, distal left ulna and radius, diaphyses of right and left tibiae, diaphyses of right and left fibulae, metacarpals and phalanges of hands, metatarsals, and phalanges of feet). Bilateral femoral cribra, bilateral thickening of the tibiae, and curvature of the fibulae are present. Anomalous striation is observed on the anterior side of the patellae. Similar periosteal lesions affect the long bones of the hands and feet. These alterations are macroscopically and distributionally consistent with thalassemia minor, a congenital hemoglobin disorder, ruling out other infectious etiologies. Further paleopathological analysis, including genetic testing, is recommended to confirm the diagnosis.

#### 1.28 Tarragona, Mas Gassol, Alcover

Contact person(s): Josep Maria Vergès

**Brief settlement history:** The site has two distinct areas. On one hand, a necropolis with six funerary structures and human remains from ten individuals, and on the other hand, about 10 meters away, the remains of an opus signinum deposit and, nearby, the rubble of a dry-stone structure, interpreted as the remains of a possible livestock enclosure (Canyellas et al., 1996).The complex is dated to between the 3rd and 5th centuries CE, though the recovered materials do not allow for greater chronological precision..

**Excavation History:** In December 1991, during the construction of the Mas Gassol urbanization in Alcover, excavators uncovered and largely destroyed a Roman-era necropolis. After this was reported by Josep Maria Vergès to the relevant authorities, the Archaeology Service of the Generalitat de Catalunya commissioned him and Lluís Piñol to conduct an urgent archaeological intervention, which took place from January 1 to January 10, 1992.

**Geographic Information:** The site is located one kilometer northeast of the town of Alcover, at the edge of the Camp de Tarragona plain, right at the base of the first elevations of the prelitoral mountain range, near the Font Major ravine, which is a constant water source that still supplies part of the town's water today.

**Summary of sampled materials:** Although the tombs were severely damaged, preserving only partial remains, their main characteristics could be documented. They share a common NW-SE orientation, with individuals buried in primary position having their heads toward the NW side. In the construction of all tombs, except for UE 104, built entirely with tegulae, and UE 108, where the individual was buried in a wooden coffin, limestone slabs from Muschelkalk were used for the base and some sides (in the case of UE 111, a tegula with trimmed wings was used), while the remaining sides were made with small stone walls bound with lime and sand mortar. These tombs likely had some form of covering or overhead structure, also made with stone and mortar, but due to the level of destruction, their characteristics are unknown. DNA preservation of these samples was minimal. The reported individual was published in Olalde et al. (2019), whose tomb was destroyed by building heavy equipment before the rescue excavation started. It was re-sequenced for this paper. however, it presented signals of contamination.

**Graves:**

- **Genetic Identifier:** I6491_enhanced.AG. **Grave Identifier:** MGA'92-Resta III. **Grave Type:** Undetermined. **Skeletal information:** Undetermined sex and age. **Grave goods:** No grave goods. **Dating:** Archeologically dated to 200-500 CE. **Ancestry summary:** Excluded from the analysis because of contamination signals. **Additional information:** Resequenced from Olalde et al 2019.

#### 1.29 Tarragona, Vilardida

Contact person(s): Adrià Cubo Córdoba, Jordi Morera

**Brief settlement history:** The Vilardida site shows a timeline of occupation from the Iberian period to contemporary times. This spans over twenty centuries, with particularly interesting phases from the Iberian period to late antiquity. The first major chronological horizon of the site is in the late Iberian or Roman Republican period (3rd-1st centuries BC). Archaeological remains from this period indicate a rural settlement identified by partially excavated areas and some storage structures related to a small Iberian rural farm. The second chronological horizon corresponds to the early Imperial period between the turn of the era and the late 3rd century AD. This is likely associated with a villa, not well-defined by archaeological work, probably located under the current town of Vilardida. The third chronological horizon of the site is between the late Empire and late antiquity and is the most well-defined. Remains from this period indicate a rural settlement with a complex historical evolution, showing different phases of occupation between the 5th and 8th centuries AD.

**Excavation History:** The archaeological intervention at the Vilardida site took place between March 2018 and July 2019, covering an area of about 3.5 hectares. This work was done as part of the improvement of the C-51 road near the municipalities of Montferri and Vila-rodona (Arrayás Morales, 2003).

**Geographic Information:** The Vilardida site is located in the small village of the same name, between the municipalities of Monferri and Vila-rodona in the Alt Camp region (Tarragona). Specifically, the excavated area is in the southern part of the village, between the Gaià River bridge and a hill behind Vilardida, on the left bank of the river, at the junction of the C-51 and T-204 roads from Montferri to Vilardida.

**Summary of sampled materials:** The site includes six burials from the Visigothic period. Individual I28357 (UE 5665; UF 46) was a young adult male buried in a double grave with bound hands, dating to 571-651 CE. Individual I28358 (UE 5725; UF 49) was an adult male in a tegulae box, dating to 382-541 CE, with a belt buckle. Individual I28359 (UE 5186; UF 41) was an adult of undetermined sex, 35-40 years old, buried with another person, dating to 550-650 CE. Individual I28360 (UE 5714; UF 48) was an adult male in a pit near a lime kiln, dating to 564-609 CE. Individual I28361 (UE 5544; UF 39) was a 45-50 year old male with health issues, buried with a woman in a double pit, dating to 661-774 CE. Individual I28362 (UE 5179; UF 40) was an adult male, 35-39 years old, buried with two others in a pit, dating to 550-650 CE (Arrayás Morales, 2003).

**Graves:**

- **Genetic Identifier:** I28357. **Grave Identifier:** UE 5665; UF 46. **Grave Type:** Double freely buried grave located at the perimeter of a Visigothic period hut floor (first individual sampled). **Skeletal information:** Young Adult, Male. **Grave goods:** No grave goods found. **Dating:** Direct InCal 20 radiocarbon dated to 571-651 calCE (1450±30 BP, Beta-537541). **Ancestry summary:** Excluded from the genome-wide analysis due to low coverage. **Additional information:** Buried in a double grave, cutting through the layers associated with productive contexts of the villa. Hands are bound. δ13C: -18.2 o/oo IRMS, δ15N: +10.0 o/oo (95.4%)

- **Genetic Identifier:** I28358. **Grave Identifier:** UE 5725; UF 49. **Grave Type:** Burial in a double-sloped tegulae box, unknown orientation. **Skeletal information:** Adult, Male. **Grave goods:** Belt buckle found at the pelvis. **Dating:** Direct InCal 20 radiocarbon dated to 382-541 calCE (1630±30 BP, Beta-537540). **Ancestry summary:** CNE_Early_medieval. **Additional information**: Buried in a deep grave, covered with double-pitched roof tiles, cutting through the layers associated with productive contexts of the villa. δ13C: -18.6 o/oo, δ15N: +7.2 o/oo (64.7%)

- **Genetic Identifier:** I28359. **Grave Identifier:** UE 5186; UF 41. **Grave Type:** Freely buried individual, unknown orientation. **Skeletal information**: Adult, 35-40 years old, undetermined sex. **Grave goods:** No grave goods found. **Dating:** Archeologically dated to 550-650 CE. **Ancestry summary:** Iberia_IA__+__WestAnatolia_Roman_Byzantine__+__North_African_Punic. **Additional information:** Buried in a pit of the villa, along with another individual. δ13C: -18.1 o/oo δ15N: +9.0 o/oo (95.4%)

- **Genetic Identifier:** I28360. **Grave Identifier:** UE 5714; UF 48. **Grave Type:** Burial located just next to I28357 hut floor, W-E orientation. **Skeletal information:** Adult, Male. **Grave goods:** No grave goods found. **Dating:** Direct InCal 20 radiocarbon dated to 564-609 calCE (1460±30 BP, Beta-537539). **Ancestry summary:** Iberia_IA__+__WestAnatolia_Roman_Byzantine__+__North_African_Punic. **Additional information:** Buried at the bottom of a pit associated with a lime kiln, δ13C: -18.7 o/oo IRMS δ15N: +9.4 o/oo (95.4%)

- **Genetic Identifier:** I28361. **Grave Identifier:** UE 5544; UF 39. **Grave Type:** Simple pit burial containing two individuals side by side. First individual sampled, unknown orientation. **Skeletal information:** Adult, 45-50 years old, Male. **Grave goods:** No grave goods found. **Dating:** Direct radiocarbon dated to 661 - 774 cal AD (1289 - 1176 cal BP). **Ancestry summary:** Iberia_IA__+__WestAnatolia_Roman_Byzantine__+__North_African_Punic. **Additional information:** Buried in a double pit cutting through the layers associated with productive contexts of the villa. The sampled male showed a benign tumor on the underside of the jaw, a rib fracture, several herniated discs, and clear signs of malnutrition. Isotopes: Beta - 528295 VMV 39;18 UE 5543 1280 +/- 30 BP IRMS δ13C: -18.3 o/oo IRMS δ15N: +9.5 o/oo (95.4%).

- **Genetic Identifier:** I28362. **Grave Identifier:** UE 5179; UF 40. **Grave Type:** Simple pit burial containing three individuals. Lowest individual sampled, unknown orientation. **Skeletal information:** Adult, 35-39 years old, Male. **Grave goods:** No grave goods found. **Dating:** Archeologically dated to 550-650 CE. Direct Radiocarbon dated to 566 - 654 cal AD (1384 - 1296 cal BP). **Ancestry summary:** Iberia_IA__+__WestAnatolia_Roman_Byzantine. **Additional information:** Buried in a pit of the villa, along with two other individuals. Sampled individual showed the knees fully bent and feet together, likely due to the pressure from the other two individuals above him. Isotopes: Beta - 528298 VMV 39;18 UE 5177 1440 +/- 30 BP IRMS δ13C: -18.1 o/oo IRMS δ15N: +9.4 o/oo (95.4%)

**References:**

Arrayás Morales, Isaías. Morfologia històrica del territorium de Tarraco en època tardo-republicana romana o ibèrica final (ss. III-I aC): cadastres i estructures rurals. Universitat Autònoma de Barcelona,, 2003.

#### 1.30 Toledo, Carranque, Santa María de Abajo

Contact person(s): Virginia García Entero & Sergio Vidal Álvarez.

**Brief settlement history:** Archaeological research indicates that the site was first occupied during the Early Imperial period (1st century CE), though the evidence of structures becomes particularly significant during the Late Roman period (4th and 5th centuries CE). This period is marked by active construction activity, including a productive area associated with oil and wine production, the erection and renovation of the Casa de Materno, adapted to the planning and decorative norms of the elite at that time, the construction of a mausoleum, and, by the late 4th century CE, the building of an imposing palace with interior decoration made from tons of ornamental rocks sourced from major quarries across the Mediterranean and the Iberian Peninsula.

This notable complex began to be dismantled by the mid-5th century CE, when a Late Antique settlement was established over the ruins, leaving little archaeological evidence. Starting in the mid-6th century CE, a large necropolis with more than 100 graves was built around a barely detectable Christian religious building. Following the Visigothic occupation, archaeological work has identified an Andalusian rural settlement with dozens of storage silos. After a period of abandonment, a Christian church and monastery (12th century) were built over the Late Roman palace ruins, with a new necropolis (14th century) associated with it.

Only a small portion of the Late Roman structure survived into the 16th century, serving as a rural chapel. Following the 19th-century disentailment, these walls became a refuge for marginalized populations who occupied the site until the early decades of the 20th century (García-Entero et al., 2017).

**Excavation History:** The Santa María de Abajo site (Carranque, Toledo) was discovered by chance in 1983 when agricultural work revealed a mosaic recognized as belonging to the residential part of a Roman villa. This finding led to archaeological interventions sponsored by the Junta de Comunidades de Castilla-La Mancha from 1985 to the present, in two phases (1985 to 2003 and 2004 onwards). The site opened to the public in 2003 as an Archaeological Park under Toledo's jurisdiction, part of the Castilla-La Mancha Archaeological Parks Network.

During the first phase of research (1985-2003), several buildings were excavated and interpreted as part of the estate of Materno Cinegio, the Praetorian Prefect of Theodosius I in the East. However, this interpretation was seriously challenged within the scientific community. Since 2004, new research has clarified questions about the function and chronology of the excavated buildings through targeted excavation in some sectors and systematic study of material culture elements recovered over the years.

These efforts have revealed a long sequence of occupation at the site, from the early imperial Roman period (1st century CE) to the early 20th century, with significant phases during the Late Roman, Visigothic, Emirate, and medieval Christian periods. This extensive sequence is based on solid stratigraphy and the exhaustive study of recovered materials, particularly ceramics and marble. This research follows an ambitious archaeometric protocol, involving mineralogical and geochemical characterization of lapidary materials (ornamental rocks and construction materials), ceramic pastes, mortars, and brick materials. It also includes physicochemical analysis of residues from work surfaces and ceramic containers, archaeobiological studies (archaeofaunal, carpological, palynological, and anthracological), anthropological analysis of human remains with C14 dating, and genomic and DNA analysis, with researchers from various Spanish and European institutions.

**Geographic Information:** The Santa María de Abajo site (Carranque, Toledo) is located in the center of the Iberian Peninsula, on a broad river terrace on the right bank of the middle course of the Guadarrama River, at the border between the present-day provinces of Toledo and Madrid. The site is positioned at a strategic point at the foot of the XXIV route of the Antonine Itinerary, which allowed north/south communication between the two Mesetas through the Fuenfría pass in the Sierra del Guadarrama. It also connected east/west, serving as an essential route for interior communications during the Roman era. This road remained in use until the conquest of Toledo in 1085, after which it was replaced by other roads and routes that emerged due to the new political and socio-economic circumstances from that point onward (García-Entero et al., 2017).

**Summary of sampled materials:**
The samples come from graves excavated during the 2009 and 2010 campaigns(García-Entero et al., 2017). All belong to a Visigothic-era necropolis, which contains at least a hundred graves excavated over several campaigns. The necropolis was established over the ruins of a Late Roman palace, which had been repurposed to house a Christian Basilica, around which the necropolis developed. There is little information about the basilica's interior decorative elements (García-Entero et al., 2017).

**Graves:**

- **Genetic Identifier:** I15419. **Grave Identifier:** UE 10829; T-5; CA07. **Grave Type:** Collective pit grave. W-E orientación. **Skeletal information:** Male, Adult. **Grave goods:** Bronze liriform belt clasp with buckle and clasp pin in the pelvic area. **Dating:** Direct InCal 20 radiocarbon dated to 607-774 calCE (1360±30 BP, Beta-471267). **Ancestry summary:** Iberia_IA__+__WestAnatolia_Roman_Byzantine. **Additional information:** Primary position in the pit grave.

- **Genetic Identifier:** I15423. **Grave Identifier:** UE 10718; T-122; CA10. **Grave Type:** Collective pit grave. **Skeletal information:** undetermined sex, jubenile. **Grave goods:** A cylindrical bronze washer, several fragments in the shape of earrings and a circular appliqué of the same material. **Dating:** Archeologically dated to 600-700 CE. **Ancestry summary:** Iberia_IA__+__WestAnatolia_Roman_Byzantine__+__North_African_Punic. **Additional information:** Raided tomb.

- **Genetic Identifier:** I15424. **Grave Identifier:** UE 10652; T-154; CA2010; maxila 3. **Grave Type:** Collective pit grave**. Skeletal information:** Undetermiend sex and age. **Grave goods:** No grave goods. **Dating:** Archeologically dated to 600-700 CE. **Ancestry summary:** Excluded from the genome wide analysis due to low coverage. **Additional information:** Yaw sampled from a pile of bones in the pit.

- **Genetic Identifier:** I15426. **Grave Identifier:** UE 10652; T-154; CA2010; maxila 1. **Type:** Collective pit grave**. Skeletal information:** Undetermiend sex and age. **Grave goods:** No grave goods. **Dating:** Archeologically dated to 600-700 CE. **Ancestry summary:** Excluded from the genome wide analysis due to low coverage. **Additional information:** Yaw sampled from a pile of bones in the pit.

- **Genetic Identifier:** I15427. **Grave Identifier:** UE 10651; T-154; CA2010. **Grave Type:** Collective pit grave**. Skeletal information:** Male, Adult, between 25 and 35 years old. **Grave goods:** No grave goods. **Dating:** Direct InCal 20 radiocarbon dated to 564-650 calCE (1460±30 BP, Beta-421969). **Ancestry summary:** Iberia_IA__+__North_African_Punic. **Additional information:** Dental calculus.

- **Genetic Identifier:** I15428. **Grave Identifier:** UE 10650; T-154; CA10. **Grave Type:** Collective pit grave**. Skeletal information:** Undetermiend sex, adult, between 25 and 35 years old.. **Grave goods:** a rectangular iron belt buckle -without needle- of iron. **Dating:** Direct InCal 20 radiocarbon dated to 597-664 calCE (1410±30 BP, Beta-421970). **Ancestry summary:** Excluded from the genome wide analysis due to low coverage. **Additional information:** Presented a slight bulge in the medial aspect of the right tibia, approximately in the middle area of the diaphysis. This may be the result of a periostitis due to a small traumatism.

#### 1.31 Zaragoza, Las Lomas

Contact Person(s): Isidro Aguilera

**Brief settlement history:** This site has a long history of human occupation. It began in the Bronze Age (around 1900-1700 BC) with outdoor settlements containing trash pits, one of which was used as a burial tomb. During the Roman Empire (1st-3rd centuries AD), it functioned as a rural agricultural settlement with two cisterns made of hydraulic mortar. In the Visigothic period (5th to early 8th centuries AD), it became a necropolis with over 450 Christian burial tombs, including monolithic sarcophagi made of alabaster or sandstone, tiled graves, and simple pits. Nearby was a Visigothic village with silos for domestic waste. In the early Islamic period (8th century AD), pottery vessels were found in some of these silos.

**Excavation History:** Discovered in 1993 during preparatory work for the Zaragoza-Madrid high-speed railway, previously unknown, excavations took place in 1994 and 1999 by Arqueo-Expert S.L. Limited to the railway construction area, the site's actual size exceeds the excavated area.

**Geographic Information:** Located in Epila municipality on a river terrace of the left bank of the Jalón River, a tributary of the Ebro River to the south. The terrace consists of sand, gravel, and rounded stones, historically used for dry farming.

**Summary of sampled materials:** During the rescue excavations 15 individuals were unearthed. These individuals were stored at the Zaragoza archeological museum storage facilities. Unfortunately, due to a loss in excavation information, little is known about the archeological context of each individual.

**Graves:**

- **Genetic Identifier:** I30586. **Grave Identifier:** Tumba 3A.85, caja 2050. **Grave Type:** untraceable. **Skeletal information:** untraceable. **Grave goods:** untraceable. **Dating:** Archeologically dated to 400-800 CE. **Ancestry summary:** Iberia_IA__+__WestAnatolia_Roman_Byzantine__+__North_African_Punic.

- **Genetic Identifier:** I30587. **Grave Identifier:** Tumba 3b.86, caja 2050. **Grave Type:** untraceable. **Skeletal information:** untraceable. **Grave goods:** untraceable. **Dating:** Archeologically dated to 400-800 CE. **Ancestry summary:** Iberia_IA__+__WestAnatolia_Roman_Byzantine__+__North_African_Punic.

- **Genetic Identifier:** I30588. **Grave Identifier:** Tumba 12, caja 2037. **Grave Type:** untraceable. **Skeletal information:** untraceable. **Grave goods:** untraceable. **Dating:** Archeologically dated to 400-800 CE. **Ancestry summary:** Excluded from the genome wide analysis due to low coverage.

- **Genetic Identifier:** I30589. **Grave Identifier:** Tumba 30, caja 2039. **Grave Type:** untraceable. **Skeletal information:** untraceable. **Grave goods:** untraceable. **Dating:** Archeologically dated to 400-800 CE. **Ancestry summary:** Iberia_IA__+__WestAnatolia_Roman_Byzantine__+__North_African_Punic.

- **Genetic Identifier:** I30590. **Grave Identifier:** Tumba 32, caja 2034. **Grave Type:** untraceable. **Skeletal information:** untraceable. **Grave goods:** untraceable. **Dating:** Archeologically dated to 400-800 CE. **Ancestry summary:** Iberia_IA__+__WestAnatolia_Roman_Byzantine__+__North_African_Punic

- **Genetic Identifier:** I30591. **Grave Identifier:** Tumba 35. **Grave Type:** untraceable. **Skeletal information:** untraceable. **Grave goods:** untraceable. **Dating:** Archeologically dated to 400-800 CE. **Ancestry summary:** Iberia_IA__+__CNE_EarlyMedieval__+__WestAnatolia_Roman_Byzantine__+__North_ African_Punic

- **Genetic Identifier:** I30592. **Grave Identifier:** Tumba 65A, caja 2052. **Grave Type:** untraceable. **Skeletal information:** untraceable. **Grave goods:** untraceable. **Dating:** Archeologically dated to 400-800 CE. **Ancestry summary:** Iberia_IA__+__CNE_EarlyMedieval__+__WestAnatolia_Roman_Byzantine__+__North_ African_Punic.

- **Genetic Identifier:** I30597. **Grave Identifier:** Tumba 184. **Grave Type:** untraceable. **Skeletal information:** untraceable. **Grave goods:** untraceable. **Dating:** Archeologically dated to 400-800 CE. **Ancestry summary:** Italy_Sicily_Punic_Roman__+__North_African_Punic.

- **Genetic Identifier:** I30598. **Grave Identifier:** Tumba 188, caja 2041. **Grave Type:** untraceable. **Skeletal information:** untraceable. **Grave goods:** untraceable. **Dating:** Archeologically dated to 400-800 CE. **Ancestry summary:** Iberia_IA__+__WestAnatolia_Roman_Byzantine__+__North_African_Punic.

- **Genetic Identifier:** I30599. **Grave Identifier:** Tumba 201.99, caja 2042. **Grave Type:** untraceable. **Skeletal information:** untraceable. **Grave goods:** untraceable. **Dating:** Archeologically dated to 400-800 CE. **Ancestry summary:** Iberia_IA__+__CNE_Early_medieval__+__North_African_Punic.

- **Genetic Identifier:** I30930. **Grave Identifier:** Tumba 8, caja 2036. **Grave Type:** untraceable. **Skeletal information:** untraceable. **Grave goods:** untraceable. **Dating:** Archeologically dated to 400-800 CE. **Ancestry summary:** Iberia_IA__+__CNE_EarlyMedieval__+__WestAnatolia_Roman_Byzantine__+__North_ African_Punic

- **Genetic Identifier:** I30931. **Grave Identifier:** Tumba 18. **Grave Type:** untraceable. **Skeletal information:** untraceable. **Grave goods:** untraceable. **Dating:** Archeologically dated to 400-800 CE. **Ancestry summary:** Iberia_IA__+__WestAnatolia_Roman_Byzantine__+__North_African_Punic

- **Genetic Identifier:** I30932. **Grave Identifier:** Tumba 21a. **Grave Type:** untraceable. **Skeletal information:** untraceable. **Grave goods:** untraceable. **Dating:** Archeologically dated to 400-800 CE. **Ancestry summary:** Iberia_IA__+__WestAnatolia_Roman_Byzantine__+__North_African_Punic.

- **Genetic Identifier:** I30933. **Grave Identifier:** Tumba 34B, caja 2045. **Grave Type:** untraceable. **Skeletal information:** untraceable. **Grave goods:** untraceable. **Dating:** Archeologically dated to 400-800 CE. **Ancestry summary:** Iberia_IA__+__WestAnatolia_Roman_Byzantine.

- **Genetic Identifier:** I30935. **Grave Identifier:** Tumba 204a, caja 2047. **Grave Type:** untraceable. **Skeletal information:** untraceable. **Grave goods:** untraceable. **Dating:** Archeologically dated to 400-800 CE. **Ancestry summary:** Excluded from the genome wide analysis due to low coverage.

#### 2 Direct AMS 14C Dates

Radiocarbon dating was performed for range of ancient samples following standard procedure (Olalde et al., 2019), at the Pennsylvania State accelerator mass spectrometry radiocarbon laboratory. The dated material (bone or tooth) was the same as the one subjected to ancient DNA analysis. Dates were calibrated to 2 sigma using OxCal v4.4.2 and the IntCal20 calibration curve (Reimer et al., 2020). We also obtained δ13C and δ15N values, informing about dietary habits.

.

#### 3 Ancient DNA laboratory procedures

We conducted aDNA extraction on a total of 255 ancient individuals from the Iberian Peninsula (Present day Spain).

Our laboratory procedures were carried out in specially designated clean rooms. To minimize the risk of exogenous DNA contamination, we removed the outermost layer of teeth and long bones. Powder collected from beneath the cleaned area was obtained through low-speed drilling to prevent DNA damage from heat (Adler et al., 2011). Cochleae were extracted from the temporal bone through sandblasting and subsequently milled (Pinhasi et al., 2019). The resulting powder underwent lysis buffer incubation. DNA was then cleaned and concentrated using silica magnetic beads (Rohland et al., 2018) in conjunction with a manual or automated protocol, employing Dabney Binding Buffer for manual extraction (Dabney et al., 2013; Korlević et al., 2015).

Barcoded double-stranded libraries, utilizing truncated adapters from the extract (equivalent to 6.2 to 8.4 mg of powder), were prepared. Prior to blunt-end repair, libraries underwent partial uracil–DNA–glycosylase (UDG) treatment to significantly reduce the characteristic damage pattern of aDNA (Briggs et al., 2009; Rohland et al., 2015).

For the enrichment of human DNA, we employed probes targeting either 1,233,013 SNPs ('1240k capture') (Fu et al., 2015) or 1,352,535 SNPs ('Twist' BioSciences) (Rohland et al., 2022), as well as the mitochondrial genome. The '1240k' reagent underwent two rounds of capture, while the 'Twist' BioSciences reagent underwent one. Subsequently, the captured libraries were sequenced on an Illumina HiSeq X10 instrument with 2x101 cycles and 2x7 cycles for reading out the two indices (Kircher et al., 2012), or on an Illumina NextSeq 500 instrument with 2x76 cycles and 2x7 cycles for reading out the two indices.

### 4 Bioinfomatic processing

Raw sequence data reads for each sample were identified and isolated based on sample-specific indices introduced during wet-lab processing, allowing for one mismatch. Adapters were trimmed, and paired-end sequences were merged into single-ended sequences, requiring a 15-base pair overlap (with allowance for one mismatch). This process was performed using a modified version of *SeqPrep 1.1* (https://github.com/jstjohn/SeqPrep), which selected the highest quality base in the merged region. Unmerged reads were discarded prior to alignment against both the human reference genome (hg19) and the RSRS version of the mitochondrial genome, employing 'samse' command in *bwa* (version 0.6.1) (Li & Durbin, 2009). Duplicates were removed based on the alignment coordinates and orientation of aligned reads. Libraries were sequenced to saturation across multiple lanes when necessary, and complexity metrics were determined using *preseq* (Daley & Smith, 2013), with merging performed as needed. The computational pipelines are accessible on GitHub (https://github.com/dReichLab/ADNA-Tools, https://github.com/dReichLab/adna-workflow).

Subsequent validation of ancient DNA authenticity involved several criteria. Libraries with a deamination rate at the terminal nucleotide below 3% were excluded from further analysis. The ratio of X-to-Y chromosome reads, mismatch rates to the consensus mitochondrial sequence (using *contamMix-1.0.10*) (Fu et al., 2013), and X-chromosome contamination estimates (using *ANGSD*) (Korneliussen et al., 2014) in males with sufficient coverage were computed. Libraries showing evidence of contamination or lacking a minimum of 15,000 SNPs with at least one overlapping sequence were excluded from genome-wide analyses.

All analyses in this study were confined to the 1,233,013 SNPs common to both 1240k and Twist reagents, as well as the mitochondrial genome.

#### 5 Mitochondrial and Y-chromosome haplogroup determination

For the determination of mtDNA haplogroups in our samples, reads mapped to the mitochondrial reference genome were utilized, considering sequences with a mapping quality (MAPQ) of ≥ 30 and a base quality of ≥ 30. A consensus sequence was derived using *bcftools* and SAMTools (Li et al., 2009) through a majority rule, requiring a minimum coverage of two. Haplogroups were then assigned using HaploGrep3 based on Phylotree (mtDNA tree build 17, Forensic Update 1.2) (Schönherr et al., 2023) .

To ascertain the Y-chromosome lineages of ancient male individuals, we annotated the path of derived mutations following the nomenclature of Yfull 8.09 (https://www.yfull.com/), following the procedure outlined in Lazaridis et al. (2022). Additionally, we annotated the haplogroup name associated with the most derived mutation for each sample, using the nomenclature of the International Society of Genetic Genealogy (http://www.isogg.org; version 15.73). Manual curation of low coverage individuals was performed by visualization of alignments to haplogroup determining positions via Integrated Genome Viewer (IGV).

#### 6 Kinship analysis and Identification of Identity by descent (IBD) DNA fragments

We conducted an analysis to identify kinship relationships among the newly reported individuals in our study, employing two methods, the Relationship Estimation from Ancient DNA (READv2) (Alaçamlı et al., 2024) and KIN (Popli et al., 2023)

READv2 has the capability to infer family relationships up to the second degree, even in samples with very low coverage. To ensure independence of kinship classification from within-population diversity, we normalized the proportion of non-matching alleles (P0) before assessing relationships between pairs of individuals with default options for both window size and median pairwise P0. Pairwise comparisons were performed among all of the newly reported individuals of the dataset. To minimize the impact of the high heterogeneity of ancestral backgrounds of the samples, we also performed a READv2 run for individuals within archeological sites, and also among individuals within the same broad period (Roman and post-Roman control period), together with a geographic divide (Northern and Southern Iberia) (Table S3).

Kin is a method that estimates the relatedness of a pair of individuals from shared identical-by-descent fragments based on a hidden Markov model (HMM) approach. This method also predicts the nature of the kinship relationship within the degree of relatedness. Pairwise comparisons were performed among all of the newly reported individuals of the dataset.

We also used this method to determine the nature of the first-degree relations in the dataset, most importantly the Vilanova-Blanes first-degree pair of individuals (Table S4). Kin results show that these two individuals (a woman from Vilanova d'Alcolea and a man from Blanes) were likely full siblings. To verify this result, we performed a run including the aforementioned pair together with individuals with similar ancestral origins (i.e. with substantial Germanic-relatic ancestry) from the Blanes necropolis, to avoid biases associated to ancestry heterogeneity. Furthermore, to test whether the amount of data overlap between these two individuals was enough to correctly determine the type of first-degree relation, we ran Kin on the individuals from the Hazleton North large pedigree, whose relationships are well-stablished (Fowler et al., 2022), downsampling at different coverage levels to match the coverage of the first-degree relatives from Vilanova-Blanes. We found that the determination of the type of first degree relationships was accurate at the coverage levels present in the Vilanova-Blanes pair (Fowler et al., 2022). The P0 value from the first run was used.

We calculated the shared IBD segments by imputing genome-wide data for our newly reported individuals according to the method described by (Ringbauer et al., 2017, 2024).

### 7 Aneuploidies and Runs of Homozygosity

We calculated possible aneuploidies by computing the mean coverage at the 1240k SNP array of each chromosome divided by the mean coverage for 1240k array recovered SNPs of all autosomes. Assuming a normal ploidy this ratio should be around 1 for the autosomes. For the sexual chromosomes: 1 for females and 0.5 for males at the X chromosome if no aneuploidies are present, and 0 for females and 0.5 for males if no aneuploidies are present. We identify no apparent aneuploidies in any of the newly reported individuals.

We assessed the possible presence of runs of homozygosity (ROHs) in the newly reported ancient individuals by applying the method described in (Ringbauer et al., 2021). We do not detect significant ROHs in any of the newly reported individuals of the dataset. The highest sum of >20cM fragments was 51.45 cM; and 98.93 cM when summing all the ROHs longer than 4cM, by individual I28354 from Cova Simanya.

#### 8 Principal Component Analysis

We conducted Principal Component Analysis (PCA) with the 'smartpca' tool in EIGENSOFT (v7.2.1). Ancient individuals were mapped onto present-day individuals' components using "lsqproject:YES" and "shrinkmode:YES" (Patterson et al., 2006; Price et al., 2006)

We computed 2 PCAs for the current analysis: The West Eurasian PCA, where PCs were computed on the HO dataset using 1036 present-day West Eurasian present-day individuals genotyped on the Human Origins array. This PCA shows the inferred affinities of our samples with European, Levantine and Middle East populations. The North-Africa West Eurasia PCA. Where PCs were computed using 1314 present-day West Eurasian and North African present-day individuals genotyped on the Human Origins array (Patterson et al., 2012).

### 9 Ancestry modelling

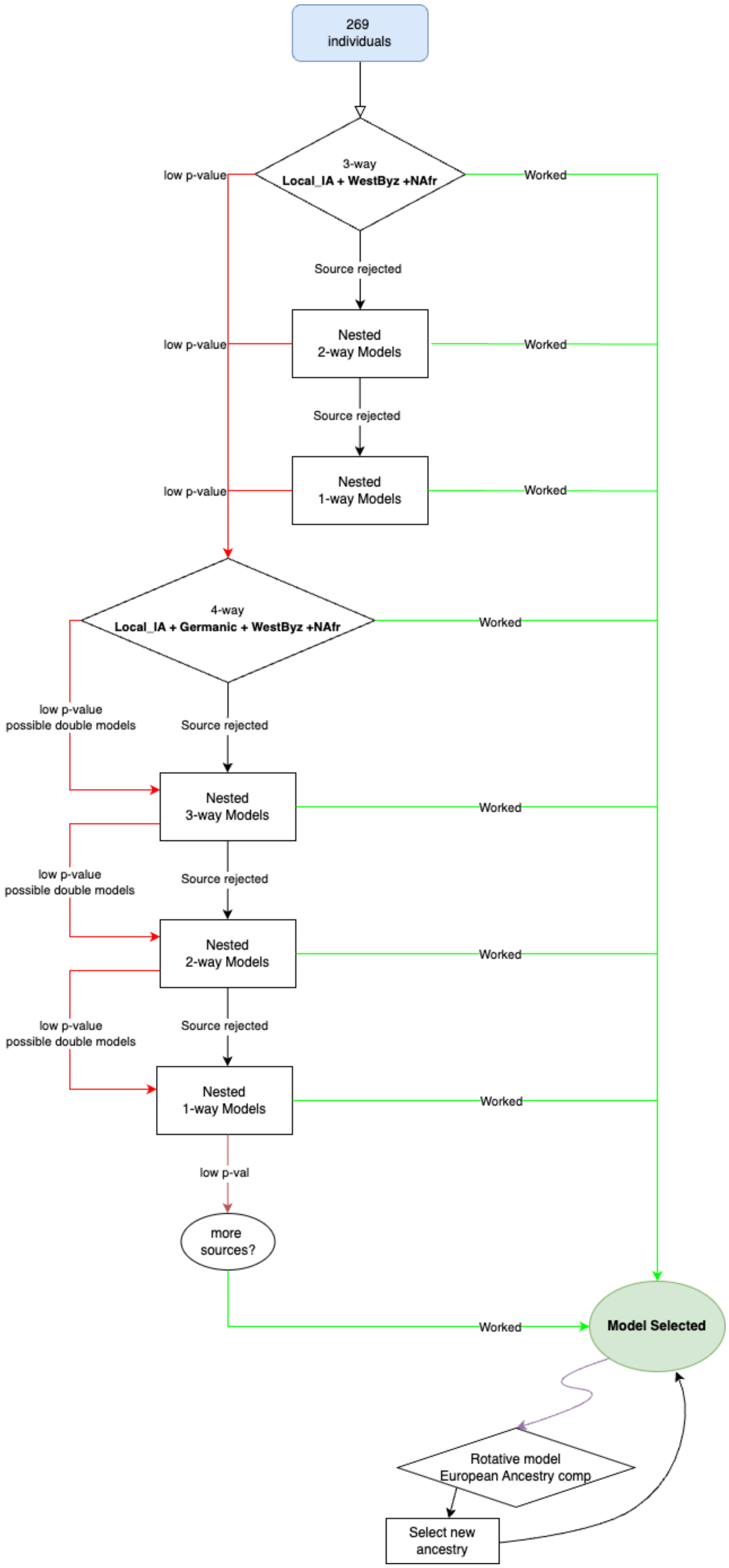

In this supplementary section we aim to understand the genetic ancestry underlying the Roman and Late Antiquity populations of the Iberian Peninsula. To accomplish this goal, we model the ancestry of the 271 newly reported individuals from this paper together with previously published 1240k captured individuals from the same region and time period (Olalde et al., 2019), using the f-statistics framework implemented in the *qpAdm* software from AdmixTools v.6.0 (https://github.com/dReichLab/AdmixTools).

The analyses described in this section were carried out in the dataset containing the 1,233,013 SNPs common to both 1240k and Twist reagents (see Supplementary section 4). Parameter "allsnps: YES" was set to use the maximum number of SNPs available for each $f_4$ statistic, and therefore minimizes the effects of missingness in our dataset. Also, to bypass the effect of allele capture bias observed when using together 1240k capture and shotgun data (Rohland et al., 2022) and its subsequent effects in f-statistics matrix calculations (Davidson et al., 2023), whenever possible we avoided the use of shotgun sequenced ancient individuals in any of our models, both as source populations and outgroups.

We are still far from fully understanding the demographic events that led to the complex demographic and linguistic landscape encountered by the Romans upon their arrival to Iberia in the 3rd century BCE (this will be a topic for future work). However, the broad ancestry make-up of Iberian Iron Age populations at the dawn of the proto-historic and historic periods has been investigated in previous work (Olalde et al., 2019) and we can use their genetic data as a proxy for the local Iberian component in individuals dated to later periods. Since our aim is to study the demographic impacts of the Roman colonization and the later Great Migration period, we develop a proximal ancestry model with Bronze Age populations preferentially as outgroups, which would be differentially related to the source populations (mainly Iron Age groups, e.g. Iberian Iron Age) contributing ancestry to our target individuals from the Roman and Late Antiquity periods dated ~0-700 CE.

We designed the following set of outgroups:

- Mota: We use the Mota cave individual (its 1240k data version), a 4500-year-old hunter-gatherer from Ethiopia without any evidence of West Eurasian admixture as the "base" outgroup (Gallego Llorente et al., 2015; Lipson et al., 2022).

- Morocco_Iberomaurusian: We use the individuals from the Iberomaurusian culture of North Africa (present-day Morrocco) as the outgroup giving leverage for teasing apart North African-related ancestry (Van de Loosdrecht et al., 2018).

- Spain_EBA_Argar: This population represents individuals from the Early Bronze El Argar culture in present-day Spain (Olalde et al., 2019; Villalba-Mouco et al., 2022). We include this outgroup for teasing apart Iberian-specific drift within Iron Age Iberian sources.

- Germany_EBA_Unetice: These individuals are associated with the Unetice culture of Early Bronze Age Germany (Mathieson et al., 2015).

- Turkey_Alalakh_MLBA: This population represents individuals from the Middle/Late Bronze Age site of Alalakh in Northern Levant (Skourtanioti et al., 2020). We use this outgroup for teasing apart Levantine-related ancestry.

- Steppe_IA: Individuals from the Steppe region during the Iron Age (Gnecchi-Ruscone et al., 2021; Jeong et al., 2020; Unterländer et al., 2017; Wang et al., 2021).

- Greece_Mediterranean_BA: This population represents individuals from the Bronze Age Aegean region (Lazaridis et al., 2022; Lazaridis & Reich, 2017; Skourtanioti et al., 2023). We use this outgroup to provide leverage for teasing apart ancestry from the Eastern Mediterranean-specific drift.

#### 9.1 Baseline model for the Roman and Late antiquity individuals from Iberia

The results of the West-Eurasian Principal Component Analysis suggest the presence of 3 main ancestral components in the majority of Roman and Late Antiquity individuals from Iberia (See Supplementary section 8., FIGURE 1C). These components are: ancestry derived from local Iberian Iron Age groups; ancestry derived from eastern Mediterranean populations, which was itself widespread during the Roman Imperial period in the city of Rome (Antonio et al., 2019), central Italy (Posth et al., 2021)and the Balkan peninsula (Olalde et al., 2023) and ancestry derived from North African populations. We made these assumptions based on:

- The PCA position of several Roman and Late Antiquity individuals overlapping or adjacent to the space occupied by previously published Iberian Iron Age individuals (Olalde et al., 2019; Patterson et al., 2022). A local Iberian component is also supported by the presence of uniparental markers with clear local Iberian origin, such as Y-chromosome lineage R1b-DF27, widespread in Iberia since the Early Bronze Age up until the present day (Olalde et al., 2019; Villalba-Mouco et al., 2022).

- The shift (higher values in PC1 of the West-Eurasian PCA) observed in many Roman and Late Antiquity individuals towards groups with high levels of Eastern Mediterranean ancestry such as Western Anatolian populations and the available population from Rome during the Imperial period the population. This is also supported by the presence of uniparental markers with a likely Eastern Mediterranean/Near Eastern proximal origin such as Y-chromosome haplogroups L1b, T1a or J2a, all absent in Iberia during earlier Bronze and Iron Age periods.

- The shift (higher values in PC1 and lower values in PC2 of the West-Eurasian PCA) observed in many Roman and Late Antiquity individuals towards ancient and present-day groups from North Africa. This shift is also clearly seen in the North-Africa West Eurasian PCA (Figure 1C). A North African component is further supported by the presence of uniparental markers with a likely North African proximal origin such as Y-chromosome haplogroups E1b-M81 and E1b-V65 and mtDNA haplogroup U6, as well as others with a more general African proximal origin such mtDNA haplogroups L1, L2 and L3. all absent in Iberia during earlier Bronze and Iron Age periods.

Based on these observations we modeled the ancestry profile of the newly reported and the previously published Roman and Late Antiquity individuals from Iberia using the following set of possible source populations:

- Iberia_IA: We use this population, which includes Iron Age individuals from the Iberian Peninsula (Olalde et al., 2019; Patterson et al., 2022), as a direct source of local ancestry. Out of the 22 individuals included in this group, 15 were excavated at non-Indo-European-speaking areas along the Mediterranean and only 7 were excavated at Indo-European-speaking areas at the northern edge of the Central Plateau. Given that these two areas were shown to carry different amounts of Central-European-related ancestry (higher in the latter), the bias towards non-Indo-European-speaking areas in the Iberia IA source could affect the estimation of local ancestry, specially in the regions where our test individuals from the Roman and Late Antiquity periods derive ancestry from local groups with high Central-European affinities.

- WestAnatolia_Roman_Byzantine: This population is composed by a group of individuals who inhabited the Aegean cost of Anatolia during the Roman-Byzantine period (Lazaridis et al., 2022). We use this source as a proxy for the Eastern Mediterranean signal present in the Italian Peninsula and the Balkans during the Imperial Period (Antonio et al., 2019; Olalde et al., 2023; Posth et al., 2021).

- North_Africa_Punic: This population is composed by ancient individuals dated to the Punic period from North Africa (Kerkouane, Tunisia) and South Spain (Villaricos, Almería), who are almost fully North African in ancestry (Ringbauer et al. in press). We consider this population would represent a relatively good proxy for the source of the North African signal observed in our test Iberian individuals from Roman and Late Antiquity periods. We caution, however, that North Africa during the second half of the 1st millennium BCE is currently severely undersampled, so the real North African source who contributed ancestry to our test individuals might differ from the North_Africa_Punic group we use as a proxy.

Due to the substantial ancestry heterogeneity observed in PCA, even among individuals from the same archaeological sites, we opted against aggregating the tested individuals into populations for this particular analysis. This decision was made to avoid losing interesting patterns that would otherwise go unnoticed when grouping several individuals. However, this approach diminishes the statistical power to reject poor-fitting models specially in low-coverage individuals, as compared to an approach where individuals are grouped into populations or clusters for modelling. As a result, caution needs to be taken when in the interpretating the results of low-coverage individuals.

For those individuals whose 3-way (*Iberia_IA* + *WestAnatolia_Roman_Byzantine* + *North_African_Punic*) model was rejected (p-value < 0.01) by the *qpAdm* algorithm, we developed a strategy to select a more suitable model. This entails examining whether there is any of the 3 components with a negative proportion (or high standard errors, which would exceed the value of the proportion). In these cases, we check the viability of the respective nested model without the rejected component/s. By iterating through this process, we aim to refine the model selection.

Out of the 271 individuals with a good-fitting model, 151 can be modeled as a mixture (with varying proportions of these 3 main components): *Iberia_IA, WestAnatolia_Roman_Byzantine, North_African_Punic.* A further 46 can be modeled as a two-way nested model: 31 as a Iberia_IA and *WestAnatolia_Roman_Byzantine*, 7 as *Iberia_IA* and *North_Africa_Punic,* and 8 as *WestAnatolia_Roman_Byzantine* with *North Africa Punic*. And a final number of 13 as a nested one-way model: 11 as *Iberia_IA*, 1 as *WestAnatolia_Roman_Byzantine*, and 1 as *North_African_Punic*. Among these individuals, we found 8 males carrying Y haplogroup DF27. However, there were 61 individuals remaining without any fitting model.

##### 9.2 Introduction of Central/Northern European-like ancestry

For those individuals who did not yield a good-fitting model with the approach explained in the previous subsection, 9.1, we hypothesized that for some of them, an additional ancestry component was missing in the initial model. Based on historical and archeological evidence, a good candidate is Central/Northern European-related ancestry brought by groups commonly known as the Vandals, Suevians, Alans and most prominently the Visigoths who migrated to the Mediterranean region during and after the decline and fall of the Roman Empire, reaching the Iberian Peninsula in the 5th century CE. Several archaeogenetic studies have identified and genetically characterized these groups, which had varying degrees of demographic impact depending on the region (Amorim et al., 2018; Antonio et al., 2019; Gretzinger et al., 2022; McColl et al., 2024; Olalde et al., 2019, 2023). The presence of Central/Northern European-related ancestry unaccounted for in our initial model is further supported by 1) the PCA position of several of our test individuals (mostly dated to the Late Antiquity), close or within the space occupied by Central/Northern European populations, including Germanic groups from the Migration Period such as the Langobards; and 2) the detection of I1 and R1a Y-chromosomes lineages of likely Central/Northern European origin in four Late Antiquity individuals in our transect.

To account for the incorporation of this ancestral component we added to the first baseline 3-way model from section 9.1 a new source population:

- CNE_EarlyMedieval: Including individuals from Northern Italy and the Carpathian basin excavated at Langobard-associated necropolises dated to the V-VI centuries CE (Amorim et al., 2018) These individuals present Central-Northern European ancestry and are contemporaneous to the sites in our transect with more PCA evidence of Central/Northern European ancestry, such as Pla de l'Horta and Blanes. We use them as a proxy for the groups arriving to the Iberian Peninsula during the Migration Period.

We, therefore, attempted to model with this 4-way model all of the individuals who did not yield a fitting model from section 9.1. We follow the same iterative approach defined in the previous section, i.e. if any of the ancestry proportions was negative, we looked for the nested model without the source(s) with negative proportion.

Following this approach, we found 51 individuals who carried a Germanic-like ancestry. 17 individuals can be modeled with a 4-way model with *Iberia_IA, WestAnatolia_Roman_Byzantine, North_African_Punic* with *CNE_EarlyMedieval* as left populations. The remaining 35 individuals had a fitting nested model: 10 as *CNE_EarlyMedieval + WestAnatolia_Roman_Byzantine + North_African_Punic*, 4 as *Iberia_IA + CNE_EarlyMedieval + WestAnatolia_Roman_Byzantine,* 4 as *Iberia_IA + CNE_EarlyMedieval +North_African_Punic*, 9 as two-way model *CNE_EarlyMedieval + WestAnatolia_Roman_Byzantine,* 4 as *CNE_EarlyMedieval + North_African_Punic and* individual I19895 from the Boadilla archeological site as a two-way admixture of *Iberia_IA + CNE_EarlyMedieval.* Finally*,* 2 individuals, a woman from Blanes (I23772) and a man (I28358) from Vilardida could only be modeled as 1-way *CNE_EarlyMedieval*.

##### 9.3 Non-fitting Individuals

After the previous steps, 10 individuals remained without a good-fitting (P-value>X) model. For these, we explored the fitting nested models of any of them starting with the basal 3-way model described in section 9.1 and then the nested models of the model proposed in section 9.2 if they still have no fitting model.

This could be explained by outlier individuals with ancestry components that are completely missing (e.g. East Asian ancestry) among our four source populations utilized so far, Alternatively, the poor fit could be the result of using source populations that, although generally similar to the actual populations contributing ancestry to our test individuals, present small differences in ancestry that drive the model failure. One could hypothesize, for instance, that people arriving to the Iberian Peninsula during the Roman colonization were not ancestrally homogenous, and that we are not fully capturing this diversity with our modelling framework and the limited array of previously published ancient populations from the relevant periods and regions that are available to us. Another equally plausible scenario is that North African populations contributing ancestry to Roman and Late Antiquity Iberia were ancestrally diverse, and the extremely limited available archaeogenetic data from this region does not fully caver this diversity.

For special outliers, we introduced extra ancestral components:

- Kazakhstan_Sarmatian_IA: A group of individuals from present-day East Kazakhstan Iron Age associated with the Sarmatian culture (Gnecchi-Ruscone et al., 2021). We use this population as a proxy the ancestry in Iron Age Steppe populations, who have also been shown to impact demographically in Great Migration-period populations in Central and Southeastern Europe (Olalde et al., 2023).

- Italy_Sicily_Punic_Roman: Individuals from Sicily during the Punic period (Rinbauer et al. in press). We use this population as a proxy for groups with a proximal origin in the central Mediterranean that could be admixing with Iberian populations but without the strong Eastern Mediterranean/Near Eastern signal that we model with WestAnatolia_Roman_Byzantine.

- Mongolia_IA, contaminated?

Four individuals (3 from which Lucena's Coracho Basilica Lucena archeological site) could be modeled as *Italy_Sicily_Punic_Roman* with *North_Africa_Punic*-related ancestry. This indicates, perhaps, a different source of Mediterranean ancestry that arrived in the Iberian Peninsula during late Roman rule, whose origin would probably have a more central Mediterranean origin.

Two individuals needed a Steppe Iron Age ancestry source for the model to fit, both dated to the Late Antiquity and excavated at the Blanes archeological site. We know that the peoples who arrived in the peninsula during the late antiquity had had contacts with nomad steppe tribes. For example, the goths settled for several centuries on the Black Sea northern shores, close to the Pontic Steppe.

##### 9.4 West European ancestry source in lower coverage samples

Due to the recent history of gene flow between Iberia and Central Europe during the Bronze Age and Iron Age periods (Olalde et al., 2018, 2019; Villalba-Mouco et al., 2022), we wondered whether our qpAdm setup was able to properly tell apart the proxy for Central/Northern European ancestry arriving to Iberia during Late Antiquity *(CNE_EarlyMedieval)* from the local Iron Age proxy (*Iberia_IA).* In principle, we would expect more issues in low coverage individuals or in cases where the amount of *Iberia_IA/CNE_EarlyMedieval*-related ancestry to be modelled was small.

To assess the extent of this possible mischaracterization of *Iberia_IA and CNE_EarlyMedieval*-related ancestries, we tried to model each of the individuals with either *Iberia_IA* or *CNE_EarlyMedieval* ancestry in their model but exchanging the European source by the other one. We observe that 40% of the individuals (71 out of 222) have a good-fitting model regardless of which of the two sources is used. In other words, the model fits well with both *Iberia_IA* or *CNE_EarlyMedieval* as the source in models like (*Iberia_IA* or *CNE_EarlyMedieval) + WestAnatolia_Roman_Byzantine + North_African_Punic*. We therefore consider that the approach in section 9.1 could be underestimating the proportion of Central-North European ancestry in some test individuals, as this ancestry will be misattributed to the *Iberia_IA* source, resulting in a good fit for the model, which in turn will result in these individuals not progessing to the step in section 9.2 where *CNE_EarlyMedieval* is specifically added as the fourth source.

To mitigate this issue, we design a rotative strategy to select the correct source. This strategy consists of placing either *Iberia_IA* or *CNE_EarlyMedieval* in the outgroups and using the other as a source. If, for instance, the model with *Iberia_IA* in the sources and *CNE_EarlyMedieval* in the outgroups yields a good fit, while the other model is rejected, then the ancestry in our test individual is more likely derived from *Iberia_IA*.

With this approach, we manage to correctly identify the ancestry source of 10% of the samples (5/71), where we can distinguish clearly the European source. 18 samples wouldn't have a fitting model when having one of the Iron Age / Late Antiquity European populations in the outgroups. The algorithm is unable to differentiate between the European ancestry component of the rest of the samples 66 samples. We consider this is due to two main reasons: The lower the coverage of a sample, the lower the statistical power for the model to reject a model with two very similar sources, and the proportion of European ancestry is too small to have any resolution for discriminating this component.

In addition, to further determine the affinity of the newly reported individuals to local Iron age Spain_IA sources in opposition to CNE_EarlyMedieval migration period proxy source, we also computed f4(Mbuti, newly_reported_individual_over250k_SNP; Iberia_IA, CNE_EarlyMedieval).

We observe that the affinities of the individuals correspond with the assigned European ancestry source with the exception of individuals I18399 who yields a significant amount of North African ancestry, and, thus, drawing affinity from Mbuti set as anchor outgroup. Individual I23276 shows more affinity for CNE_EarlyMedieval in these tests, although having Spain_IA assigned as its European ancestry source. However, we detect this could be a matter of yielding more steppe-like ancestry as shown by the inclusion of Kazakhstan_IA as an ancestry component in its model.

##### 9.5 Modern individuals Modelling IBS (Model and FDstat)

To analyze the long-term impact of the migration period on the Iberian demographics we designed an approach involving present-day Iberian populations from the 1000k Genomes project (Clarke et al., 2012). We attempted to model the individuals from this dataset with the same models described in sections 9.1 and 9.4. However, none of the models yielded statistical fit. To ensure these results were not driven by known biases when co-analyzing different data types: ancient data from the outgroups and sources derived from SNP capture while the present-day test populations derived from shotgun sequencing (Davidson et al., 2023; Rohland et al., 2022), we explored the behavior of the models when pruning out from the dataset the SNPs with the strongest bias and keeping 620,466 SNPs (Davidson et al., 2023; Rohland et al., 2022).

Still, we cannot find a fitting model for present-day Iberian populations, however, the p-values significantly increase when pruning the SNPs with the strongest bias. We tested if this increase is due to a lower number of SNPs in the dataset, decreasing the ability to reject non-fitting models. We did so by randomly pruning out the same number of random. In this case the p-values do not increase, indicating that the increase in p-values when pruning out SNPs with strong bias is not due to a decrease in resolution.

Yet, we can observe patterns when analyzing the results of this approach. The model that most closely approaches a good-fitting p-value is the 4-way of *Iberia_IA, WestAnatolia_Roman_Byzantine, North_African_Punic* with *CNE_EarlyMedieval.* We observe that the present-day Basques have the least amount of North African Ancestry. Whereas the highest is the present-day Galician, Andalusian, and Murcian populations. We also observe that the highest proportion of *WestAnatolia_Roman_Byzantine* is in the insular Balearic population, implying a higher impact of Mediterranean migration into the Balearic islands and therefore a differential demographic impact of Roman colonization.

Perhaps the non-fitting of this model is due to the demographic transformations that occurred in the Iberian peninsula during the Medieval period.

To further analyze the affinities of present-day Iberian populations we computed the following f-statistics tests:

- f3(IBS_region,African_HO_SG,Chimp): We performed this f3 test to detect the affinities of the present-day Iberian populations to a broad sample of African populations (north African and Sub-Saharan African).

- f4(IBS_region,Roman-Visigoth_metaPop; North_Africa_SGDP,Mbuti): We performed this f4 setup to identify if the population dynamics subsequent to the end of the Germanic rule in the Iberian Peninsula, further impacted its demographics. If we detect more affinity from present-day north African populations with the Roman and Visigothic period metapopulations, we would conclude that the North African signal in the peninsula was decreased during medieval times. However, we detect that present-day north African populations have more affinities (positive Z scores) for present-day Iberian populations.

- f4(Mbuti,IBS_region; Iberia_IA,CNE_EarlyMedieval): We computed this f4 setup to analyze if there was differential demographic impact from the migration period peoples in the local population, that endured during time. We detect no significant population affinities, but all of the Z-scores are displaced to Spain_IA further suggesting that the Germanic impact in the peninsula was limited.
